# *De novo* design of ligases for programmable site-specific protein ubiquitylation

**DOI:** 10.64898/2026.11.08.765184

**Authors:** Saman Salike, Maximilian Gantz, Maximilian Fottner, Timo Flügel, Stanislav S. Moroz, Andrew C. Hunt, Michael Groll, Donald Hilvert, Kathrin Lang, David Baker

**Affiliations:** Department of Biochemistry, Institute for Protein Design, University of Washington, Seattle, WA, 98195, USA; Institute for Protein Design, University of Washington, Seattle, WA, USA; Department of Chemical Engineering, University of Washington, Seattle, WA, USA; Max Planck Institute for Terrestrial Microbiology, Marburg, Germany; Laboratory of Organic Chemistry, Department of Chemistry and Applied Biosciences, ETH Zurich, Zurich, Switzerland; Graduate Program in Biological Physics, Structure and Design, University of Washington, Seattle, WA, USA; Technical University of Munich, School of Natural Sciences, Department of Bioscience, Center for Protein Assemblies, Garching, Germany; Howard Hughes Medical Institute, University of Washington, Seattle, WA, USA

**Author notes:** These authors are co-first authors with equal contributions (Saman Salike, Maximilian Gantz, Maximilian Fottner).

## Abstract

Protein ubiquitylation regulates numerous cellular processes, but difficulties in generating site-specifically ubiquitylated proteins limit our understanding of individual modification events. In cells, ubiquitylation is mediated by an E1-E2-E3 enzyme cascade, with substrate recognition largely determined by more than 600 human E3 ligases, making it challenging to redirect ubiquitin transfer to a chosen lysine. Here we use RFdiffusion3 to design ubiquitin ligases from scratch that use a cysteine-based catalytic triad to conjugate an activated ubiquitin donor to a selected lysine in a chosen target substrate. Of 48 tested designs, 35 ubiquitylated a model peptide, with the most active ligases achieving near-quantitative ubiquitin transfer. We next challenged the approach with the homotrimeric sliding clamp proliferating cell nuclear antigen (PCNA), whose ubiquitylation at lysine (K) 164 governs DNA damage tolerance while displaying 45 off-target lysines for ubiquitination. Screening of 48 designs delivered a site-selective ligase. Incorporation of a single activity-increasing substitution and coupling to E1-dependent donor activation resulted in 90% conversion from unactivated ubiquitin, delivering higher rates and superior selectivity compared to a natural promiscuous E2. Using this cascade, we transferred di-, tri- and tetraubiquitin chains of defined length to PCNA K164 through a native isopeptide bond. These results establish *de novo* enzyme design as a route towards programmable, site-specific protein ubiquitylation. That a single *de novo* designed protein can bind both an activated ubiquitin donor and a globular target, and use a proficient catalytic machinery to selectively transfer ubiquitin to one lysine among 45, illustrates the level of functional complexity now within reach of *de novo* enzyme design.

## Main

Post-translational modifications (PTMs) regulate protein localisation, interactions, stability and activity, allowing cells to dynamically control protein function with dysregulation of PTMs implicated in a wide range of diseases^1^. Among these modifications, ubiquitylation is unusual because the modification is itself a protein: the 76-residue ubiquitin is attached through an isopeptide bond between its C-terminus and a lysine side chain of the target (Fig. 1A). Ubiquitylation regulates processes such as DNA damage tolerance, receptor endocytosis, transcription and proteasomal protein degradation^2^. A recent proteome-wide map identified approximately 100,000 ubiquitylation sites, but the function of most remains uncharacterised^3^. In nature, ubiquitylation is mediated by an E1–E2–E3 cascade in which an E1 enzyme activates ubiquitin and transfers it to E2 enzyme, while E3 ligases recruit target proteins and promote ubiquitin transfer to a substrate lysine (Fig. 1B). The human genome encodes more than 600 E3 ligases - with most putative ubiquitylation sites lacking assigned E3 ligases^4^. Understanding the functions of individual ubiquitylation events therefore depends on the ability to generate proteins modified at defined lysine residues. Chemical and semisynthetic approaches have enabled the preparation of precisely defined ubiquitin conjugates, but often require elaborate synthesis and protein refolding^5^. Enzymatic and chemoenzymatic strategies have expanded access to modifications of folded proteins, but are frequently constrained by recognition sequences or engineered attachment sites^6,7^. Programmable ligases that directly transfer ubiquitin to a chosen native lysine would therefore provide a powerful means to investigate the functions of individual ubiquitylation events.

**Figure 1.**
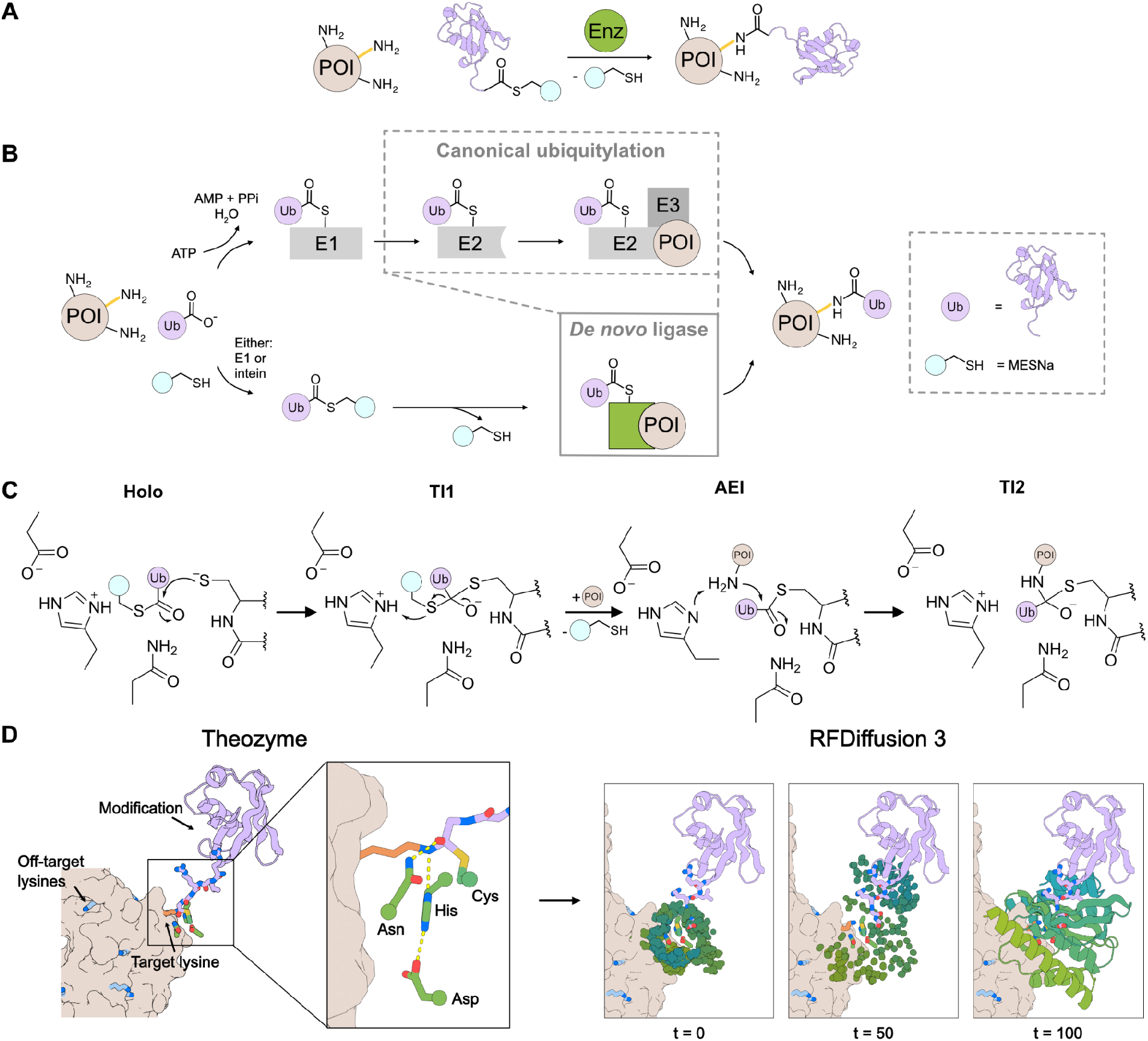
*De novo* design of ubiquitin ligases. **(A)** In this work we tackle the challenge to design enzymes for the selective ubiquitylation of lysines in a target protein via isopeptide bonds. **(B)** In the canonical E1-E2-E3 ubiquitylation cascade, ubiquitin is charged onto the active-site cysteine of an E1 enzyme, transferred to an E2 enzyme and site-specifically ligated onto a target lysine with the support of an E3 ligase. In this work, we design ubiquitin ligases that accept ubiquitin thioesters, charge them onto their own active-site cysteine and transfer ubiquitin site-specifically onto the target lysine. **(C)** A cysteine catalytic triad catalyses transfer of the substrate onto the active-site cysteine via a first tetrahedral intermediate (TI1) which collapses to form an acyl-enzyme intermediate (AEI). Attack of the incoming amine leads to formation of a second tetrahedral intermediate (TI2), followed by product formation. **(D)** The design pipeline involves generation of a theozyme in which the target lysine and ubiquitin are modelled into the active site of Ulp1 in the acyl-enzyme intermediate state. RFdiffusion3 is then used to generate ubiquitin ligases that both scaffold the active site in space and bind to the target protein.

We reasoned that *de novo* protein design could combine ubiquitin-transfer chemistry and target recognition within a single enzyme, enabling transfer from an activated ubiquitin donor to a specified lysine without requiring the natural E2–E3 machinery (Fig. 1 A,B), and thus providing a route to programmable ubiquitylation on demand. We set out to design ligases that incorporate cysteine-based catalytic triads, with the activated ubiquitin donor and target protein positioned to enable selective isopeptide bond formation at the desired lysine. By programming specificity at the design stage rather than attempting to remodel an existing protein, we reasoned that we could achieve high specificity for single lysines in both peptide and protein substrates.

### Design strategy for site-specific ubiquitin transfer

We adopted a two-step ping-pong mechanism in which the active-site cysteine first attacks the carbonyl carbon of an activated ubiquitin thioester, forming a tetrahedral intermediate (TI1) that collapses to generate a ubiquitin–enzyme thioester (acyl-enzyme intermediate, AEI). The ε-amine of the target lysine then attacks the AEI, forming a second tetrahedral intermediate (TI2) that resolves to the isopeptide-linked product and regenerates the catalytic cysteine (Fig. 1C).

To define the catalytic geometry, we modelled ubiquitin linked as a thioester to the active-site cysteine of Ulp1, using the structure of a covalently captured Ulp1–SUMO complex as a template^8^, and positioned the target lysine ε-amine for attack on the thioester carbonyl (Fig. 1D). We fixed the cysteine–histidine–aspartic acid catalytic triad and oxyanion-hole hydrogen-bond donors in space to create a minimal active-site model, or theozyme. The oxyanion hole comprised side-chain amide and backbone hydrogen-bond donors positioned to stabilise the negative charge on oxygen in TI1 and TI2. To bind and organise the flexible ubiquitin C-terminus, we included residues designed to form β-strand-pairing interactions with the acyl donor, building on previous protease designs^9,10^. For direct transfer to the selected lysine, we included the structural context of the target peptide/protein during the generative diffusion process (Fig. 2B and Fig. 3B).

**Figure 2.**
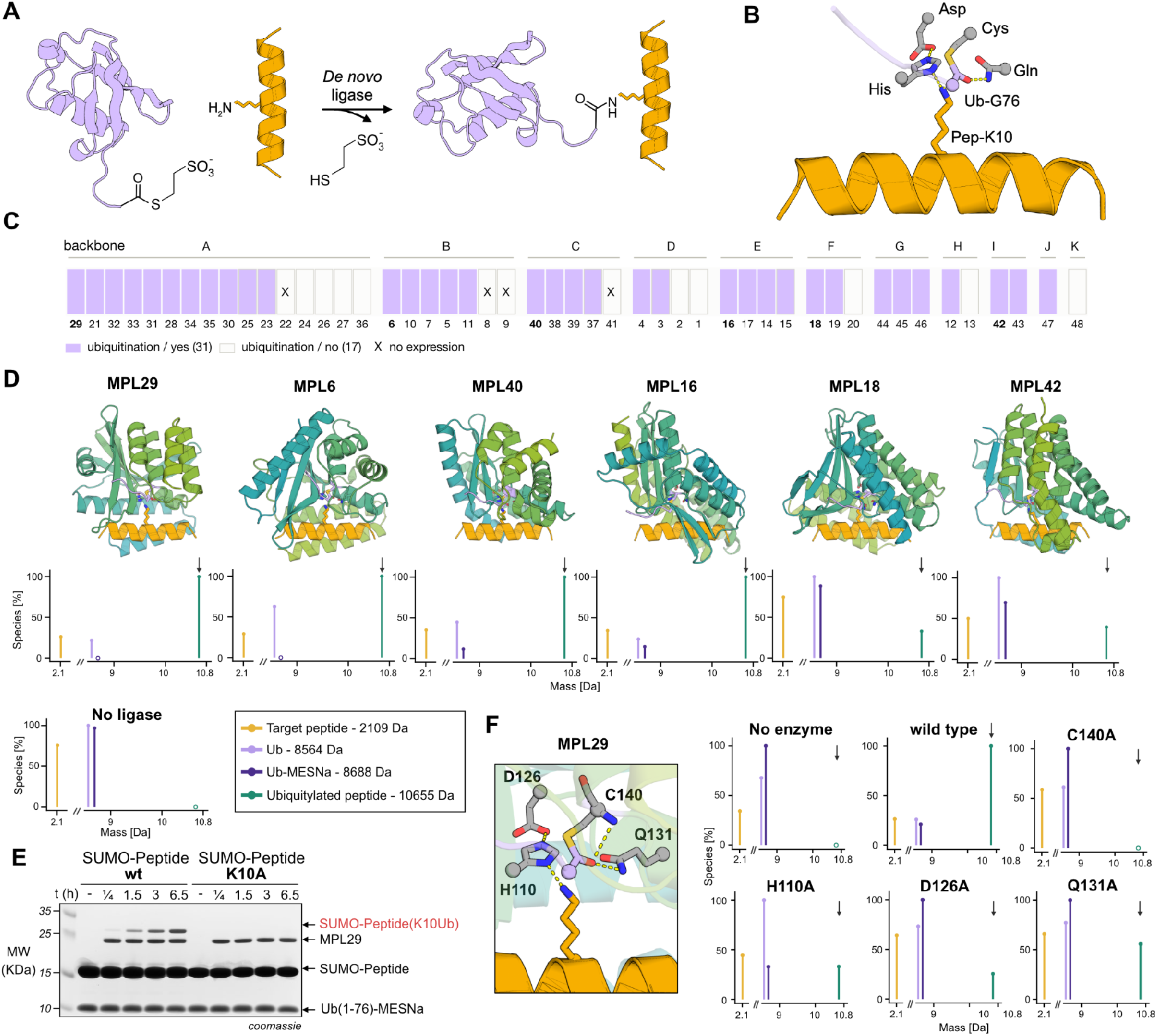
*De novo* design of a peptide ubiquitin ligase. **(A)** Reaction scheme. The *de novo* ligase transfers ubiquitin from Ub(1-76)-MESNa onto the lysine side chain of the target helix, forming an isopeptide bond and releasing MESNa. **(B)** Theozyme: the target peptide and ubiquitin modelled into the active site of Ulp1 in the AEI state. **(C)** Activity is distributed across most designed backbones (A–K). Non-expressing designs are highlighted with X. 35 of 48 tested designs show ubiquitylation activity (lavender). Reaction conditions: 125 μM Peptide, 50 μM Ub-MESNa, 50% (v/v) crude purified ligase in 10 mM Hepes pH 7.5, 150 mM NaCl at 37 °C for 20 h **(D)** Selected ligases with the highest ubiquitylation activity, assayed with 200 μM peptide, 100 μM Ub-MESNa and 20 μM ligase in 10 mM HEPES pH 7.5, 150 mM NaCl at 37 °C after 20 h. Ubiquitylated product is shown in green and highlighted with arrows, the target peptide in yellow, ubiquitin in light purple and ubiquitin thioester in purple. **(E)** Ubiquitylation of a SUMO–peptide fusion construct. Product was observed with the canonical peptide, whereas no ubiquitylation activity was detected upon mutation of the target lysine. Reaction conditions: 200 μM SUMO-Peptide, 100 μM Ub-MESNa, 20 μM Ligase 29 in 10 mM Hepes pH 7.5,150 mM NaCl at 37 °C. **(F)** Purified active-site mutants were assayed with 200 μM peptide, 100 μM Ub-MESNa and 20 μM ligase in 10 mM HEPES pH 7.5, 150 mM NaCl at 37 °C with 20 h of incubation. Ubiquitylated product is shown in green and highlighted with arrows, the target peptide in yellow, ubiquitin in pink and ubiquitin thioester in purple. No activity was observed for the active-site cysteine mutant, while residual activity remained for mutations of the other triad residues (H110 and D126) and of Q131.

**Figure 3.**
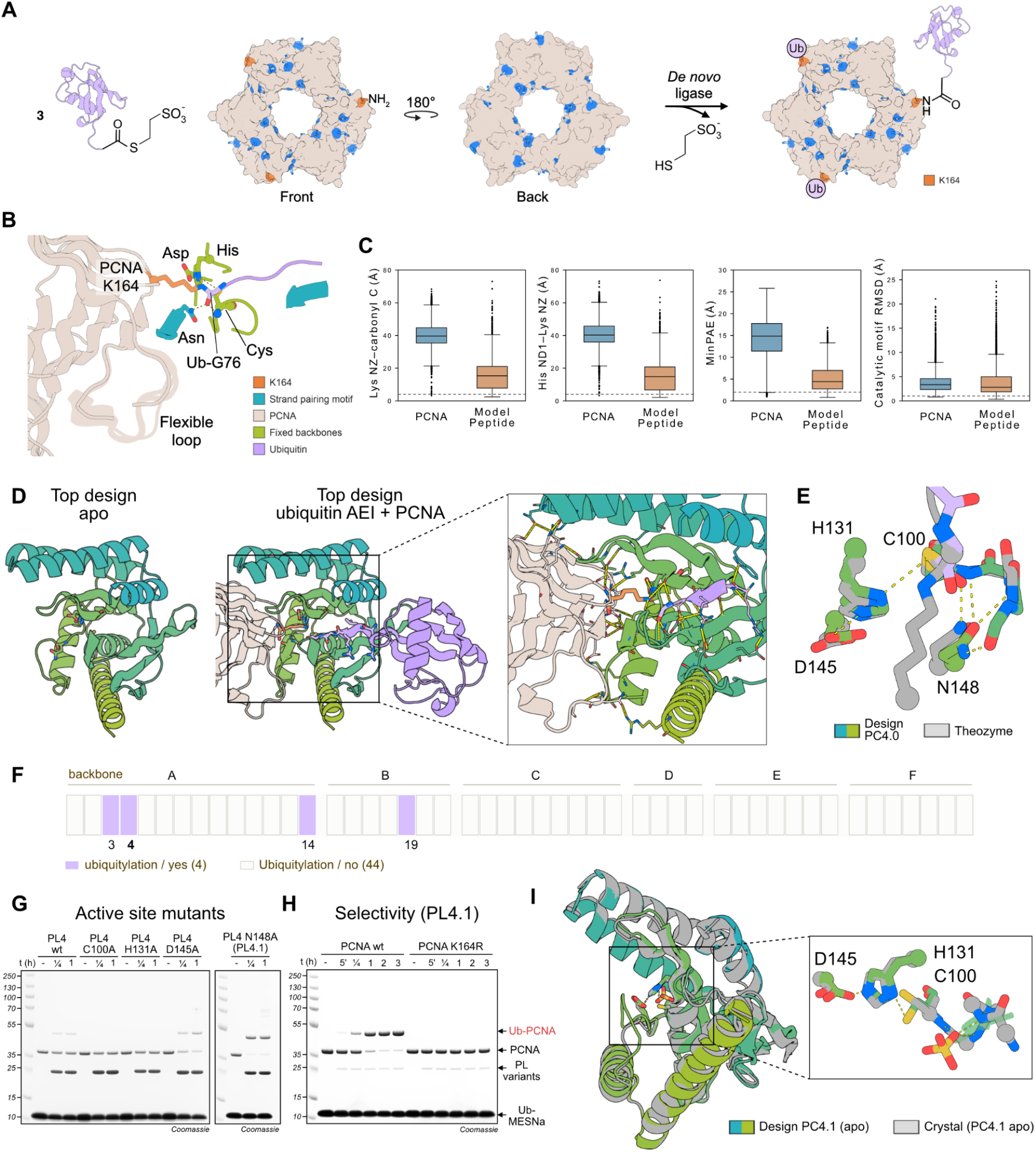
*De novo* design of a site-specific PCNA ubiquitin ligase. **(A)** Homotrimeric PCNA is targeted for ubiquitylation at its three K164 residues. **(B)** Theozyme: PCNA and ubiquitin modelled into the Ulp1 active site in the AEI state. **(C)** Benchmark of catalytic motif recovery and acceptor placement for PCNA (blue) and the model peptide (orange), using 900 backbones and ten LigandMPNN sequences per backbone. Box plots show, from left to right, the lysine Nζ–carbonyl carbon distance, histidine Nδ1–lysine Nζ distance, minimum predicted aligned error (minPAE) and catalytic motif root mean square deviation (RMSD). Dashed lines indicate design targets. **(D)** Design selection and predicted structures of PL4. Designs were filtered for active-site recovery in the apo state, accommodation of intact ubiquitin in the holo state, and confident substrate interfaces and productive lysine attack geometry in the AEI state. AF3 models show PL4 alone (left) and its ubiquitin AEI bound to PCNA (centre). The enlarged view (right) shows K164 positioned for attack on the thioester carbonyl carbon, with an Nζ–carbonyl carbon distance <3.5 Å and a Bürgi–Dunitz angle >90°. The PCNA strand beneath K164 and the ubiquitin C-terminus occupy the open groove and form the designed β-strand-pairing contacts. PL4 forms a shape complementary interface with several polar contacts to PCNA surface residues. PCNA, beige; ubiquitin, purple; K164, orange. **(E)** Theozyme overlaid with design 4 reveals that a slight miss-scaffolding of N148 results in it blocking the backbone hydrogen bond donor serving as oxyanion hole, flipping its amide bond so that it also cannot serve as hydrogen bond donor itself. **(F)** The 48 designed ligases were screened for transfer of Ub onto PCNA. A 2x master mix containing 200 μM Ub-MESNa and 100 μM PCNA was prepared in 20 mM HEPES pH 7.5, 150 mM NaCl and mixed 1:1 (v/v) with each Ni-NTA-purified, rebuffered ligase, giving final concentrations of 100 μM Ub-MESNa, 50 μM PCNA and 50 % (v/v) ligase eluate. Reactions were incubated at 37 °C for 20 h revealing four hits. PL4 is the most active and most specific ligase. **(G)** Active-site mutagenesis reveals a complete loss of activity upon mutation of the active-site cysteine, but increased ubiquitylation activity upon N148A and D145A mutation. Reaction conditions: 200 μM PCNA wt, 1800 μM Ub-MESNa, 200 μM PL4 variants in 10 mM Hepes pH 7.5 100 mM NaCl at 37 °C. **(H)** PL4.1 catalyses near-quantitative and selective ubiquitylation with 800 μM Ub-MESNa, 100 μM PCNA and 10 μM Ligase 4 N148A in 10 mM HEPES pH 7.5, 100 mM NaCl. **(I)** A crystal structure of PL4.1 apo at 1.6A shows 0.91 A Ca RMSD, 0.81 all atom RMSD with the catalytic triad with an AF3 prediction of PL4.1. The structure has a sulfate ion bound with an oxygen atom placed close to the expected position of the charged oxygen in the tetrahedral intermediate. This reveals that in the absence of N148, backbone atoms in this loop can serve as an oxyanion hole.

We used RFdiffusion3^11^ to generate protein scaffolds around the catalytic motif, the target recognition surface and the last five residues of ubiquitin (Fig. 1D). Starting from random coordinates, the model progressively denoised protein structures conditioned on these components to position the active-site residues and form contacts with both substrates. We designed sequences for the resulting backbones with LigandMPNN^12^ and used AlphaFold3 (AF3)^13^ to evaluate recovery of the designed catalytic geometry and binding to the desired substrates. We modelled the enzyme alone (apo) to assess recovery of the scaffolded active site, and the enzyme–ubiquitin complex (holo) to assess accommodation of intact ubiquitin without explicitly modelling the donor leaving group. We required that models of the ubiquitin–enzyme thioester bound to the target (AEI) had confidently predicted interfaces (interface minPAE < 2 Å) with the lysine ε-amine placed for nucleophilic attack on the thioester carbonyl carbon. Selected designs were subjected to further sequence optimisation with LigandMPNN and reevaluated with AF3, repeating this cycle to improve agreement with the target catalytic geometry and the position and orientation of the acceptor amine.

### *De novo* ubiquitin ligases for a model peptide

We first targeted a 19-residue model peptide containing a single lysine using ubiquitin activated as a MESNa thioester (Ub-MESNa) as the donor (Fig. 2A, Supplementary Fig. 1). To position both substrates for ligation, the design schema included the full peptide target, the catalytic residues, and strand-pairing contacts with the last five residues of the ubiquitin C-terminus (Fig. 2B). We generated 48 designs from 11 RFdiffusion 3 backbones, with multiple LigandMPNN sequences from most backbones passing the AF3 filters across the modelled reaction states. Of the 48 designs, 43 expressed well in *E. coli* and were purified for screening by liquid chromatography–mass spectrometry (LC–MS) (Supplementary Fig. 2). 35 designs showed measurable ubiquitylation activity, spanning 10 of the 11 backbones, with backbone K the sole exception (Fig. 2C). The designed catalytic arrangement thus supported ubiquitin transfer in a range of protein scaffolds.

Productive ligation requires transfer of ubiquitin to the target amine in competition with hydrolysis of the activated donor. Model peptide ligases (MPL) 16, 21, 29 and 40 gave near quantitative transfer of Ub-MESNa to the peptide after overnight incubation with 200 μM peptide, 100 μM Ub-MESNa and 20 μM ligase with only minimal hydrolysis of Ub-MESNa (Fig. 2D and Supplementary Fig. 3). More than half of the Ub-MESNa hydrolysed spontaneously over this period in the absence of enzyme, but most of the characterised designs did not increase hydrolysis above this background (Supplementary Fig. 3-6). The designed ligases therefore promote isopeptide bond formation without substantially accelerating donor hydrolysis in the presence of acceptor peptide.

We selected MPL29 for further characterisation. To determine whether recognition of the model peptide could select a single lysine in a larger protein context, we fused the peptide to SUMO, which presents nine additional surface-accessible lysines. Under optimised conditions, MPL29 delivered 64% conversion to ubiquitinated product after 3 h (Supplementary Fig. 7). Mutation of the target lysine to alanine (K10A) abolished detectable ubiquitylation, indicating that MPL29 is specific for the intended site in the presence of the nine alternative lysines (Fig. 2E, Supplementary Fig. 7). To determine whether ligation depended on the designed active site, we individually mutated the catalytic residues of MPL29. The C140A substitution abolished detectable activity, whereas H110A, D126A and Q131A retained some residual activity (Fig. 2F and Supplementary Fig. 8). These results support the importance of the designed catalytic motif for function.

### *De novo* ubiquitin ligases for PCNA

We next sought to extend site-specific ubiquitylation from a model peptide to a native protein. We selected proliferating cell nuclear antigen (PCNA), a homotrimeric sliding clamp that maintains DNA polymerase processivity during replication. Following DNA damage, monoubiquitylation of PCNA at K164 by Rad6-Rad18 recruits translesion synthesis polymerases, enabling lesion bypass and contributing to mutagenesis and chemoresistance in cancer (Fig. 3A)^14^. Preparation of site-specifically ubiquitylated PCNA for functional studies can require substantial experimental effort^15–19^. We therefore chose PCNA both as a stringent test of whether our approach could generalise from peptide recognition to direct recognition of a globular protein and because such ligases could facilitate downstream functional studies.

PCNA presented a substantially harder recognition problem than the model peptide, with three symmetry-related K164 sites and 45 other lysines in the trimer. The highly polar surface surrounding K164 lacks a clear hydrophobic patch for anchoring an enzyme, making it a challenging target for *de novo* interface design (Supplementary Fig. 9)^20,21^. An adjacent loop is partly unresolved in the crystal structure and adopts variable conformations across AF3 predictions, further complicating recognition (Fig. 3B). We reasoned that pairing with the β-strand beneath K164 could anchor the enzyme on PCNA and position the target lysine relative to the catalytic residues. We therefore incorporated a short strand designed to pair with PCNA into the motif, alongside the strand that binds the ubiquitin C-terminus (Fig. 3B). Scaffolding the resulting 14 fixed enzyme residues together with the five C-terminal residues of ubiquitin and the PCNA target tested the limits of current design methods. A benchmark comparing the model peptide and PCNA, using 900 backbones and ten LigandMPNN sequences per backbone, made this difference explicit (Fig. 3C). Although predicted structures recovered the enzyme motif, very few PCNA models positioned the lysine ε-amine for nucleophilic attack. Precise substrate placement across the enzyme–PCNA interface was therefore the limiting step, leaving too few suitable starting points for the iterative LigandMPNN/AF3 refinement used for the target helix.

To overcome this bottleneck, we used the designed enzyme–PCNA interaction as a template for guided RF3 predictions with the Catalytic Guidance Engine^22^. We progressively reduced classifier-free guidance from 0.75 to 0.5, 0.25 and finally 0 to optimise sequences for recovery of the desired geometry as the imposed interaction guidance was withdrawn. At each stage, we sampled new sequences with LigandMPNN and selected models that recovered both the AEI catalytic arrangement and the lysine attack geometry. The resulting designs underwent further local LigandMPNN sequence resampling and independent AF3 evaluation. We selected designs that recovered the designed active site in apo predictions, accommodated intact ubiquitin with its C-terminus positioned at the active site in holo predictions, and positioned the K164 ε-amine for nucleophilic attack on the thioester carbonyl in AEI predictions (Fig. 3D). We carried forward 48 designs from six backbones for experimental testing.

All 48 designs expressed solubly in *E. coli* (Supplementary Fig. 10). SDS–PAGE screening identified four designs capable of ubiquitylating PCNA (Fig. 3F). To assess site selectivity, we tested these designs against PCNA K164R. PCNA ligases (PL) PL3, PL14 and PL19 retained some off-target activity, whereas PL4 showed no detectable ubiquitylation of PCNA K164R, supporting selective modification of the intended lysine (Supplementary Fig. 11).

The PL4 design closely recapitulated the intended catalytic-triad architecture (Fig. 3E). However, in the AF3 prediction of the AEI, the asparagine side chain designed to contribute to oxyanion stabilisation is slightly mispositioned, blocking the intended backbone hydrogen-bond donor while its own amide is oriented away from the oxyanion. Thus, neither of the two designed oxyanion-hole interactions is formed.

To assess the contribution of the individual residues in the designed catalytic motif, we individually replaced the active site residues with alanine. Mutation of either C100 or H131 abolished detectable ubiquitylation, whereas D145A increased activity approximately threefold and mutation of the designed oxyanion-hole residue, N148A, increased activity approximately 26-fold (Fig. 3G and Supplementary Fig. 12). Thus, ubiquitin transfer depended on the designed cysteine and histidine, but removal of either the aspartate or asparagine side chain improved ligation. We therefore used PL4 N148A (PL4.1) for subsequent experiments. This variant converted 90% of PCNA to ubiquitylated product in 3 h with 800 μM Ub-MESNa, 100 μM PCNA and 10 μM ligase, while retaining selectivity for K164 (Fig. 3H).

To structurally characterise the design and investigate the origin of its improved activity, we determined the crystal structure of PL4.1 at 1.6 Å resolution. The structure closely matched the design with an overall Cα RMSD of 0.91 Å to the design and a 0.81 Å all atom RMSD with the catalytic triad (Fig. 3I). At the position vacated by N148, the structure coordinates a sulfate ion from the crystallisation solution, with one oxygen positioned 0.7 Å from the expected position of the oxyanion in the tetrahedral intermediate (Fig. 3I, Supplementary Fig. 9). The sulfate forms hydrogen bonds with R129 and the backbone amides of C100, G98 and Y99, indicating that this loop can provide an oxyanion-stabilising environment in the absence of the N148 side chain. Together with removal of the unintended interactions formed by N148 in the predicted PL4 structure, these additional backbone interactions may help explain the increased activity of PL4.1.

To enable ubiquitylation directly from unactivated ubiquitin, we used an E1 enzyme in the presence of ATP and MESNa (Fig. 4A, Supplementary Fig. 13). E1-mediated activation followed by transthioesterification with MESNa continuously generated Ub-MESNa *in situ* for transfer to PCNA by the designed ligase. Starting from unactivated ubiquitin, this cascade converted approximately 90% of PCNA to ubiquitylated product after 24 h with 200 μM ubiquitin, 100 μM PCNA and 10 μM ligase, achieving similar conversion to reactions using preactivated Ub-MESNa with 4-fold less ubiquitin (Fig. 4B). Thus, E1-dependent donor activation could be coupled directly to site-specific ubiquitin transfer by the designed ligase without requiring an E2 or native E3 ligase. For comparison, we tested Ubch5c S22R, an engineered promiscuous E2 that has been repurposed for E3-independent PCNA monoubiquitylation and represents an established enzymatic approach for preparing ubiquitylated PCNA *in vitro*^18^. Under similar E1-coupled conditions, Ubch5c modified PCNA more slowly than PL4.1 (60% vs 90% conversion after 24 h, respectively) and generated multiple ubiquitylated species, in contrast to the site-specificity of PL4.1 (Fig. 4C).

**Figure 4.**
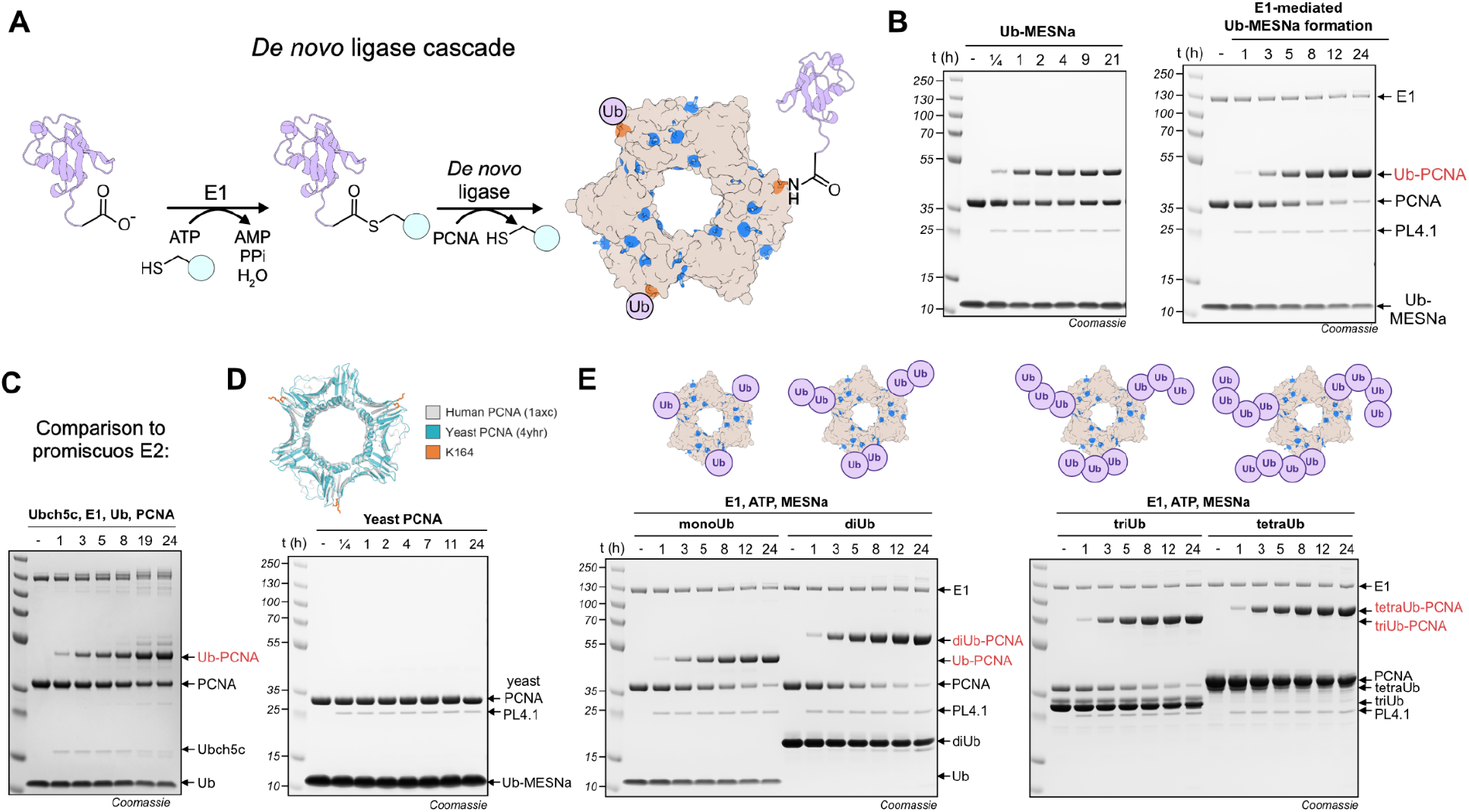
Quantitative site specific ubiquitylation with a designed ligase cascade. **(A)** A ubiquitylation cascade with E1 enzyme and MESNa enables *in situ* generation of the activated thioester, which is transferred onto PCNA by the designed ligases. **(B)** A *de novo* ligase cascade enables near-quantitative ubiquitylation of PCNA with 200 μM Ub-MESNa, 100 μM PCNA and 10 μM Ligase 4 N148A after 24 h of incubation. **(C)** The PL4.1 ligase cascade compares favourably with a promiscuous E2 ubiquitylation PCNA at K164 slower and with inferior selectivity with 100 μM PCNA wt, 200 μM Ub, 10 μM Ubch5c, 8 μM E1, 5 mM ATP, 6 mM MgCl_2_, in 10 mM Hepes pH 7.5, 100 mM NaCl at 37 °C **(D)** Despite a Cα RMSD of 1.3 Å, 100 μM yeast PCNA wt, 800 μM Ub-MESNa, 10 μM Ligase 4 N148A in 10 M Hepes pH 7.5 100 mM NaCl at 37 °C showed no detectable ubiquitylation. **(E)** Generation of defined, native polyubiquitylated PCNA using the *de novo* ligase cascade. Preformed linear (M1-linked) di-, tri- and tetra-ubiquitin was charged onto K164 using 100 μM PCNA WT, 200 μM mono/di/tri/tetra Ub WT, 10 μM Ligase 4 N148A, 8 μM E1, 5 mM ATP, 6 mM MgCl_2_ and 5 mM MESNa in 10 mM HEPES pH 7.5, 100 mM NaCl at 37 °C.

No ubiquitylation activity was observed with yeast PCNA, which adopts the same overall fold as human PCNA but shares only 35% sequence identity (Fig. 4D). Thus, PL4.1 discriminates between PCNA homologues with conserved overall structure, indicating that substrate recognition depends on specific features of the designed protein–protein interface rather than overall target shape alone.

We investigated whether the same cascade could be used to generate site-specifically polyubiquitylated PCNA with a defined chain length. Using the *de novo* ligase cascade with PL4.1, linear M1-linked diubiquitin, triubiquitin and tetraubiquitin chains were transferred to the target lysine with near-quantitative PCNA conversion (Fig. 4E).

## Discussion

We demonstrate that *de novo* enzyme design can generate target- and site-specific ubiquitin ligases for near-quantitative protein ubiquitylation. 35 of 48 tested designs transferred ubiquitin to a model peptide, with four achieving near-quantitative conversion without experimental optimisation. For the substantially more complex target PCNA, four of 48 designs showed ubiquitylation activity, one of which selectively modified K164 and, following a single active-site substitution, achieved near-quantitative conversion. Because target recognition and lysine selectivity are specified at the design stage rather than inherited from an evolved ligase, the same pipeline can in principle be directed at any lysine on any protein for which a structure or confident model is available.

Natural ubiquitylation distributes activation, conjugation and substrate selection across an E1–E2–E3 cascade, with specificity encoded in more than 600 human E3 ligases^4^. Our designs collapse the E2 and E3 functions into a single protein and make selectivity for both the ubiquitin donor as well as the acceptor lysine on the target protein programmable by design, agnostic to the target protein. Coupling the designed ligase to E1-dependent ubiquitin activation enabled site-specific ubiquitylation directly from unactivated ubiquitin without requiring an E2 or native E3 ligase. The same cascade also transferred defined di-, tri- and tetraubiquitin chains, extending the approach to site-specific installation of defined ubiquitin architectures. Delivered into cells, designed ligases could make individual ubiquitylation events installable on demand, turning proteome-wide site ubiquitylation maps^3^ from a catalogue into a set of testable hypotheses. Beyond dissecting regulation, the ability to direct ligases towards selected lysines could ultimately enable installation of degradative or regulatory ubiquitin signals onto a protein of choice. The accompanying work on designed small-molecule ligases shows that the underlying catalytic strategy is not restricted to ubiquitin transfer^23^. Similar design principles involving an activated acyl donor and a correctly positioned lysine should give access to *de novo* writers for the many lysine acylations for which the natural donors are acyl-CoA thioesters as for example lysine acetylation^24^, malonylation, succinylation, glutarylaton^25^ or lactylation^26^.

The PCNA ligase is a milestone in enzyme design, integrating extended and specific substrate recognition with the precise catalytic geometry required for ubiquitin transfer. The theozyme comprised 14 enzyme residues together with the five C-terminal residues of ubiquitin. Scaffolding this large motif while positioning K164 for catalysis on a polar protein surface beside a partly unresolved loop distinguishes this challenge from previous *de novo* enzyme design tasks. Selective ubiquitylation of PCNA shows that current methods can meet these combined demands, supporting extension of the platform to other biologically relevant targets. We expect advances in protein design tools such as RFdiffusion4-Proteina to further expand this capability^27^.

## Acknowledgements

This work was funded by the German Research Foundation (DFG) through SFB 1309 (project ID 325871075; to Michael Groll), from ETH Zurich and the European Research Council (ERC under the European Union’s Horizon 2020 research and innovation programme, grant agreement no. 101003289─Ubl-tool to K.L.), by an EMBO postdoctoral fellowship ALTF 226-2026 to M.G, DARPA D26AP50001 (T.F., S.S.M), a Schmidt fellowship to A.C.H., Schmidt Sciences (D.B.) and the Howard Hughes Medical Institute (D.B.). The authors thank the staff of beamline X06DA at the Paul Scherrer Institute (PSI, Villigen, Switzerland) for their support during data collection. Beamtime was allocated under proposal 20260403.

## References

1. Walsh, C. T., Garneau-Tsodikova, S. & Gatto Jr., G. J. Protein Posttranslational Modifications: The Chemistry of Proteome Diversifications. Angew. Chem. Int. Ed. 44, 7342–7372 (2005).

2. Dikic, I. & Schulman, B. A. An expanded lexicon for the ubiquitin code. Nat. Rev. Mol. Cell Biol. 24, 273–287 (2023).

3. van Gerwen, J. et al. The functional landscape of the human ubiquitinome. bioRxiv 2025.10.08.681129 (2025) doi:10.1101/2025.10.08.681129.

4. Chua, N. K. et al. The E3-ome gene-centric compendium reveals the human E3 ligase landscape. Cell 189, 2167–2188.e27 (2026).

5. Kriegesmann, J. & Brik, A. Synthesis of ubiquitinated proteins for biochemical and functional analysis. Chem. Sci. 14, 10025–10040 (2023).

6. Hofmann, R., Akimoto, G., Wucherpfennig, T. G., Zeymer, C. & Bode, J. W. Lysine acylation using conjugating enzymes for site-specific modification and ubiquitination of recombinant proteins. Nat. Chem. 12, 1008–1015 (2020).

7. Fottner, M. et al. Site-specific ubiquitylation and SUMOylation using genetic-code expansion and sortase. Nat. Chem. Biol. 15, 276–284 (2019).

8. Mossessova, E. & Lima, C. D. Ulp1-SUMO Crystal Structure and Genetic Analysis Reveal Conserved Interactions and a Regulatory Element Essential for Cell Growth in Yeast. Mol. Cell 5, 865–876 (2000).

9. Choi, H. et al. De novo design of cysteine proteases. Nature 1–9 (2026) doi:10.1038/s41586-026-11143-w.

10. Chen, A. et al. Computational design of metalloproteases. Nature 1–10 (2026) doi:10.1038/s41586-026-11167-2.

11. Butcher, J. et al. Atomic functional site scaffolding in biopolymers with RFdiffusion3. Science 0, eaeh2982 (2026).

12. Dauparas, J. et al. Atomic context-conditioned protein sequence design using LigandMPNN. Nat. Methods 22, 717–723 (2025).

13. Abramson, J. et al. Accurate structure prediction of biomolecular interactions with AlphaFold 3. Nature 630, 493–500 (2024).

14. Hoege, C., Pfander, B., Moldovan, G.-L., Pyrowolakis, G. & Jentsch, S. RAD6-dependent DNA repair is linked to modification of PCNA by ubiquitin and SUMO. Nature 419, 135–141 (2002).

15. Garg, P. & Burgers, P. M. Ubiquitinated proliferating cell nuclear antigen activates translesion DNA polymerases η and REV1. Proc. Natl. Acad. Sci. 102, 18361–18366 (2005).

16. Carlile, C. M., Pickart, C. M., Matunis, M. J. & Cohen, R. E. Synthesis of Free and Proliferating Cell Nuclear Antigen-bound Polyubiquitin Chains by the RING E3 Ubiquitin Ligase Rad5*. J. Biol. Chem. 284, 29326–29334 (2009).

17. Freudenthal, B. D., Gakhar, L., Ramaswamy, S. & Washington, M. T. Structure of monoubiquitinated PCNA and implications for translesion synthesis and DNA polymerase exchange. Nat. Struct. Mol. Biol. 17, 479–484 (2010).

18. Hibbert, R. G. & Sixma, T. K. Intrinsic Flexibility of Ubiquitin on Proliferating Cell Nuclear Antigen (PCNA) in Translesion Synthesis*. J. Biol. Chem. 287, 39216–39223 (2012).

19. Yang, K. et al. Chemical Protein Ubiquitylation with Preservation of the Native Cysteine Residues. ChemBioChem 17, 995–998 (2016).

20. Cao, L. et al. Design of protein-binding proteins from the target structure alone. Nature 605, 551–560 (2022).

21. Zhang, J. Z. et al. De novo design of Ras isoform selective binders. Cell Chem. Biol. 33, 396–408.e7 (2026).

22. Qian, T. et al. Complex enzyme active site scaffolding by iteratively detuned catalytic guidance. <2026.10.02.756307 Preprint at 10.64898/2026.10.02.756307 (2026).>

23. Salike, S. et al. Programmable amide bond formation using de novo ligases. (2026) Manuscript in preparation.

24. Ali, I., Conrad, R. J., Verdin, E. & Ott, M. Lysine Acetylation Goes Global: From Epigenetics to Metabolism and Therapeutics. Chem. Rev. 118, 1216–1252 (2018).

25. Hirschey, M. D. & Zhao, Y. Metabolic Regulation by Lysine Malonylation, Succinylation, and Glutarylation *. Mol. Cell. Proteomics 14, 2308–2315 (2015).

26. Ren, H., Tang, Y. & Zhang, D. The emerging role of protein L-lactylation in metabolic regulation and cell signalling. Nat. Metab. 7, 647–664 (2025).

27. Corley, N. et al. Biomolecular design through latent reasoning with RFD4-Proteina. (2026) https://rfd4-proteina.org/assets/paper/RFD4-Proteina.pdf.

