## Supplementary Information for "*De novo* design of ligases for programmable site-specific protein ubiquitylation"

#### Table of contents

1 Supplementary figures

2 Supplementary methods

2.1 Computational methods

2.1.1 Model Peptide–ubiquitin design campaign

2.1.1.1 Catalytic motif and molecular representations

2.1.1.2 Backbone generation and complementary-strand sampling

2.1.1.3 Peptide burial and opening angles

2.1.1.4 Sequence design and structure-prediction metrics

2.1.1.5 Partial diffusion and refinement from predicted complexes

2.1.1.6 State-specific structural selection

2.1.2 PCNA Ligase pipeline

2.1.2.1 Molecular representations and catalytic metrics

2.1.2.2 Backbone generation

2.1.2.3 PCNA interface and pocket selection

2.1.2.4 Initial sequence design and AF3 selection

2.1.2.5 Sequence refinement with decreasing RF3 guidance

2.1.2.6 Composite scoring and backbone family selection

2.1.2.7 Confidence-dependent sequence redesign

2.1.2.8 AF3 evaluation of apo and PCNA-bound acyl states

2.2 Experimental methods

2.2.1 General methods: plasmids and reagents

2.2.2 Protein expression and purification

2.2.2.1 Generation of Ub-MESNa by intein thiolysis

2.2.2.2 Expression and purification of Ub

2.2.2.3 Generation of Ub-MESNa by E1-mediated capture

2.2.2.4 Expression and purification of the SUMO–peptide fusions and the tagless peptides

2.2.2.5 Expression and purification of de novo designed ligases in plate format

2.2.2.6 Expression and purification of de novo designed ligases in shaker flasks

2.2.2.7 Expression and purification of human PCNA and yeast PCNA
2.2.2.8 Expression and purification of linear di/tri/tetraUb
2.2.2.9 Expression and purification of Ubch5c and H6-UBA1
2.2.3 In vitro de novo Ub ligase assays
2.2.3.1 LC-MS based screening assay for de novo Ub-peptide ligases
2.2.3.2 LC-MS based kinetic characterisation of Ub-peptide ligases
2.2.3.3 LC-MS based thioester donor specificity assays
2.2.3.4 In vitro ubiquitylation of SUMO-peptide fusion with Ub-peptide ligase 29
2.2.3.5 SDS-PAGE based screening assay for de novo Ub-PCNA ligases activity
2.2.3.6 In vitro PCNA ubiquitylation assays with Ub-PCNA ligase 4 and its variants
2.2.3.7 E1-coupled in vitro (poly)ubiquitylation of PCNA
2.2.3.8 Ubch5c-mediated ubiquitylation of PCNA
2.2.4 Intact protein mass spectrometry
2.2.5 X-ray crystallography
3 Amino acid sequences of proteins
3.1 General proteins
3.2 De novo ubiquitin ligases
4 Supplementary references

### 1 Supplementary figures

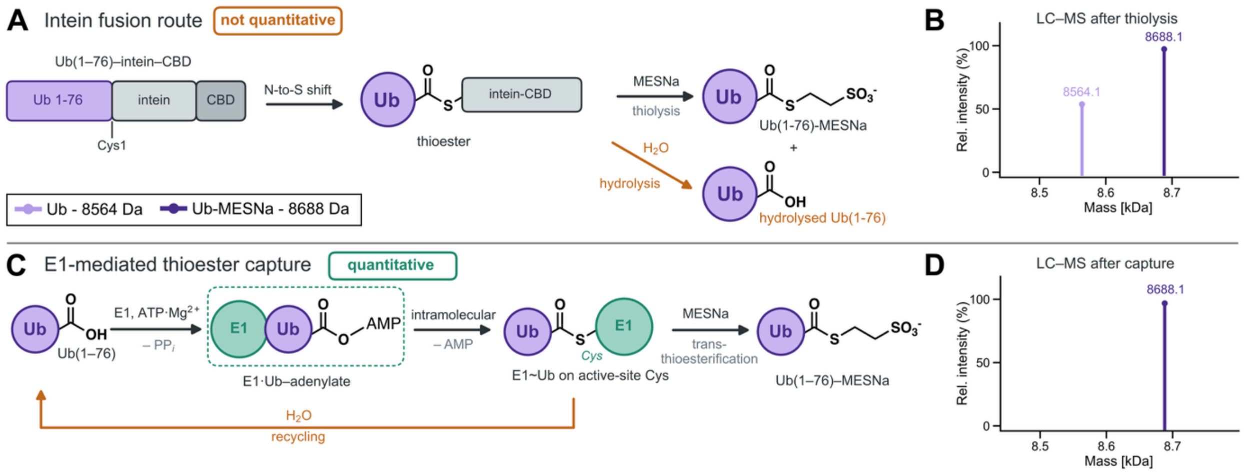

**Supplementary Figure 1 | Generation of the Ub(1-76)-MESNa thioester by intein thiolysis and by E1-mediated capture. (A)** Route via an intein fusion. Ub(1-76) is expressed as an N-terminal fusion to an MxeGyrA-chitin-binding domain (CBD) cassette. An N-to-S acyl shift onto the N-terminal cysteine of the intein (Cys1) converts the amide junction into a thioester, which is cleaved by thiolysis with MESNa to release Ub(1-76)-MESNa. Hydrolysis of the thioester competes with thiolysis and liberates unactivated Ub(1-76), limiting the yield of Ub-MESNa. **(B)** Deconvoluted mass spectrum of the thiolysis eluate. Two species are observed: Ub(1-76)-MESNa (obs. 8688.1 Da) and hydrolysed Ub(1-76) (obs. 8564.1 Da), showing that the intein route does not deliver the thioester quantitatively. **(C)** Route via E1-mediated capture. Uba1 (E1) binds Ub(1-76) together with ATP-$Mg_{2+}$ and catalyses both steps of C-terminal activation: adenylation of the Gly76 carboxylate with release of PPi gives the E1-bound Ub-adenylate, which is then attacked intramolecularly by the E1 active-site cysteine with release of AMP to give the E1~Ub thioester. Transthioesterification with 20 mM MESNa releases Ub(1-76)-MESNa and regenerates free E1. Ub(1-76) liberated by hydrolysis re-enters the cycle, driving the reaction to completion. **(D)** Deconvoluted mass spectrum of the capture reaction. A single species is observed, Ub(1-76)-MESNa (obs. 8688.1 Da), with no detectable unactivated Ub(1-76), demonstrating quantitative thioester formation. Deconvoluted masses are plotted as relative intensity normalised to the base peak of each spectrum. Consistent results were obtained over 3 distinct replicate experiments.

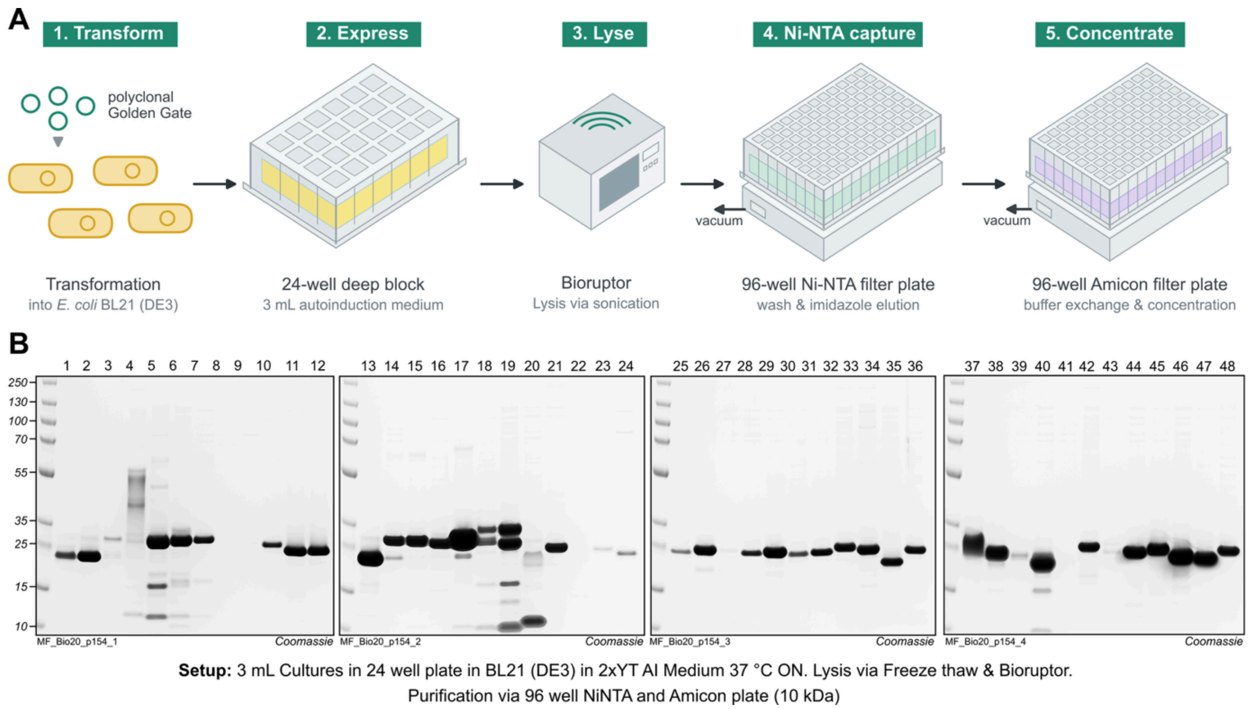

**Supplementary Figure 2 | Parallel expression and purification of the 48 de novo designed ubiquitin-peptide ligases. (A)** Schematic depiction of the workflow. The Golden Gate assembly product of each design was transformed into *E. coli* BL21 (DE3), and the resulting polyclonal transformants were cultured in 3 mL in 2xYT autoinduction medium in a 24-well deep block at 37 °C overnight (1/2). Cells were lysed by freeze-thaw followed by sonication in a Bioruptor (3). Clarified lysates were transferred to a 96-well Ni-NTA filter plate, where the His-tagged ligases were captured, washed and eluted with imidazole-containing buffer using a vacuum manifold (4), and the eluates were buffer-exchanged and concentrated on a 96-well Amicon filter plate (10 kDa MWCO), again under vacuum (5). All steps downstream of transformation were carried out in plate format, allowing the 48 designs to be expressed and purified in parallel. **(B)** SDS-PAGE analysis of the purified ligases visualised by Coomassie staining. Lanes are numbered by design (1-48).

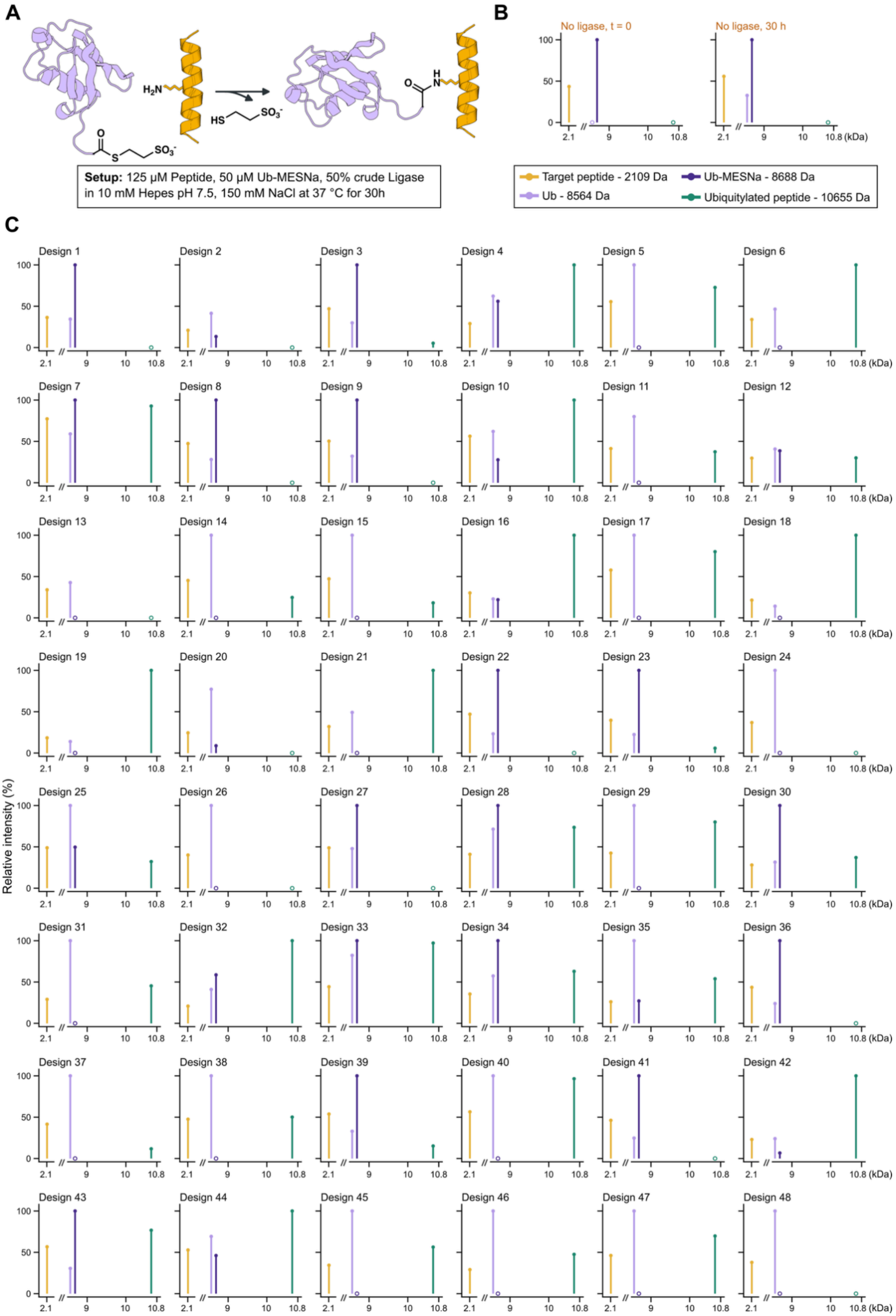
A
B
C
H2N
HS
SO3-
Setup: 125 µM Peptide, 50 µM Ub-MESNa, 50% crude Ligase in 10 mM Hepes pH 7.5, 150 mM NaCl at 37 °C for 30h
No ligase, t = 0
No ligase, 30 h
Target peptide - 2109 Da
Ub - 8564 Da
Ub-MESNa - 8688 Da
Ubiquitylated peptide - 10655 Da
Relative intensity (%)
(kDa)
Design 1
Design 2
Design 3
Design 4
Design 5
Design 6
Design 7
Design 8
Design 9
Design 10
Design 11
Design 12
Design 13
Design 14
Design 15
Design 16
Design 17
Design 18
Design 19
Design 20
Design 21
Design 22
Design 23
Design 24
Design 25
Design 26
Design 27
Design 28
Design 29
Design 30
Design 31
Design 32
Design 33
Design 34
Design 35
Design 36
Design 37
Design 38
Design 39
Design 40
Design 41
Design 42
Design 43
Design 44
Design 45
Design 46
Design 47
Design 48

**Supplementary Figure 3 | Screening of the 48 de novo designed ligases for ubiquitylation of the target helix. (A)** Reaction scheme. The ligase transfers ubiquitin from Ub(1-76)-MESNa onto the lysine side chain of the target helix, forming an isopeptide bond and releasing MESNa. Reactions contained 125 µM peptide, 50 µM Ub(1-76)-MESNa and 50% (v/v) crude ligase preparation in 10 mM HEPES pH 7.5, 150 mM NaCl and were incubated at 37 °C for 30 h. **(B)** No-ligase controls. Deconvoluted mass spectra of the complete reaction mixture lacking ligase, recorded at t = 0 and after 30 h. The target peptide (2109 Da) and Ub(1-76)-MESNa (8688 Da) are the only species present at t = 0; after 30 h, free Ub(1-76) (8564 Da) was formed through background hydrolysis of the thioester, but no ubiquitylated peptide (10655 Da) was detected at either time point, establishing that product formation is strictly ligase-dependent. **(C)** Deconvoluted mass spectra of the 48 designs after 30 h, one panel per design. Ubiquitylated peptide is detected for 35 of the 48 designs. Intensities are plotted relative to the base peak of each spectrum; the mass axis is broken between 2.4 and 8.45 kDa. Open circles on the baseline denote species that were not among the reported peaks of the respective deconvolution.

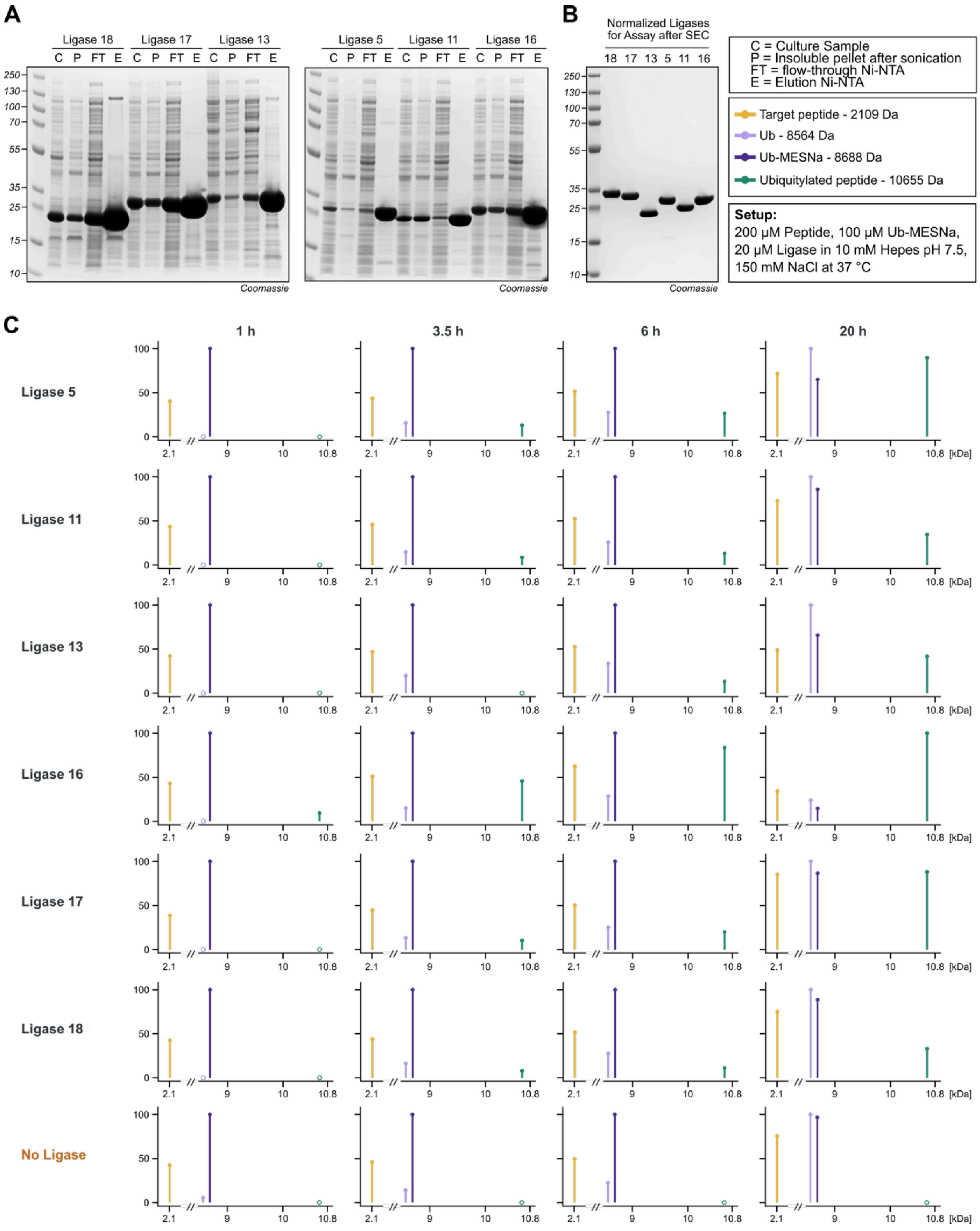

**Supplementary Figure 4 | Purification and kinetic characterisation of ligase designs 5, 11, 13, 16, 17 and 18 under defined conditions.** **(A)** Ni-NTA purification of the six designs selected from the primary screen, expressed from sequence-verified monoclonal cultures. For each design, samples of the culture (C), the insoluble pellet remaining after sonication (P), the Ni-NTA flow-through (FT) and the imidazole eluate (E) were analysed by SDS-PAGE and Coomassie staining. **(B)** Ni-NTA eluates were further polished by size-exclusion chromatography (SEC) and adjusted to equal concentration before use. SDS-PAGE with

Coomassie staining of the normalised preparations used in the assay. **(C)** Ubiquitylation time course for the six purified ligases and a no-ligase control. Reactions contained 200 µM peptide, 100 µM Ub(1-76)-MESNa and 20 µM ligase in 10 mM HEPES pH 7.5, 150 mM NaCl and were incubated at 37 °C. Reactions were analysed by LC-MS after 1, 3.5, 6 and 20 h. Intensities are plotted relative to the base peak of each spectrum; the mass axis is broken between 2.4 and 8.45 kDa. Open circles on the baseline denote species that were not among the reported peaks of the respective deconvolution.

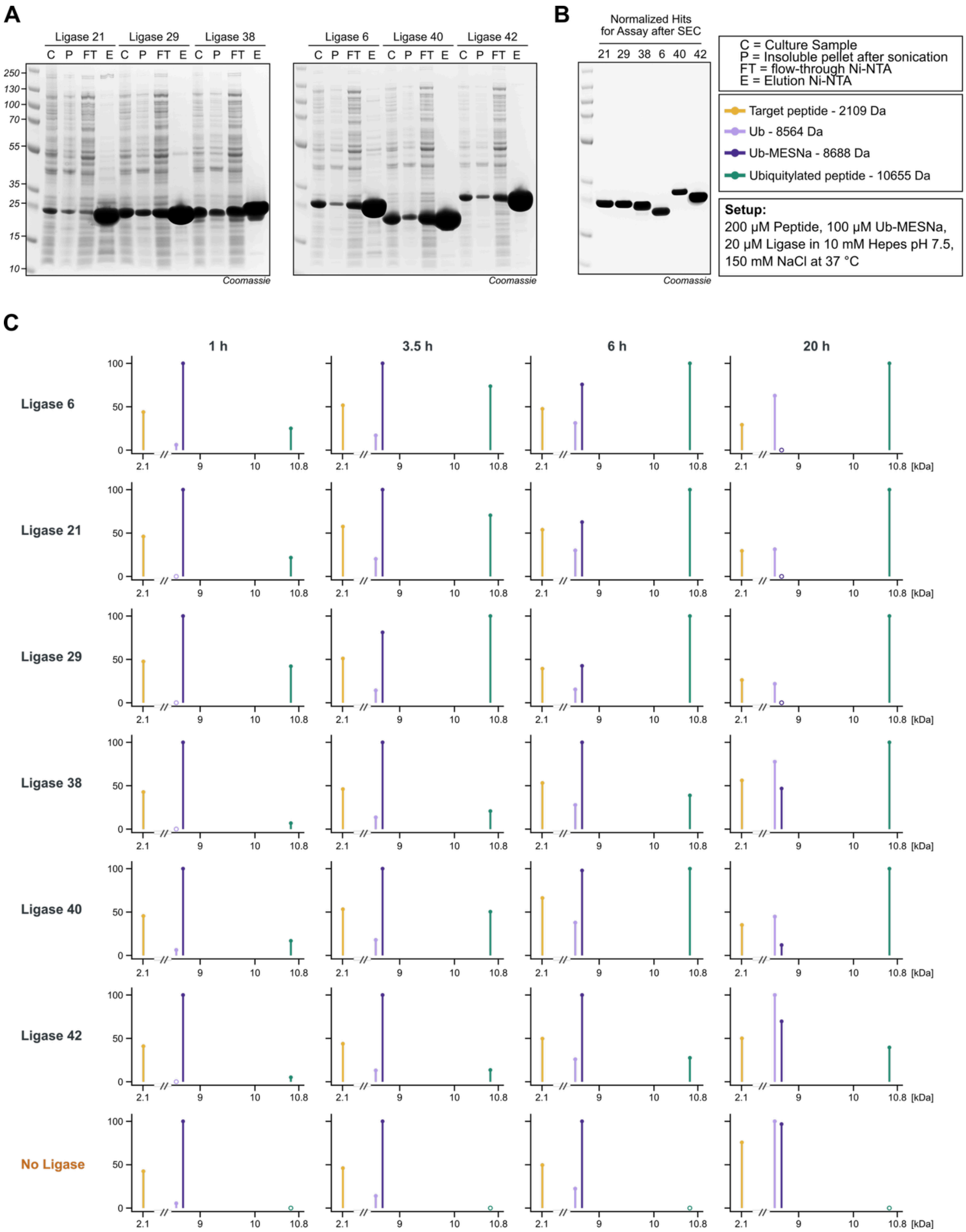

**Supplementary Figure 5 | Purification and kinetic characterisation of ligase designs 6, 21, 29, 38, 40 and 42 under defined conditions. (A)** Ni-NTA purification of the six designs selected from the primary screen, expressed from sequence-verified monoclonal cultures. For each design, samples of the culture (C), the insoluble pellet remaining after sonication (P), the Ni-NTA flow-through (FT) and the imidazole eluate (E) were analysed by SDS-PAGE and Coomassie staining. **(B)** Ni-NTA eluates were further polished by size-exclusion chromatography (SEC) and adjusted to equal concentration before use. SDS-PAGE with Coomassie staining of the normalised preparations used in the assay. **(C)** Ubiquitylation time course for the six purified ligases and a no-ligase control. Reactions contained 200 µM peptide, 100 µM Ub(1-76)-MESNa and 20 µM ligase in 10 mM HEPES pH 7.5, 150 mM NaCl and were incubated at 37 °C. Reactions were analysed by LC-MS after 1, 3.5, 6 and 20 h. Intensities are plotted relative to the base peak of each spectrum; the mass axis is broken between 2.4 and 8.45 kDa. Open circles on the baseline denote species that were not among the reported peaks of the respective deconvolution.

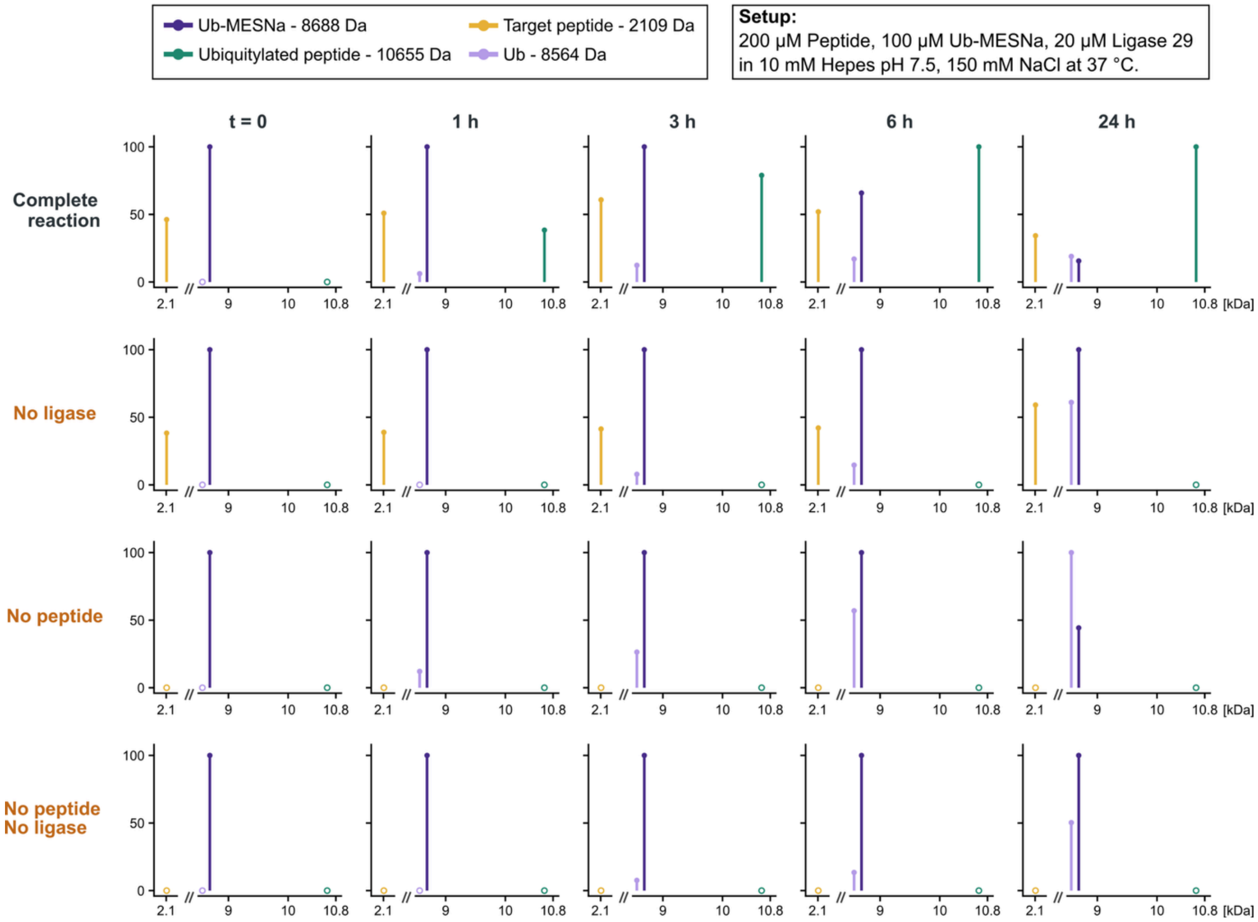

**Supplementary Figure 6 | Control reactions for ubiquitylation of the target helix by Ligase 29.** Deconvoluted mass spectra of the complete reaction and of three omission controls recorded at t = 0 and after 1, 3, 6 and 24 h. Reactions contained 200 µM peptide, 100 µM Ub(1-76)-MESNa and 20 µM Ligase 29 in 10 mM HEPES pH 7.5, 150 mM NaCl and were incubated at 37 °C. In the complete reaction the ubiquitylated peptide (10655 Da) appears within 1 h, becomes the base peak by 6 h and is accompanied by depletion of Ub(1-76)-MESNa (8688 Da). No ubiquitylated peptide is detected at any time point in any of the three controls, confirming that product formation requires both the ligase and the acceptor peptide. Free Ub(1-76) (8564 Da), formed by hydrolysis of the thioester, accumulates in all four conditions. Intensities are plotted relative to the base peak of each spectrum; the mass axis is broken between 2.4 and 8.45 kDa. Open circles on the baseline denote species that were not among the reported peaks of the respective deconvolution.

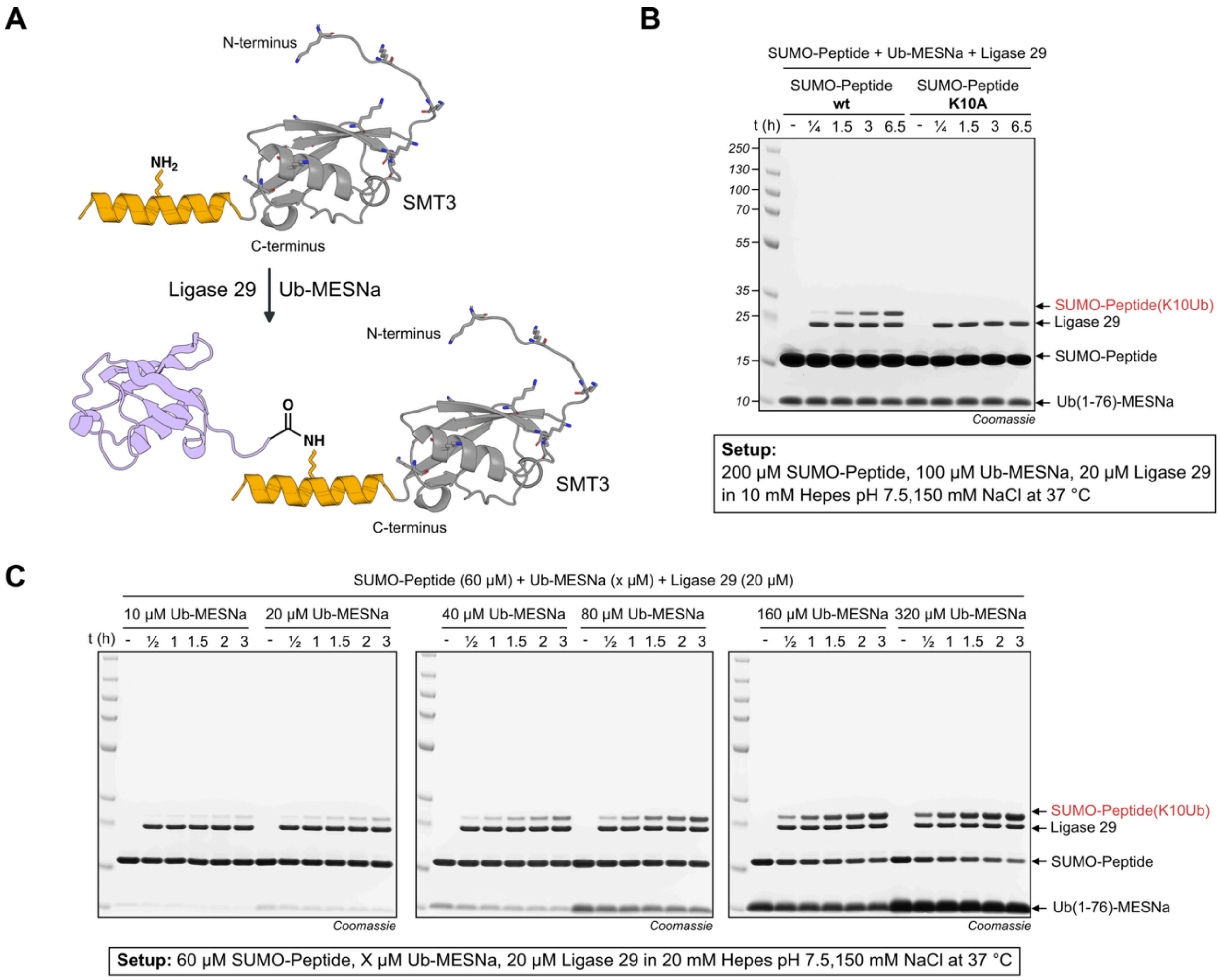

**Supplementary Figure 7 | Site-specific ubiquitylation of a SUMO-peptide fusion. (A)** Schematic representation of the SUMO-peptide substrate (top) and of its ubiquitylated product (bottom). The target helix (orange) is fused C-terminally to the Smt3 (SUMO) domain (grey). Smt3 itself contributes nine lysine side chains (shown as sticks), which are solvent-exposed thereby presenting multiple chemically equivalent acceptor sites in addition to the target lysine. The helix presents the acceptor lysine, whose free side-chain amine is highlighted; incubation with Ligase 29 and Ub(1-76)-MESNa attaches ubiquitin (lavender) to this lysine through an isopeptide bond. **(B)** Ligase 29 modifies the SUMO-peptide at K10. Reactions containing 200 µM SUMO-peptide, either wild type or the K10A helix variant, 100 µM Ub(1-76)-MESNa and 20 µM Ligase 29 in 10 mM HEPES pH 7.5, 150 mM NaCl were incubated at 37 °C and sampled after 0, 0.25, 1.5, 3 and 6.5 h; samples were resolved by SDS-PAGE and visualised by Coomassie staining. A band of the mass expected for mono-ubiquitylated SUMO-peptide (SUMO-Peptide(K10Ub), red arrow) accumulates with time in the wild-type reaction and is absent at every time point from the K10A reaction, identifying Lys10 as the modified residue. **(C)** Dependence of product formation on the concentration of the ubiquitin thioester. Reactions containing 60 µM SUMO-peptide, 20 µM Ligase 29 and 10, 20, 40, 80, 160 or 320 µM Ub(1-76)-MESNa in 20 mM HEPES pH 7.5, 150 mM NaCl were incubated at 37 °C and sampled after 0, 0.5, 1, 1.5, 2 and 3 h. Product formation increases with thioester concentration.

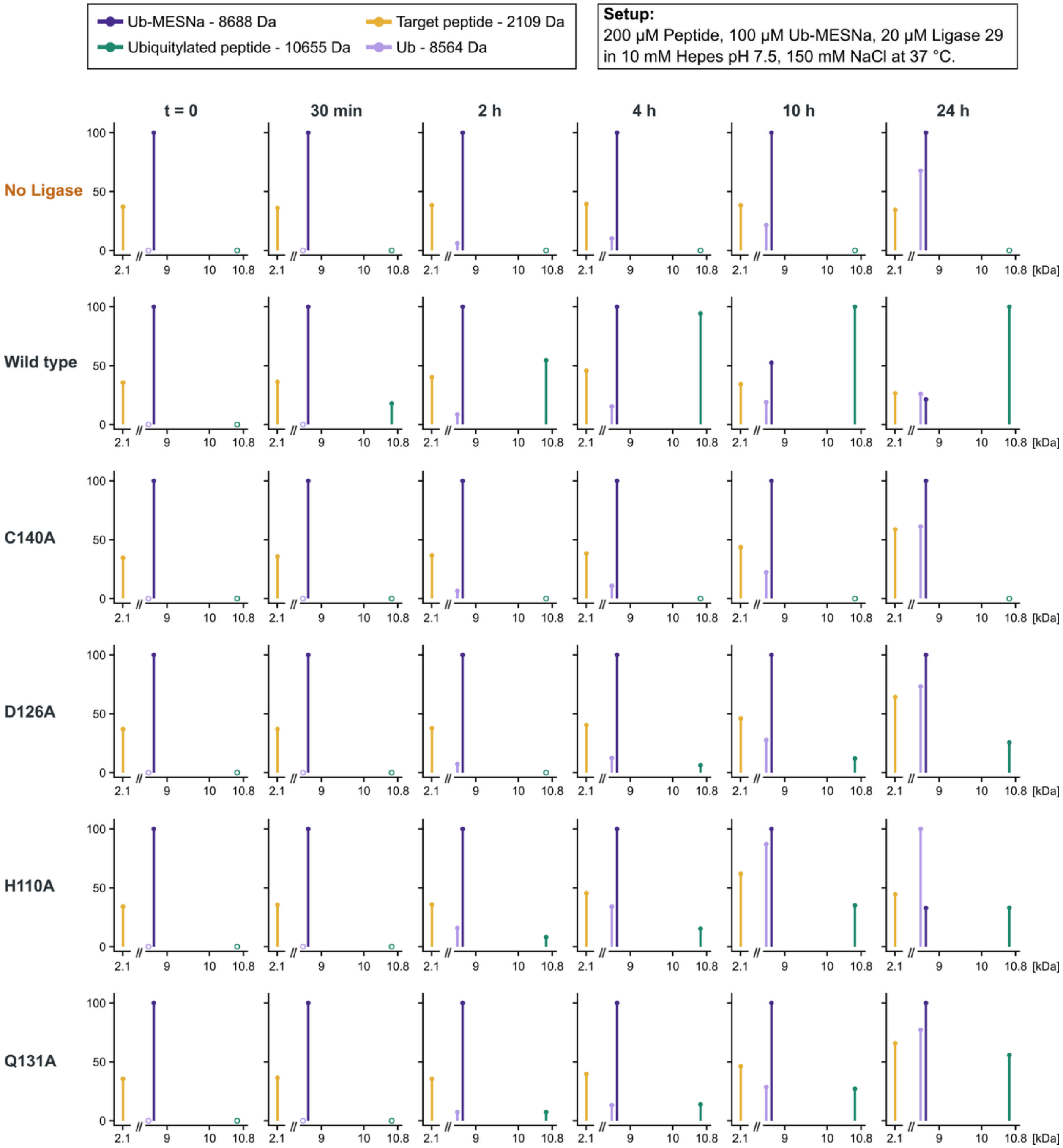

**Supplementary Figure 8 | Kinetic characterisation of catalytic tetrad mutants of Ligase 29.** Deconvoluted mass spectra of ubiquitylation reactions carried out with wild-type Ligase 29 or with the single active-site variants C140A, D126A, H110A and Q131A recorded at t = 0 and after 30 min, 2, 4, 10 and 24 h. A reaction without ligase is shown for comparison. Reactions contained 200 µM peptide, 100 µM Ub(1-76)-MESNa and 20 µM of the respective ligase in 10 mM HEPES pH 7.5, 150 mM NaCl and were incubated at 37 °C. The wild-type enzyme forms ubiquitylated peptide (10655 Da) within 30 min and converts the thioester almost completely by 24 h. Mutation of the catalytic cysteine (C140A) abolishes activity reproducing the no-ligase control. The three remaining substitutions are strongly impaired but retain residual activity, forming product only from 2-4 h onwards. Together, the data are consistent with C140 acting as the catalytic nucleophile and with H110, D126 and Q131 contributing to, but not being essential for, catalysis. Intensities are plotted relative to the base peak of each spectrum; the mass axis is broken between 2.4 and 8.45 kDa. Open circles on the baseline denote species that were not among the reported peaks of the respective deconvolution.

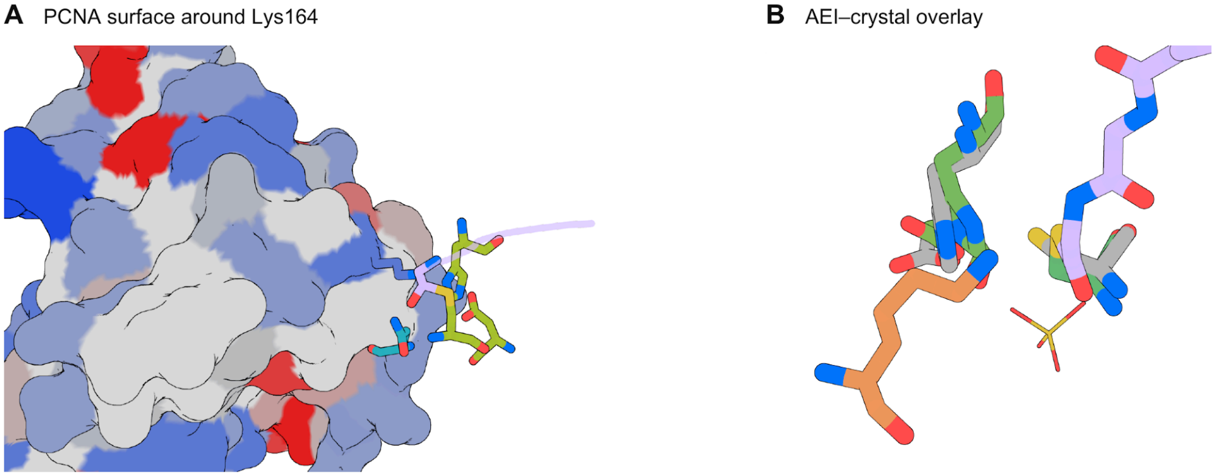

**Supplementary Figure 9 | Surface chemistry around PCNA Lys164 and structural comparison of the ligase active site. (A)** PCNA surface around Lys164 with the catalytic motif placed in the geometry of the second tetrahedral intermediate (TI2). The motif is shown as green and teal sticks, Lys164 as blue sticks and the donor segment as a lavender cartoon. Surface colours report Eisenberg residue hydrophobicity, graded from blue (hydrophilic) through grey (zero) to red (hydrophobic). Colours are assigned to whole residues only when their polar side-chain groups or nonpolar side chains meet solvent-exposure criteria; remaining residues are grey. Exposure was calculated for isolated PCNA in PyMOL using a 1.4-Å water probe. No prominent exposed hydrophobic patch is apparent immediately around Lys164, highlighting the challenge of simultaneously scaffolding the active site and designing binding interactions around the target lysine. **(B)** Superposition of the AlphaFold3-predicted N148A ligase acyl-enzyme intermediate (AEI) bound to PCNA with the ligase crystal structure (grey), aligned over 198 enzyme Cα atoms. The predicted enzyme is green, PCNA Lys164 orange and the ubiquitin acyl donor lavender; the crystallographic sulfate is yellow with red oxygens. One sulfate oxygen lies within 0.7 Å of the predicted acyl carbonyl oxygen.

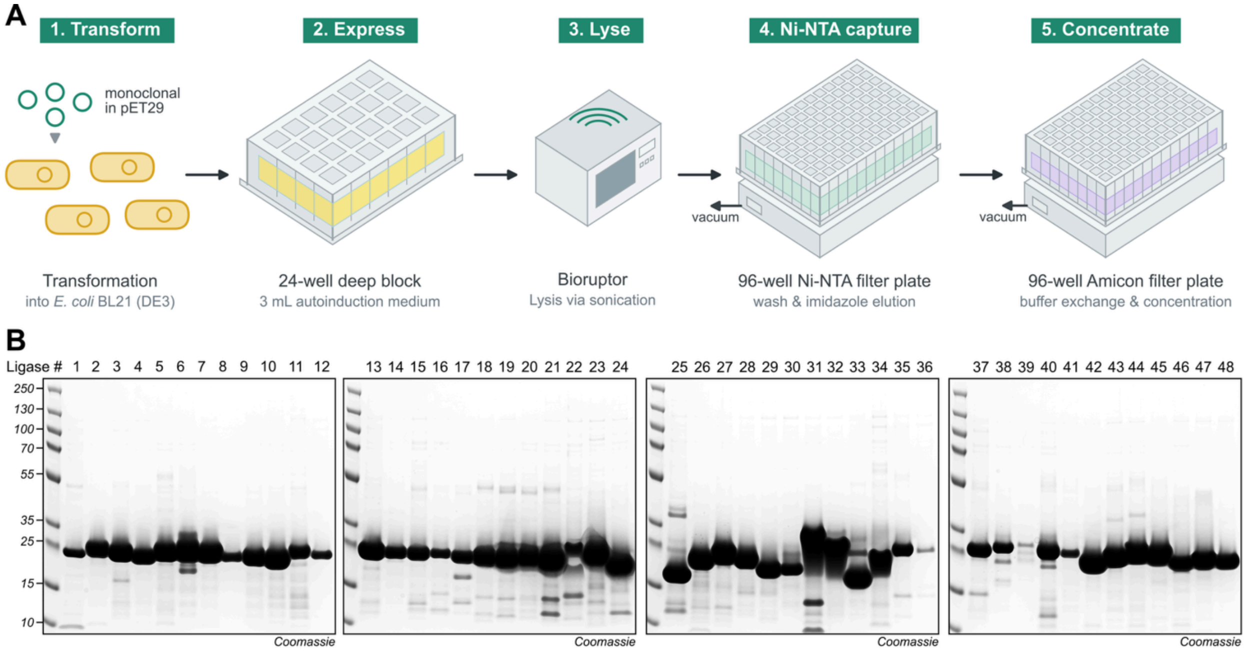

**Setup:** 3 mL Cultures in 24 well plate in BL21 (DE3) in 2xYT AI Medium 37 °C ON. Lysis via Freeze thaw + Bioruptor followed by 96 well NiNTA and Amicon plate (10 kDa)

**Supplementary Figure 10 | Parallel expression and purification of the 48 de novo designed ubiquitin-PCNA ligases. (A)** Schematic depiction of the workflow. Sequence-verified monoclonal Ub-PCNA Ligase designs in pET29 vectors were transformed into *E. coli* BL21 (DE3) followed by expression in 3 mL in 2xYT autoinduction medium in a 24-well deep block at 37 °C overnight (1/2). Cells were lysed by freeze-thaw followed by sonication in a Bioruptor (3). Clarified lysates were transferred to a 96-well Ni-NTA filter plate, where the His-tagged ligases were captured, washed and eluted with imidazole-containing buffer using a vacuum manifold (4); the eluates were then buffer-exchanged and concentrated on a 96-well Amicon filter plate (10 kDa MWCO), again under vacuum (5). All steps downstream of transformation are carried out in plate format, allowing the 48 designs to be expressed and purified in parallel. **(B)** SDS-PAGE analysis of the purified ligases visualised by Coomassie staining. Lanes are numbered by design (1-48).

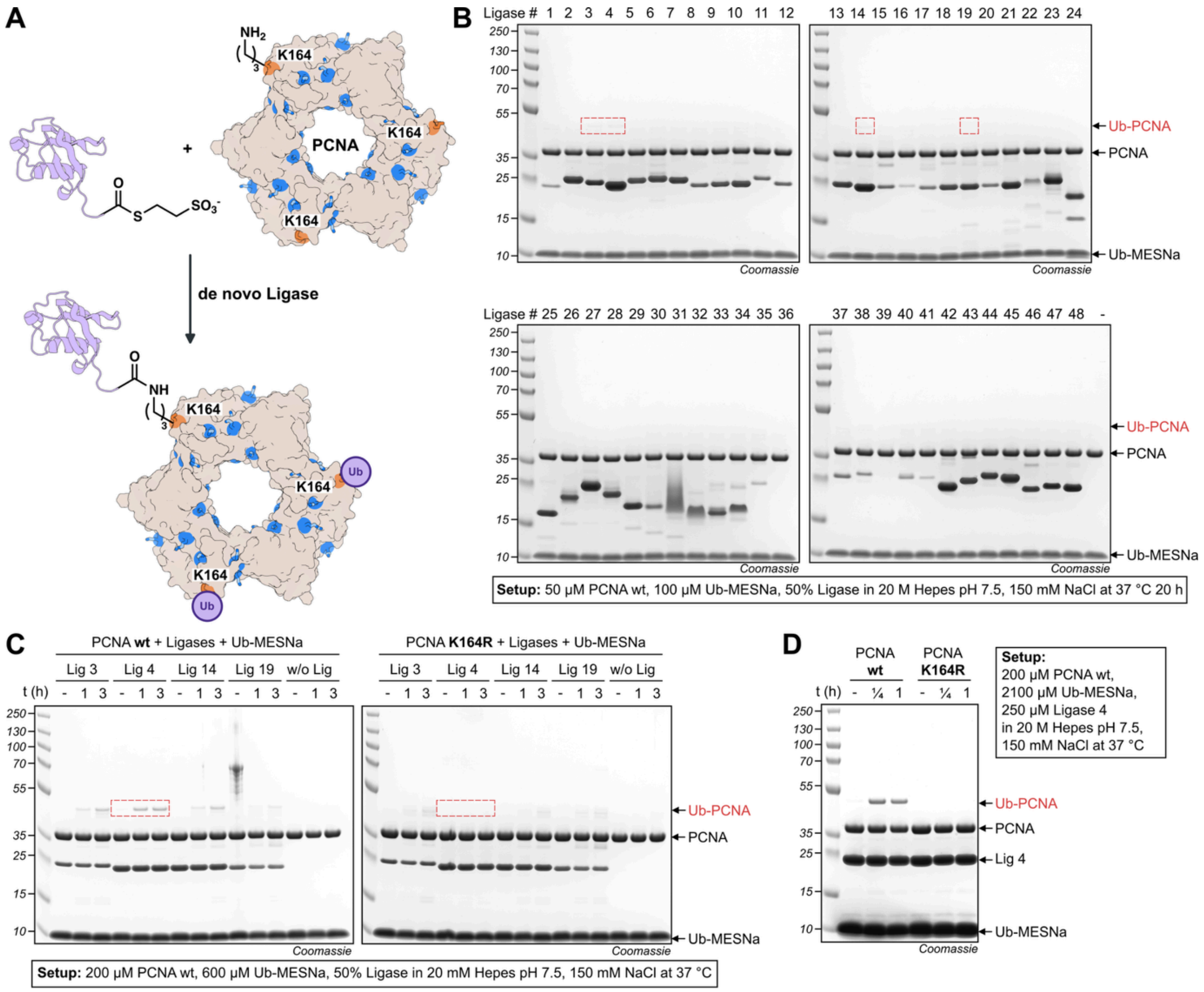

**Supplementary Figure 11 | Screening of the de novo designed ligases for ubiquitylation of PCNA at K164. (A)** Schematic depiction of a de novo designed ligase transferring ubiquitin from Ub(1-76)-MESNa onto K164 of the PCNA homotrimer, forming an isopeptide bond. PCNA is shown in surface representation with its lysine side chains in blue and Lys164, the physiological ubiquitylation site, highlighted. **(B)** Primary screen of the 48 designs. Reactions containing 50 µM PCNA, 100 µM Ub(1-76)-MESNa and 50% (v/v) of the respective ligase preparation in 20 mM HEPES pH 7.5, 150 mM NaCl were incubated at 37 °C for 20 h and analysed by SDS-PAGE with Coomassie staining. Lanes are numbered by ligase design. Faint product bands are detected for designs 3, 4, 14 and 19 (boxed). **(C)** Follow-up of the four hits on wild-type and K164R PCNA. Reactions containing 200 µM PCNA, either wild type or the K164R variant, 600 µM Ub(1-76)-MESNa and 50% (v/v) of the respective ligase in 20 mM HEPES pH 7.5, 150 mM NaCl were incubated at 37 °C and sampled after 0, 1 and 3 h; a reaction without ligase (w/o Lig) is included for each substrate. Designs 3, 14 and 19 generate a product band of the same mobility on wild-type and on K164R PCNA, showing that these ligases modify lysines other than Lys164 and are not site-specific. In contrast, Ligase 4 produces a product that is lost upon mutation of the target lysine (boxed), identifying K164 as the site of modification. **(D)** Ubiquitylation of PCNA by Ligase 4 at elevated reagent concentrations. Reactions containing 200 µM PCNA, wild type or K164R, 2100 µM Ub(1-76)-MESNa and 250 µM Ligase 4 in 20 mM HEPES pH 7.5, 150 mM NaCl were incubated at 37 °C and sampled after 0, 0.25 and 1 h. A distinct Ub-PCNA band is formed within 15 min on wild-type PCNA and is not detected with K164R, confirming site-specific modification of K164.

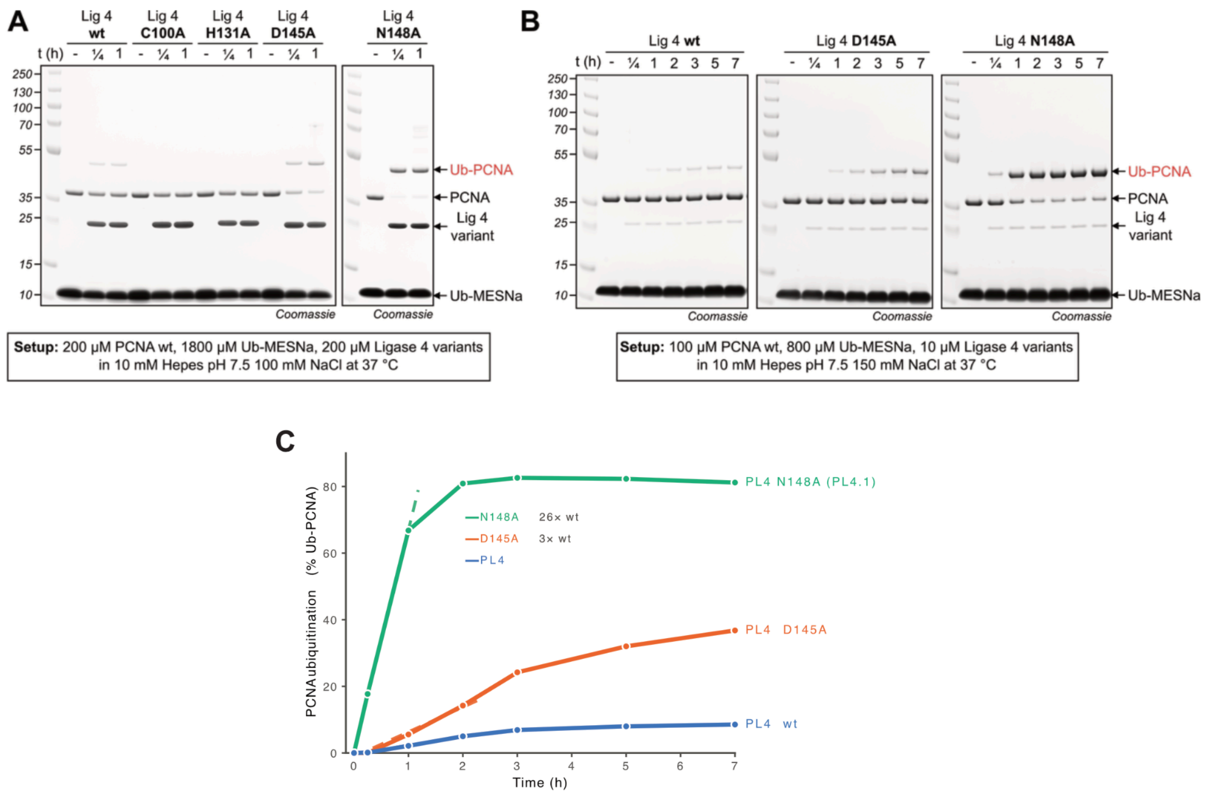

**Supplementary Figure 12 | The active-site variant Ligase 4 N148A (PL4.1) is a fast and site-specific PCNA ubiquitin ligase. (A)** Mutational analysis of the catalytic tetrad of Ligase 4. Reactions containing 200 µM PCNA, 1800 µM Ub(1-76)-MESNa and 200 µM of the indicated Ligase 4 variant in 10 mM HEPES pH 7.5, 100 mM NaCl were incubated at 37 °C and sampled after 0, 15 min and 1 h followed by SDS-PAGE analysis with Coomassie staining. The wild-type ligase produces only traces of Ub-PCNA, and no product is formed by C100A or H131A, identifying the Cys-His pair as catalytically essential. D145A and, most pronounced, N148A convert the majority of PCNA within 15 min (**(B)** Time course of the active-site substitutions. Reactions containing 100 µM PCNA, 800 µM Ub(1-76)-MESNa and 10 uM of wild-type Ligase 4, Ligase 4 D145A or Ligase 4 N148A in 10 mM HEPES pH 7.5, 150 mM NaCl were incubated at 37 °C and sampled after 0, 15 min, 1, 2, 3, 5 and 7 h. **(C)** Dependence on thioester donor concentration. Reactions containing 100 µM PCNA, 10 µM Ligase 4 N148A and 100, 200, 400, 800 or 1600 µM Ub(1-76)-MESNa in 10 mM HEPES pH 7.5, 100 mM NaCl were incubated at 37 °C and sampled after 0, 5 min, 15 min, 1 and 2 h. Product formation increases with donor concentration. At 800 µM and above, PCNA is almost completely consumed within 2 h. **(C)** Densitometry analysis of data shown in (B). After 1h, N148A shows 66% conversion, D145A 7.4% conversion and wt 2.6% conversion. Comparing initial rates gives a 26-fold improvement for N148A and a 3-fold improvement for D145A over PL4..

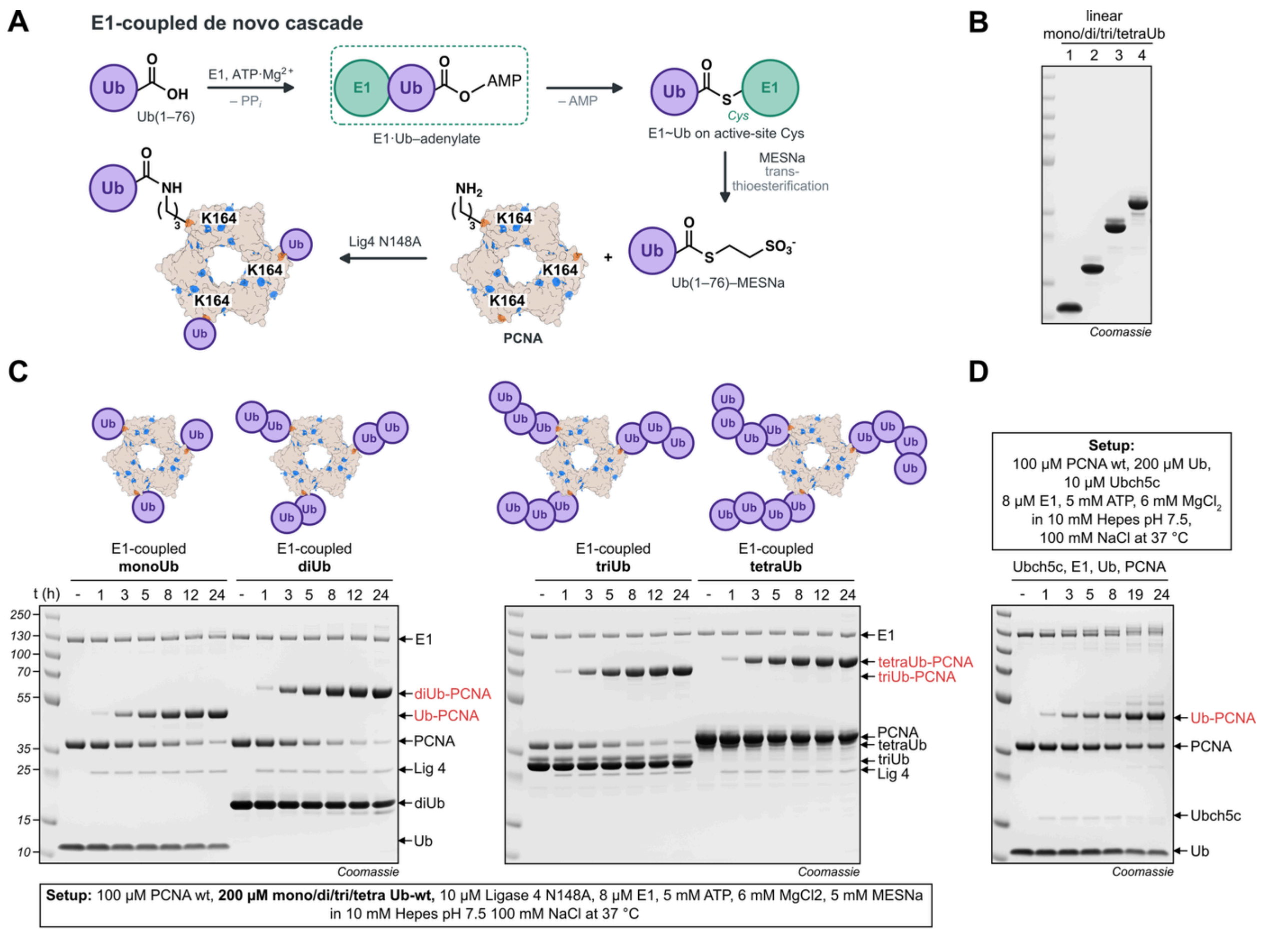

**Supplementary Figure 13 | An E1-coupled cascade generates the ubiquitin thioester in situ and extends the system to linear ubiquitin chains. (A)** Scheme of the E1-coupled de novo cascade. E1 activates the C-terminal carboxylate of Ub(1-76) as the ubiquitin adenylate under consumption of ATP and $Mg^{2+}$ and transfers ubiquitin onto its active-site cysteine with release of AMP. Transthioesterification with MESNa releases Ub(1-76)-MESNa, which Ligase 4 N148A uses to modify Lys164 of PCNA. All components are combined in one pot, so that the thioester donor is formed and consumed in situ and does not have to be prepared and purified separately. **(B)** Purified linear ubiquitin chains used as donors. Mono- and M1-linked di-, tri- and tetraubiquitin (lanes 1 to 4) **(C)** One-pot transfer of mono- and linear polyubiquitin onto PCNA. Reactions containing 100 µM PCNA, 200 µM of Ub or the respective linear ubiquitin species, 10 µM Ligase 4 N148A, 8 µM E1, 5 mM ATP, 6 mM $MgCl_2$ and 5 mM MESNa in 10 mM HEPES pH 7.5, 100 mM NaCl were incubated at 37 °C and sampled after 0, 1, 3, 5, 8, 12 and 24 h. Each donor is transferred onto PCNA, giving Ub-PCNA, diUb-PCNA, triUb-PCNA and tetraUb-PCNA; product appears within 1 h in every case and PCNA is largely consumed by the end of the time course. Cartoons above the gels indicate the expected modification of the PCNA homotrimer. **(D)** Comparison with E3-independent ubiquitylation by Ubch5c[1]. The promiscuous E2 Ubch5c (UBE2D3) is the most widely used alternative for installing ubiquitin on a target protein in vitro in the absence of a cognate E3. A reaction containing 100 µM PCNA, 200 µM ubiquitin, 10 µM Ubch5c, 8 µM E1, 5 mM ATP and 6 mM MgCl2 in 10 mM HEPES pH 7.5, 100 mM NaCl was incubated at 37 °C and sampled after 0, 1, 3, 5, 8, 19 and 24 h. Mono-ubiquitylated PCNA accumulates slowly and is accompanied from 8 h onwards by additional, more slowly migrating products and by further ubiquitylated species at the top of the gel, indicating unspecific modification of additional lysines.

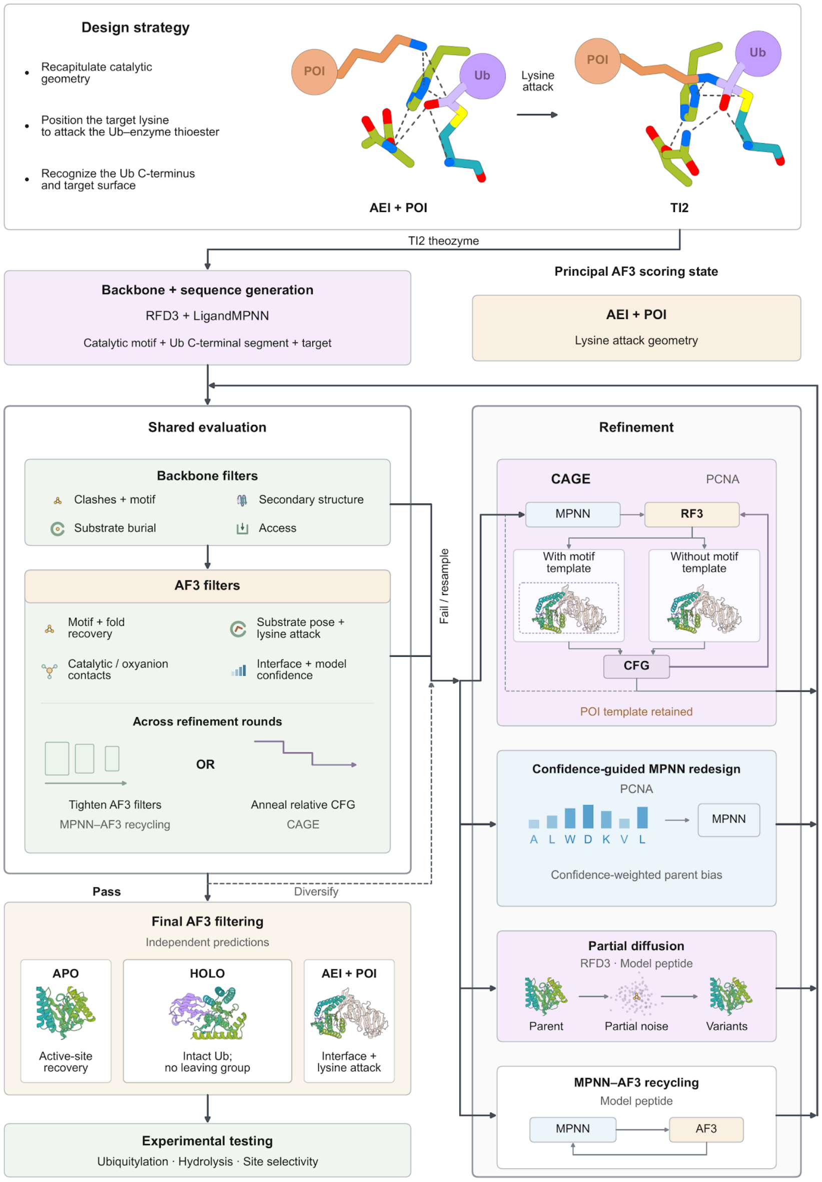

**Supplementary Figure 14 | Computational workflow for ubiquitin-ligase design.** TI2 theozymes combine the Cys–His–Asp catalytic triad, oxyanion-hole donors, ubiquitin C-terminal segment and target for RFdiffusion3 (RFD3) backbone generation and MPNN sequence design. Backbone filters assess clashes, motif retention, secondary structure, substrate burial and access. AlphaFold3 (AF3) evaluates fold, motif and substrate-pose recovery, catalytic and oxyanion contacts, lysine attack geometry and prediction confidence, principally in AEI plus protein of interest (POI). For the model peptide, refinement uses partial diffusion of selected original RFD3 backbones and MPNN–AF3 recycling with progressively tighter AF3 filters. For PCNA, the Catalytic Guidance Engine (CAGE) alternates MPNN redesign with RF3 prediction. Classifier-free guidance (CFG) combines RF3 prediction updates with and without relative motif templating; relative CFG decreases across redesign rounds while the absolute PCNA template is retained. A separate PCNA route biases LigandMPNN continuously towards parent amino acids according to AF3 confidence, with stronger retention bias at more confident positions. Refined candidates return to shared evaluation. Final independent AF3 predictions in the relevant states assess active-site recovery (APO), intact-ubiquitin accommodation (HOLO), and interface and lysine attack geometry (AEI plus target) before experimental tests of ubiquitylation, donor hydrolysis and site selectivity. The purple Ub and orange POI spheres provide schematic protein context; molecular views show design or predicted structures. TI2, second tetrahedral intermediate; AEI, acyl-enzyme intermediate; APO, unmodified enzyme alone; HOLO, unmodified enzyme with intact ubiquitin.

### 2 Supplementary methods

#### 2.1 Computational methods

##### Overall design pipeline

We designed ligases to recognize the ubiquitin C-terminus and target surface and position a specified lysine for attack on the ubiquitin–enzyme thioester (Supplementary Fig. 14). Both campaigns began from a theozyme of the second tetrahedral intermediate (TI2), which we scaffolded with RFdiffusion3 (RFD3) before MPNN sequence design. We used structure prediction and iterative refinement to improve recovery of the catalytic motif and placement of the reacting amine.

We principally evaluated lysine attack geometry with AlphaFold3 (AF3) in the acyl-enzyme intermediate (AEI) bound to the acceptor. We encoded the RLRGG ubiquitin tail as a noncanonical modification of catalytic cysteine, forming the thioester, and supplied the acceptor as a separate chain. HOLO predictions paired unmodified enzyme with intact ubiquitin, without an acceptor or donor leaving group; APO predictions contained enzyme alone. For PCNA, we sampled multiple random seeds and selected confident subsets of the AF3 ensembles for structural assessment. The next two sections detail the model-peptide (HELIX–ubiquitin) and PCNA design campaigns.

##### 2.1.1 Model Peptide–ubiquitin design campaign

We designed enzymes to position the C-terminal RLRGG segment of ubiquitin and a model peptide acceptor for lysine attack on a cysteine-bound acyl intermediate. The catalytic motif specified the reacting groups, while a complementary enzyme β-strand provided backbone contacts to orient the donor tail. We combined RFdiffusion3 (RFD3)[2] scaffold generation with three rounds of ProteinMPNN sequence design and AlphaFold3 (AF3)[3] evaluation. Alternative complementary-strand placements were sampled before scaffolding, and partial diffusion expanded the selected backbone families before the second sequence-design round. Selection required recovery of the enzyme fold, catalytic motif and substrate poses, followed by assessment of donor-tail binding without the acceptor.

###### 2.1.1.1 Catalytic motif and molecular representations

The catalytic motif comprised Cys, His and Asp, together with a Gln side chain and the cysteine backbone amide positioned to stabilise the oxyanion. The model acceptor peptide had the sequence SIAEAFREAKEYLLAAAQQ, with Lys10 as the reacting residue. The donor was represented by the five-residue RLRGG tail, and its terminal glycine supplied the acyl carbonyl. The acceptor and donor sequences were fixed throughout the campaign.

The input motif used reference labels A3, A8, A12 and A14 for Cys, His, Asp and Gln, respectively, B1–B19 for the acceptor, and D94–D98 for the RLRGG tail. Reference residues C140–C141 specified the complementary strand and were incorporated into the generated enzyme. These reference labels were mapped to the corresponding positions in each generated structure. In the resulting three-chain models, chain A was the enzyme, chain B the acceptor, and chain C the donor tail, numbered C1–C5. We denote the enzyme residue corresponding to reference C141 as P, and the enzyme residue two positions N-terminal to it as P−2.

For acyl-state AF3 prediction, the donor tail was represented as a custom modification of the catalytic cysteine, with the terminal glycine carbonyl carbon bonded to Cys Sγ. The acceptor remained a separate chain. Within this modification, C36 and O37 were the terminal donor carbonyl carbon and oxygen. For subsequent sequence design and relaxation, the modified residue was separated into standard cysteine and an RLRGG chain while preserving the predicted coordinates. The same carbonyl atoms then corresponded to C and O of donor Gly5. Apo predictions contained the unmodified enzyme alone. Holo predictions contained the unmodified enzyme and a separate RLRGG peptide, without the helical acceptor; the donor occupied chain B, residues 1–5. These predictions assessed recognition of the donor tail without modelling an activated-ester leaving group or the remainder of ubiquitin.

**Supplementary Table 1 | Motif atoms specified during RFD3 scaffolding and used for catalytic-motif RMSD.** Backbone denotes N, Cα, C and O; labels refer to the input motif.

| Reference residue | Fixed coordinates during scaffolding | Atoms used for catalytic-motif RMSD |
|---|---|---|
| A2 | Backbone | Backbone |
| A3, Cys | Backbone, Cβ, Sγ | Backbone, Cβ, Sγ |
| A4 | Backbone | Backbone |
| A5, A9, A10, A11, A13 | Backbone | None |
| A8, His | Backbone, Cβ, Cγ, Nδ1, Cδ2, Cε1, Nε2 | Cβ, Cγ, Nδ1, Cδ2, Cε1, Nε2 |
| A12, Asp | Backbone, Cβ, Cγ, Oδ1, Oδ2 | Backbone, Cβ, Cγ, Oδ1, Oδ2 |
| A14, Gln | Cγ, Cδ, Oε1, Nε2 | Cγ, Cδ, Oε1, Nε2 |
| B1–B19, helical acceptor | All supplied atoms | None |
| C140–C141, complementary enzyme strand | Backbone | None |
| D94–D98, donor tail | Backbone | None |

**2.1.1.2 Backbone generation and complementary-strand sampling**

RFD3 scaffolding used a total-length parameter of 175–275 residues. The indexed enzyme segments comprised reference A8–A10 and A11–A13, connected by 10–14 generated residues and flanked by 40–150 generated residues at each terminus. The cysteine-containing segment A2–A5, Gln A14 and complementary-strand residues C140–C141 were included without fixed sequence positions. The acceptor and donor tail were specified as separate chains. Amino-acid identities were released at reference A2, A4, A5, A9, A10, A11, A13, C140 and C141 during initial scaffolding.

Alternative complementary-strand placements were sampled while retaining the catalytic fragment, acceptor and donor-tail coordinates. Each placement was scaffolded with the origin centred on either catalytic Cys Sγ or acceptor Lys Nζ. The two origin coordinates were (2.886, 3.648, −5.019) and (0.038, 4.139, −5.645) Å, respectively, in the input-motif frame. Each origin was perturbed independently along each Cartesian axis by a uniform displacement between −4 and +4 Å. Generation used 200 diffusion steps, an origin-jitter parameter of 3.0, step scale 2 and $\gamma_0$ = 0.8, with virtual atoms and guideposts removed from the resulting structures.

Backbone selection combined chain integrity, local secondary structure and enzyme secondary-structure content with burial and opening-angle criteria (Supplementary Table 2). Secondary structure was assigned using DSSP, with H classified as helix (a), E/B as strand (b), and all other assignments as c. The enzyme non-loop fraction was the combined helix and strand fraction. Three-residue windows centred on reference A2, A3, A8 and A12 were required to match, respectively, {cca, ccc}, {cca, caa}, {cbb, ccb, bbb, bbc, bcc} and {cbb, ccb, bbb, bbc, bcc}. All four windows had to pass.

**2.1.1.3 Peptide burial and opening angles**

We combined acceptor burial with openings around both peptide components to favour a groove that accommodated the donor tail and helical acceptor. This geometry was intended to shield the reacting groups while preserving access for substrate binding and product release. Burial and angular clearance were structural selection criteria, rather than measurements of water access or product inhibition.

Acceptor burial was defined as $1 - \mathrm{clip}(SASA_{complex}/SASA_{isolated}, 0, 1)$, where $SASA_{complex}$ was the summed solvent-accessible surface area of chain B in the full complex and $SASA_{isolated}$ was its surface area at the same coordinates with all other chains removed. The ratio was clipped to [0, 1], giving burial values from 0 for unchanged solvent exposure to 1 for complete loss of accessible surface. The calculation included occlusion by both the enzyme and donor tail. Donor-tail burial was calculated by the same procedure for chain C, but backbone selection used only acceptor burial. Surface areas were calculated with the Shrake–Rupley algorithm in Biotite using element-based Single radii, a 1.4 Å probe radius and 1,000 Fibonacci-distributed surface points per atom.

An opening angle was calculated separately for the acceptor and donor tail. For each peptide, the longitudinal axis was the first principal component of its mean-centred N, Cα and C coordinates, and the projection origin was the centroid of these atoms. N, Cα and C atoms of amino-acid residues in all other chains were projected onto the plane perpendicular to this axis. The projected azimuthal angles were sorted, and the largest gap between consecutive angles, including the interval across 360°, defined the opening angle. Thus, the acceptor calculation included the enzyme and donor tail, whereas the donor calculation included the enzyme and acceptor. Both calculations used atom centres without radial or axial cutoffs; larger angles indicated a wider unoccupied sector in the corresponding projection.

Initial scaffolds required acceptor burial ≥0.45, an acceptor opening angle of 40–180° and a donor opening angle ≥15°. After partial diffusion, the same burial and minimum-angle criteria were applied without the upper acceptor-angle limit (Supplementary Table 2).

**Supplementary Table 2 | Backbone-selection criteria.** All criteria within each column were required.

| Criterion | Initial scaffolds | Partial-diffusion scaffolds |
|---|---|---|
| RFD3-reported chain breaks | 0 | Not applied |
| Four motif secondary-structure windows | All pass | All pass |
| Enzyme helix/strand fraction | ≥0.65 | ≥0.65 |
| Acceptor burial | ≥0.45 | ≥0.45 |

| | | |
|---|---|---|
| Acceptor opening angle | 40–180°, inclusive | ≥40° |
| Donor-tail opening angle | ≥15° | ≥15° |

**2.1.1.4 Sequence design and structure-prediction metrics**

ProteinMPNN designed enzyme sequences at a sampling temperature of 0.1, with side-chain packing and side-chain context enabled. Cysteine, methionine and catalytic residues were excluded at designable positions. The catalytic Cys, His and Asp, oxyanion-hole Gln, complete acceptor and complete donor tail retained their identities. The two complementary-strand residues retained the identities assigned during initial scaffolding. Other scaffold residues and the backbone-only enzyme motif positions A2, A4, A5, A9, A10, A11 and A13 were designable. AF3 inputs used the designed enzyme sequence with empty paired and unpaired multiple-sequence alignments and no protein structural templates. Acyl predictions included the cysteine modification at its mapped position and the fixed helical acceptor sequence. Separate apo and holo predictions used the corresponding unmodified enzyme sequence. Structural metrics were calculated from the returned AF3 model for each input.

Enzyme Cα RMSD measured recovery of the fold after alignment to the corresponding sequence-design or relaxed input structure. For acyl and apo predictions, catalytic-motif RMSD was calculated after independently fitting the 32 atoms in Supplementary Table 1 to the reference motif. Acceptor RMSD used the N, Cα, C and O atoms of residues B8–B12 after alignment on enzyme Cα atoms, measuring placement of the segment surrounding Lys10 relative to the enzyme. Donor-tail RMSD similarly used the N, Cα, C and O atoms of all five donor residues after enzyme Cα alignment. The acceptor and donor comparisons used the corresponding design input as reference, with the donor reconstructed as a separate chain for the acyl-state calculation. Round 1 selected acyl predictions for recovery of the fold, catalytic motif and both peptide poses, followed by apo predictions for recovery of the fold and catalytic motif without either peptide (Supplementary Table 3).

**2.1.1.5 Partial diffusion and refinement from predicted complexes**

Backbone family selection retained the trajectory with the lowest acyl-state acceptor RMSD from each original RFD3 backbone after round 1. Partial diffusion used the original generated coordinates, with partial times of 1, 5, 10 and 15, while retaining the catalytic Cys–His–Asp/Gln motif, the complete acceptor and donor tail, and the complementary-strand motif residues. The resulting backbones were screened using the partial-diffusion criteria in Supplementary Table 2 and provided the inputs for round 2.

Rounds 2 and 3 alternated ProteinMPNN sequence sampling, side-chain packing and constrained Rosetta relaxation. Each relaxed structure supplied the coordinates for the next sequence-sampling cycle, and the outputs from all cycles were evaluated against the relaxation filters. The sequence restrictions and sampling temperature were unchanged from round 1. Round 3 started from passing round-2 acyl predictions converted to three-chain models. Round-3 sampling was distributed among surviving descendants within each original-backbone family.

**2.1.1.6 State-specific structural selection**

Round-2 AF3 filtering tightened fold-recovery limits and added distance criteria for placing the lysine amine near the donor carbonyl and catalytic histidine, together with the donor-carbonyl contact to the cysteine backbone amide. Candidates had to pass the acyl criteria, including donor-tail placement, before apo assessment. Donor-tail RMSD was applied after reconstruction of the donor chain as a separate filter following the other round-2 acyl criteria. Passing acyl structures supplied coordinates for round 3 only after the corresponding apo predictions passed. Round 3 tightened catalytic-motif RMSD and the donor-carbonyl–cysteine-amide distance, and added a Cys–His distance requirement. The two lysine-placement distances retained their 4.0 Å limits. Apo predictions then had to recover the fold and motif under the tighter round-3 criteria (Supplementary Table 3).

**Supplementary Table 3 | AF3 acyl and apo acceptance criteria.** All distances and RMSDs are in Å, and all inequalities are inclusive. "Not applied" denotes absence of an individual criterion from that round's filter.

| Criterion | Round 1 | Round 2 | Round 3 |
|---|---|---|---|
| Acyl enzyme Cα RMSD | ≤2.0 | ≤1.5 | ≤1.5 |
| Acyl catalytic-motif RMSD | ≤1.25 | ≤1.25 | ≤1.0 |
| Acyl acceptor B8–B12 RMSD | ≤2.0 | ≤1.5 | ≤1.5 |
| Acyl donor-tail RMSD | ≤2.5 | ≤2.5 | ≤2.5 |
| Acyl Lys Nζ–donor carbonyl C | Not applied | ≤4.0 | ≤4.0 |
| Acyl His Nδ1–Lys Nζ | Not applied | ≤4.0 | ≤4.0 |
| Acyl Cys Sγ–His Nδ1 | Not applied | Not applied | ≤4.0 |
| Acyl donor carbonyl O–Cys N | Not applied | ≤3.8 | ≤3.7 |
| Apo enzyme Cα RMSD | ≤2.0 | ≤1.5 | ≤1.5 |
| Apo catalytic-motif RMSD | ≤1.5 | ≤1.5 | ≤1.5 |

Candidates passing round-3 acyl and apo filters were evaluated in the holo state to test whether the enzyme retained its fold and positioned the Ubiquitin tail near the catalytic cysteine without the acceptor. Holo filtering required Ub–enzyme strand contacts and proximity of the terminal donor carbonyl carbon to Cys Sγ (Supplementary Table 4). For this state, catalytic-motif RMSD was measured after alignment on enzyme Cα atoms to the round-3 relaxed design, rather than after an independent fit of the motif atoms. The completed selection combined all three state-specific screens.

**Supplementary Table 4 | AF3 holo acceptance criteria.** The holo donor was free RLRGG, and P denotes the enzyme residue corresponding to reference C141. All values are in Å and inequalities are inclusive.

| Criterion | Threshold |
|---|---|

| | |
|---|---|
| Enzyme Cα RMSD | ≤1.5 |
| Catalytic-motif RMSD after enzyme Cα alignment | ≤1.0 |
| Cys Sγ–donor Gly5 C | ≤4.0 |
| Donor Leu2 O–P N | ≤4.0 |
| Donor Gly4 N–P O | ≤4.0 |

#### 2.1.2 PCNA Ligase pipeline

We designed ligases to position PCNA Lys164 for attack on a cysteine-bound acyl intermediate carrying the C-terminal ubiquitin tail. RFdiffusion3 (RFD3)[2] scaffolding, ProteinMPNN sequence design and AlphaFold3 (AF3)[3] selection established starting complexes for iterative refinement. Sequence design then alternated with RF3 structure prediction while relative motif guidance was progressively reduced. A related protocol used confidence-dependent sequence biases to redesign parent complexes from an earlier RF3 and AF3 refinement campaign, followed by ensemble assessment of the apo enzyme and the PCNA-bound acyl-enzyme intermediate.

**2.1.2.1 Molecular representations and catalytic metrics**

Initial scaffolding used the C-terminal domain of PCNA, and subsequent structure prediction and refinement used a 255-residue PCNA monomer. The donor was represented by the five-residue RLRGG ubiquitin tail throughout. The catalytic motif comprised Cys, His and Asp, together with an Asn side chain and the cysteine backbone amide positioned to stabilise the oxyanion. Reference labels A3, A8, A12 and A14 denoted Cys, His, Asp and Asn, respectively, and were mapped to the corresponding positions in each designed enzyme. In full-monomer models, PCNA occupied chain B with native residue numbering and Lys164 as the reacting residue. The donor tail corresponded to reference D94–D98 and was represented as chain C, residues 1–5, for sequence design and Rosetta relaxation.

For RF3 and acyl-state AF3 prediction, the donor tail was incorporated into a custom cysteine modification through a bond between the terminal glycine carbonyl carbon and Cys Sγ. Within this modification, C36 and O37 denoted the donor carbonyl carbon and oxygen. For sequence design, the modification was separated into standard cysteine and the RLRGG chain while preserving its coordinates. The corresponding carbonyl atoms were then C and O of donor Gly5. Apo AF3 predictions contained the unmodified enzyme alone.

Catalytic-motif RMSD was calculated after independent superposition of the 36 atoms in Supplementary Table 5. Initial AF3 and subsequent RF3 motif comparisons used the common PCNA theozyme reference, whereas AF3 comparisons after confidence-dependent redesign used the corresponding parent-derived sequence-design structure. Enzyme Cα RMSD measured recovery of the enzyme fold after Cα alignment to the structure supplied for sequence design. The principal catalytic distances measured Cys Sγ–His Nδ1, PCNA Lys164 Nζ–donor carbonyl C, His Nδ1–Lys164 Nζ and donor carbonyl O–Cys N. These distances assessed placement of the catalytic dyad, attacking amine and backbone oxyanion donor.

**Supplementary Table 5 | Motif atoms specified during RFD3 scaffolding and used for catalytic-motif RMSD.** Backbone denotes N, Cα, C and O; labels refer to the input motif.

| Reference residue | Fixed coordinates during scaffolding | Atoms used for catalytic-motif RMSD |
|---|---|---|
| A1, A5 | Backbone | None |
| A2 | Backbone | Backbone |
| A3, Cys | Backbone, Cβ, Sγ | Backbone, Cβ, Sγ |
| A4 | Backbone | Backbone |
| A7 | C, O | None |
| A8, His | Backbone, Cβ, Cγ, Nδ1, Cδ2, Cε1, Nε2 | Cβ, Cγ, Nδ1, Cδ2, Cε1, Nε2 |
| A9 | Backbone | None |
| A12, Asp | Cα, Cβ, Cγ, Oδ1, Oδ2 | Backbone, Cβ, Cγ, Oδ1, Oδ2 |
| A14, Asn | Cα, C, O, Cβ, Cγ, Oδ1, Nδ2 | Backbone, Cβ, Cγ, Oδ1, Nδ2 |
| A15, A16 | Backbone | None |
| B184, PCNA anchor | Backbone | None |
| C140–C141, complementary enzyme strand | Backbone | None |
| D94–D98, donor tail | Backbone | None |

**2.1.2.2 Backbone generation**

To position the lysine-containing PCNA surface beside the donor tail, RFD3 scaffolded the catalytic motif together with PCNA residues 127–188 and 191–255, the RLRGG tail and a complementary enzyme strand. The indexed enzyme segments comprised reference A7–A9, A12 and A14–A16, connected by 10–14 and two generated residues, respectively, and flanked by 40–150 generated residues at each terminus. Reference A1–A5 and complementary-strand residues C140–C141 were included without fixed sequence positions. The total indexed-contig length parameter was 200–350 residues, including the PCNA fragment and donor tail. The fixed motif atoms are listed in Supplementary Table 5. Amino-acid identities were released at A1, A2, A4, A5, A7, A9, A15, A16, C140 and C141, while the catalytic identities were retained.

Scaffolding sampled origins on a grid with 5 Å coordinate offsets around (−1.687, 6.302, −5.776) Å in the motif frame, and perturbed this central origin independently along each Cartesian axis by a uniform displacement between −2 and +2 Å. RFD3 generation used 200 diffusion steps, an origin-jitter parameter of 3.0, step scale 2 and $\gamma_0 = 0.8$.

**2.1.2.3 PCNA interface and pocket selection**

Backbone selection combined motif retention and secondary structure with geometric criteria intended to bury the reacting PCNA patch while limiting contacts with the surrounding PCNA surface (Supplementary Table 6). The enzyme non-loop fraction was the combined helix and strand fraction assigned using DSSP. Three-residue windows centred on reference A2, A3, A8 and A12 were required to match, respectively, {cca, ccc}, {cca, caa}, {cbb, ccb, bbb, bbc, bcc} and {cbb, ccb, bbb, bbc, bcc}, where a denotes DSSP H, b denotes E or B, and c denotes all other protein assignments. All four windows had to pass.

The lysine-tunnel axis joined Lys164 Nζ and catalytic Cys Sγ, and its length was the distance between these atoms. The tunnel opening was the largest empty angular sector after projection of enzyme Cα atoms onto the plane perpendicular to this axis through its midpoint. The calculation included only Cα atoms within 4 Å of this plane and 10 Å of the axis, excluding atoms within 3 Å of Cys Sγ. The donor-tail opening was defined separately using the first principal component of the donor N, Cα and C coordinates as the axis and their centroid as the projection origin. All enzyme N, Cα and C atoms were projected onto the perpendicular plane, and the largest angular gap, including the interval across 360°, defined the donor opening. This calculation excluded PCNA and used no axial or radial cutoffs.

Enzyme compactness was assessed by the radius of gyration of its heavy atoms. For each terminal enzyme residue, the terminal-pocket distance was the minimum distance from any of its heavy atoms to either Lys164 Nζ or Cys Sγ. The limits on terminal-pocket distances excluded backbones with either terminus close to the reacting groups.

**2.1.2.4 Initial sequence design and AF3 selection**

ProteinMPNN designed enzyme sequences at temperature 0.1 with side-chain packing and side-chain context enabled. The catalytic Cys, His, Asp and Asn identities and the PCNA and donor sequences were fixed. Cysteine, methionine and the catalytic residues were excluded at designable enzyme positions. AF3 predicted the resulting sequences in the PCNA-bound acyl state with empty paired and unpaired multiple-sequence alignments and no enzyme structural template. The PCNA sequence was expanded to the full monomer, with a structural template covering all residues except 189–190.

Starting sequences for RF3 refinement had to satisfy all five initial AF3 criteria in Supplementary Table 6. PCNA placement RMSD used backbone N, Cα, C and O atoms of residues 160–168 and the Lys164 side-chain atoms Cβ, Cγ, Cδ, Cε and Nζ after alignment on enzyme Cα atoms. The remaining metrics used the atom sets and alignments defined above. This selection evaluated individual AF3 predictions without ensemble aggregation.

To construct the RF3 starting complexes, the selected sequences retained their original ProteinMPNN enzyme and donor coordinates. The retained PCNA domain was restored to native residue numbering, and residues 1–126 were grafted from PDB 1AXC chain E after local alignment on PCNA residues 160–168. The terminal donor oxygen was restored from the corresponding RFD3 structure. AF3 thus selected starting sequences, while the grafted sequence-design structures supplied the coordinates for the initial RF3 prediction.

**Supplementary Table 6 | Initial backbone and AF3 sequence-selection criteria.** All criteria within each stage were required. Distances and RMSDs are in Å, and angles are in degrees; strict and inclusive inequalities are distinguished explicitly.

| Stage | Criterion | Threshold |
|---|---|---|
| Backbone | RFD3-reported chain breaks | ≤1 |
| Backbone | Motif insertion RMSD | ≤0.75 |
| Backbone | Four motif secondary-structure windows | All pass |
| Backbone | Enzyme helix/strand fraction | ≥0.65 |
| Backbone | Donor-tail opening angle | ≥40 |
| Backbone | Lysine-tunnel opening angle | ≤180 |
| Backbone | PCNA local-patch relative SASA | ≤0.35 |
| Backbone | PCNA nonlocal-patch relative SASA | ≥0.88 |
| Backbone | Nonlocal-minus-local patch relative SASA | ≥0.55 |
| Backbone | Enzyme radius of gyration | ≤18 |
| Backbone | Enzyme N- and C-terminal pocket distances | Each ≥10 |
| Initial AF3 | Enzyme Cα RMSD | <2.0 |
| Initial AF3 | Catalytic-motif RMSD | <2.0 |
| Initial AF3 | PCNA residues 160–168 placement RMSD | <3.0 |
| Initial AF3 | Lys164 Nζ–donor carbonyl C | <5.0 |
| Initial AF3 | Cys Sγ–His Nδ1 | <5.0 |

##### 2.1.2.5 Sequence refinement with decreasing RF3 guidance

The annealing protocol began from the grafted enzyme–PCNA–donor complexes described above. At each RF3 stage, PCNA coordinates in the incoming complex provided the absolute template, excluding residues 185–195 from the template selection. Relative guidance additionally specified the enzyme motif and its placement with respect to PCNA, including the donor-bearing cysteine modification. The relative classifier-free guidance[4] scale decreased through 0.75, 0.50, 0.25, 0.10 and 0.00, while the absolute PCNA template scale remained 1.0 throughout. The final stage therefore removed relative motif guidance while retaining structural guidance for PCNA.

The initial RF3 prediction used the supplied enzyme sequence. Subsequent stages redesigned sequences on the selected RF3 coordinates using ProteinMPNN, with sampling temperatures decreasing from 0.5 to 0.4, 0.25 and 0.1 (Supplementary Table 7). Only the enzyme sequence was designed; PCNA and RLRGG retained their sequences. The catalytic Cys, His, Asp and Asn identities were fixed, while the remaining enzyme positions were designable. Cysteine, methionine and catalytic residues were excluded at designable positions. Side-chain packing and side-chain context were enabled. Each selected prediction supplied the coordinates for the next sequence-design stage, allowing subsequent sequences to accommodate the structure recovered under weaker guidance.

**Supplementary Table 7 | RF3 relative-guidance and sequence-design schedule.** The absolute PCNA template guidance scale was 1.0 at every stage. The terminal stage includes iterative sequence design and constrained Rosetta relaxation.

| Stage | Relative guidance scale | Sequence-design temperature | Structure used for sequence design |
|---|---|---|---|
| Initial prediction | 0.75 | Supplied sequence | Starting complex |
| First redesign | 0.50 | 0.50 | Selected 0.75 prediction |
| Second redesign | 0.25 | 0.40 | Selected 0.50 prediction |
| Third redesign | 0.10 | 0.25 | Selected 0.25 prediction |
| Terminal redesign | 0.00 | 0.10 | Selected 0.10 prediction |

**2.1.2.6 Composite scoring and backbone family selection**

To retain alternative sequences while favouring catalytic geometry, the current RF3 protocol combines individual metrics into a weighted composite score (Supplementary Table 8). For a distance or RMSD x, the component score is clip[(b − x)/(b − a), 0, 1], where a is the full-score boundary and b the zero-score boundary. The clash component is 1 when RF3 reports no clash and 0 otherwise; missing metric values contribute 0. The composite score is the weighted mean of the six components, with a total weight of 6.5. The component boundaries are soft preferences, so a candidate can compensate for a low value in one component through higher values in the others.

The current selection rule requires a composite score ≥0.5 and retains at most ten candidates per original RFD3 backbone family. Candidates are ordered by decreasing composite score, with higher enzyme-chain pLDDT resolving ties and the input order resolving any remaining ties. No global rank cutoff is applied. In the completed initial 0.75-guidance stage, backbone family selection instead retained the highest-ranked candidate from each family without a minimum composite score; the subsequent 0.50, 0.25 and 0.10 stages used the score threshold and family cap above.

**Supplementary Table 8 | Soft components of the RF3 composite score.** Distances and RMSDs are in Å. Continuous scores decrease linearly between the full-score and zero-score boundaries.

| Component | Full-score boundary, a | Zero-score boundary, b | Weight |
|---|---|---|---|
| Catalytic-motif RMSD | 1.0 | 2.0 | 1.5 |

| | | | |
|---|---|---|---|
| Cys Sγ–His Nδ1 | 4.0 | 4.5 | 1.0 |
| Lys164 Nζ–donor carbonyl C | 4.0 | 5.0 | 1.0 |
| His Nδ1–Lys164 Nζ | 4.0 | 5.0 | 1.0 |
| Donor carbonyl O–Cys N | 3.0 | 3.8 | 1.0 |
| RF3-reported clash | Absent | Present | 1.0 |

**2.1.2.7 Confidence-dependent sequence redesign**

To preserve sequence features that supported reproducible structure predictions while allowing less certain regions to change, we biased LigandMPNN towards the parent amino acid according to its prediction confidence. Starting complexes and confidence values came from AF3 evaluation of parents in an earlier RF3 refinement campaign. For each enzyme residue, p was the mean Cα pLDDT across 50 parent acyl-state predictions containing PCNA. The additive bias for the parent amino acid was $B(p) = 5/[1 + \exp\{-0.25(p - 80)\}]$. Values with $B(p) \leq 0.1$ were omitted, and retained biases were rounded to two decimal places. This continuous function weakly favoured the parent identity at low-confidence positions and approached a bias of 5 at high-confidence positions. The bias applied throughout the designable enzyme, including residues near the donor and PCNA, with no pocket-distance exclusion. It favoured retention of the parent identity without fixing it; the catalytic Cys, His, Asp and Asn were separately fixed.

LigandMPNN sequence sampling used temperature 1.0, side-chain context and side-chain packing. Only chain A was designed, and PCNA and RLRGG retained their identities. Cysteine and the catalytic residues were excluded at designable positions; methionine remained allowed. LigandMPNN redesigned and packed sequences on the parent backbone without intervening Rosetta relaxation.

**2.1.2.8 AF3 evaluation of apo and PCNA-bound acyl states**

AF3 predictions of the redesigned apo enzyme assessed fold recovery, while predictions of the PCNA-bound acyl-enzyme intermediate assessed the acceptor interface and catalytic geometry. Each state used two randomised seeds and five diffusion samples per seed, giving ten predicted structures for ensemble analysis. The enzyme was predicted without a protein structural template. For the PCNA-containing state, a PCNA structural template covered the acceptor except residues 189–190; the apo state contained no protein structural template.

The evaluation protocol calculates metrics separately for every available ensemble member and averages members with valid values for each metric. Enzyme confidence is the mean pLDDT over enzyme Cα atoms within each member. The interface confidence metrics are AF3 ipTM and the arithmetic mean of the two directional chain-pair minimum predicted aligned errors (PAEs) between the enzyme and PCNA. The mean clash indicator is the fraction of models for which AF3 reports a clash. Catalytic-motif RMSD uses an independent fit to the corresponding sequence-design structure, while enzyme Cα RMSD uses a separate fit of the enzyme Cα atoms. Interatomic distances are measured directly within each predicted structure.

The current terminal selection protocol requires all ensemble-mean apo and PCNA-containing acyl criteria in Supplementary Table 9. These criteria assess average fold confidence, interface confidence and catalytic geometry; they are not limits on every individual prediction or on mean-plus-standard-deviation bounds. The configured filter does not impose a minimum number of valid ensemble members. Candidates passing these filters were evaluated in the holo state similar to the model peptide case to test whether the enzyme retained its fold and positioned the Ubiquitin tail near the catalytic cysteine without the acceptor (Supplementary Table 4).

**Supplementary Table 9 | Configured ensemble-mean acceptance criteria for confidence-dependent redesign.** All criteria are mandatory and apply to means across available AF3 ensemble members. Distances, RMSDs and PAE values are in Å.

| State | Metric | Threshold |
|---|---|---|
| Apo | Enzyme Cα pLDDT | ≥85 |
| Apo | Enzyme Cα RMSD | ≤1.5 |
| Acyl with PCNA | ipTM | ≥0.75 |
| Acyl with PCNA | Mean of directional chain-pair minimum PAEs | ≤3.0 |
| Acyl with PCNA | Fraction of models with an AF3-reported clash | ≤0.10 |
| Acyl with PCNA | Enzyme Cα pLDDT | ≥85 |
| Acyl with PCNA | Catalytic-motif RMSD | ≤0.9 |
| Acyl with PCNA | Cys Sγ–His Nδ1 | ≤4.0 |
| Acyl with PCNA | Lys164 Nζ–donor carbonyl C | ≤4.5 |
| Acyl with PCNA | His Nδ1–Lys164 Nζ | ≤4.5 |
| Acyl with PCNA | Donor carbonyl O–Cys N | ≤3.9 |

#### 2.2 Experimental methods

##### 2.2.1 General methods: plasmids and reagents

Codon optimised genes encoding de novo Ub-Peptide Ligases, Ub-PCNA ligases, SUMO-Peptide fusions, Ub-intein fusions, Ubch5c, human PCNA wt/K164R, yeast PCNA and linear di/tri/tetraUb were purchased as DNA Strings (Twist Bioscience) and cloned into the pBAD, pCool, pTXB1, pET17 or pET28 vectors via restriction cloning or via Golden Gate assembly (see Supplementary Table 10). The gene coding for UBA1 was acquired from (Addgene #63571). Amino acid sequences of all proteins are listed in Section 3. All chemical reagents were purchased from Sigma Aldrich, Senn, Acros Organics, or Fisher Scientific. All Reagents were used without further purification unless stated otherwise. Ni-NTA agarose (Cat. No. 17-5318-02), His MultiTrap plates (Cat. No. 28-4009-90) AcroPrep Advance 96-well filter plate, 10K MWCO (Cat. No. 8034) was purchased from Cytiva. Bolt 4-12 % Bis-Tris gradient gels (Invitrogen) gels were run (at 165 V for 45 min) using a Bolt™ Mini Gel Tank system (Invitrogen) using a MES running buffer (50 mM, pH 6, 50 mM Tris, 0.1% (w/v) SDS, 1 mM EDTA). Gels were stained with Quick Coomassie Stain (Generon). PageRuler Prestained Plus Protein Ladder 10-250 kDa (ThermoFisher) was used as the protein marker. Protein and DNA concentrations were measured on a NanoPhotometer® NP60 (Implen).

**Supplementary Table 10 | Plasmids**

| Plasmid | Description |
|---|---|
| pET17_Ub_wt | Tagless human Ub under an T7 promotor (ampicillin resistance). |
| pTXB1_Ub_MxeGyrA_CBD | Human Ub fused to MxeGyrA intein followed by a chitin binding domain (CBD) (ampicillin resistance). |
| pFastBac1_H6-UBA1 | N-terminally H6-tagged UBA1 in FastBac backbone for Bacmid generation (ampicillin resistance). |
| pBAD_H6_TEV_linear_di/tri/tetraUb(G76V) | N-terminally H6-TEV tagged linear di/tri/tetraubiquitin with G76V mutations between the proximal Ubs under an arabinose promotor (ampicillin resistance) |
| pBAD_H6_SUMO_Peptide wt/K10A | N-terminally H6-tagged SUMO (SMT3) Peptide fusions (wt and K10A) under an arabinose promotor (ampicillin resistance) |
| pBAD_hPCNA_wt/K164R_H6 | C-terminally H6-tagged human PCNA (wt and K164R) under an arabinose promotor (ampicillin resistance) |
| pBAD_yPCNA_wt_H6 | C-terminally H6-tagged yeast PCNA under an arabinose promotor (ampicillin resistance) |
| pET28a_Ubch5c_S22R | Tagless human Ubch5c with a S22R mutation under a T7 promotor (kanamycin resistance) |
| pCool_Ub-Peptide-Ligases_H6 | C-terminally H6-tagged Ub-peptide Ligases 1-48 and mutants under a T7 promotor (kanamycin resistance) |

| | |
|---|---|
| pET28a_Ub-PCNA-Ligases_H6 | C-terminally H6-tagged Ub-PCNA Ligases 1-48 and mutants under a T7 promotor (kanamycin resistance) |

##### 2.2.2 Protein expression and purification

###### 2.2.2.1 Generation of Ub-MESNa by intein thiolysis

Chemically competent *E. coli* BL21(DE3) were transformed with the Ub(1-76)-intein-CBD (chitin-binding domain) fusion plasmid (see Supplementary Table 10). After recovery with 1 mL of SOC medium for 1 h at 37 °C, the cells were cultured overnight in 50 mL of 2x YT medium containing ampicillin (100 µg/mL) and 1 % glucose at 37 °C, 200 rpm. The overnight culture was diluted into 1.5 L of fresh 2x YT medium supplemented with ampicillin (50 µg/mL) in a 5 L flask and cultured at 37 °C with shaking (200 rpm) until $OD_{600}$ reached 0.6. The culture was cooled to 16 °C, IPTG (isopropyl beta-D-1-thiogalactopyranoside) was added to a final concentration of 0.5 mM and protein expression was induced overnight. The cells were harvested by centrifugation (4,000 x g, 40 min, 4 °C) and resuspended in 50 mL of lysis buffer (20 mM HEPES pH 7.5, 300 mM NaCl, 1 mM EDTA (ethylenediaminetetraacetic acid)) containing one tablet of protease inhibitor cocktail (Roche). The cell suspension was lysed by high pressure cell disruption using a Constant Systems CF1 (2x 30,000 psi) followed by centrifugation (14,000 x g, 40 min, 4 °C). The cleared lysate was applied to 2 x 5 mL of chitin resin (NEB) followed by incubation at 4 °C for 1 h. After washing with 20 column volums (CV) lysis buffer, thiolysis was initiated by addition of 10 mL of lysis buffer containing 100 mM MESNa (sodium 2-mercaptoethanesulfonate) followed by incubation at 4 °C for 24 h. Afterwards the flow-through was collected and concentrated using Amicon centrifugal filter units with a 3 kDa MWCO (molecular weight cut-off) (Millipore) and further purified via size-exclusion chromatography (SEC) using a Superdex 75 16/600 (GE Healthcare) with a buffer containing 20 mM HEPES pH 6.8, 150 mM NaCl. Fractions containing Ub-MESNa were pooled and concentrated. Protein concentration was determined by bicinchoninic acid assay (BCA) (Thermo Scientific) and Bradford (Sigma-Aldrich) assays. Liquid chromatography-mass spectrometry (LC-MS) analysis of the pooled material showed approximately one third of hydrolysed Ub alongside the thioester. Purified Ub-MESNa was flash frozen using liquid nitrogen and stored at -80 °C until further use.

###### 2.2.2.2 Expression and purification of Ub

Chemically competent *E. coli* Rosetta2 (DE3) were transformed with pET17b-Ub plasmid (see Supplementary Table 10). After recovery with 1 mL of SOC medium for 1 h at 37 °C, the cells were cultured overnight in 50 mL of 2x YT medium containing ampicillin (100 µg/mL), chloramphenicol (50 µg/mL) and 1 % glucose at 37 °C, 200 rpm. The overnight culture was diluted to an $OD_{600}$ of 0.05 in 3 L of fresh 2x YT medium supplemented with ampicillin

(50 µg/mL) and chloramphenicol (25 µg/mL) and cultured at 37 °C with shaking (200 rpm) until $OD_{600}$ reached 0.8 - 1.0. IPTG was added to a final concentration of 1 mM and protein expression was induced for 3 h at 37 °C. The cells were harvested by centrifugation (4000 x *g*, 40 min, 4 °C) and resuspended in lysis buffer (50 mM Tris pH 7.6, 10 mM $MgCl_2$, 1 mM EDTA). The cell suspension was lysed by high pressure cell disruption using a Constant Systems CF1 (2x 30,000 psi) followed by centrifugation (14,000 x *g*40 min, 4°C). The cleared lysate was transferred into a glass beaker in an ice-water bath placed on a magnetic stirrer. Precipitation was performed with 35 % perchloric acid until pH 4.0 - 4.5 was reached. After 5 min incubation at 4 °C while stirring, the milky solution was centrifuged (14,000 x *g*, 40 min, 4 °C) and the supernatant was transferred into a dialysis tubing with a MWCO of 2 kDa (Roth). Dialysis was performed overnight at 4 °C with 50 mM ammonium acetate buffer pH 4.5. The dialysed solution was centrifuged (14,000 x *g*, 40 min, 4 °C), filtered and purified via a HiTrap SP FF 5 mL cation exchange chromatography (GE, gradient 0 - 1 M NaCl). Fractions that showed > 95% purity, as judged by sodium dodecyl sulfate-polyacrylamide gel electrophoresis (SDS-PAGE), were pooled, concentrated, and further purified via SEC using a Superdex Increase 75 16/600 (GE Healthcare) with a buffer containing 20 mM HEPES pH 7.5, 150 mM NaCl. Fractions containing pure Ub were pooled together and concentrated using Amicon® centrifugal filter units with a 3 kDa MWCO (Millipore). Protein concentration was determined by BCA (Thermo Scientific) and Bradford (Sigma-Aldrich) assays. Purified Ub was flash frozen using liquid nitrogen and stored at -80 °C until further use.

###### 2.2.2.3 Generation of Ub-MESNa by E1-mediated capture

For E1-mediated Ub-MESNa generation a 500 µL reaction containing 500 µM Ub and 1 µM UBA1 was set up in 50 mM HEPES pH 7.5, 5 mM ATP, 10 mM $MgCl_2$ and 20 mM MESNa and incubated overnight at 37 °C. Conversion was followed by LC-MS and was complete after 16 h, with Ub-MESNa as the only detectable species. The reaction was purified by SEC on a Superdex 75 10/300 column equilibrated in 10 mM HEPES pH 6.8, 150 mM NaCl. Fractions containing pure Ub-MESNa were pooled, concentrated using Amicon centrifugal filter units with a 3 kDa MWCO (Millipore). Protein concentration was determined by BCA (Thermo Scientific) and Bradford (Sigma-Aldrich) assays. Purified Ub-MESNa was flash frozen using liquid nitrogen and stored at -80 °C until further use.

###### 2.2.2.4 Expression and purification of the SUMO–peptide fusions and the tagless peptides

Chemically competent *E. coli* K12 were transformed with pBAD-H6-SUMO-peptide plasmid (see Supplementary Table 10). After recovery with 1 mL of SOC medium for 1 h at 37 °C, the cells were cultured overnight in 50 mL of 2x YT medium containing ampicillin (100 µg/mL) at 37 °C, 200 rpm. The overnight culture was diluted into 1.5 L of fresh 2x YT medium supplemented with ampicillin (50 µg/mL) in a 5 L flask and cultured at 37 °C with shaking (200 rpm) until $OD_{600}$ reached 0.6. The culture was cooled to 16 °C, arabinose was added to a final concentration of 0.02 (w/v) and protein expression was induced overnight.

The cells were harvested by centrifugation (4000 x g, 10 min, 4 °C) and resuspended in lysis buffer (20 mM Tris pH 8.0, 300 mM NaCl, 30 mM Imidazole, 1 mM PMSF). The cell suspension was lysed by high pressure cell disruption using a Constant Systems CF1 (2x 30,000 psi) followed by centrifugation (14,000 x *g* 40 min, 4°C). The cleared lysate was added to 1 mL Ni-NTA slurry/1 L culture (Cytiva) equilibrated with wash buffer (20 mM Tris pH 8.0, 300 mM NaCl and 30 mM imidazole). Afterwards, the mixture was incubated with agitation for 1 h at 4 °C. After incubation, the mixture was transferred to an empty plastic column and washed with 10 CV of wash buffer. The protein was eluted with wash buffer supplemented with 300 mM imidazole pH 8.0. The fractions containing the SUMO-peptide fusion were pooled and concentrated using Amicon centrifugal filter units (Millipore) with a suitable MWCO followed by purification via SEC using a Superdex 75 16/600 (Cytiva) with SEC buffer (10 mM HEPES pH 7.5, 100 mM NaCl). Fractions containing the SUMO-peptide fusion were pooled and concentrated. Protein concentration was calculated from the measured A280 absorption (extinction coefficients were calculated with ProtParam (https://web.expasy.org/protparam/)). SUMO-peptide fusions were flash frozen using liquid nitrogen and stored at −80 °C until further use.

For accessing tagless peptides the NiNTA eluate was treated with SUMO protease (Sigma Aldrich) for 2h at room temperature followed by reverse NiNTA purification. The flow-through was concentrated using an Amicon centrifugal filter unit with a 3 kDa MWCO and purified on a Superdex 30 Increase 10/300 (Cytiva) equilibrated in 10 mM HEPES pH 7.5, 100 mM NaCl. Fractions containing the tagless peptide were pooled and concentrated using an Amicon centrifugal filter unit with a 3 kDa MWCO. Protein concentration was calculated from the measured A280 absorption and by BCA (Thermo Scientific) and Bradford (Sigma-Aldrich) assays. SUMO-peptide fusions were flash frozen using liquid nitrogen and stored at −80 °C until further use.

###### 2.2.2.5 Expression and purification of de novo designed ligases in plate format

Genes encoding de novo designed C-terminally H6-tagged Ub-Peptide or Ub-PCNA Ligases were either cloned via Golden Gate assembly into a pCool vector (see Supplementary Table 10) or ordered from Twist Biotech in a pET29 vector. Chemically competent *E. coli* BL21(DE3) were transformed with pCool/pET29 vectors encoding the de novo ligases followed by recovery with 1 mL of SOC medium for 1 h at 37 °C. Afterwards the cells were collected by centrifugation (4,000 x g, 2 min), resuspended in 100 µL 2x YT and transferred into 3 mL of 2x YT medium containing lactose/glucose/glycerol autoinduction mix and kanamycin (25 µg/mL) in a 24-well deep-well plate followed by incubation overnight at 37 °C while shaking (200 rpm). Cultures were harvested (4,000 x g, 2 min, 4 °C) and the pellets were resuspended in 300 µL of lysis buffer (20 mM Tris pH 8.0, 300 mM NaCl and 30 mM imidazole) and subjected to two freeze-thaw cycles in liquid nitrogen followed by lysis via sonication in a Bioruptor (Diagenode) (2 x 10 min). The lysates were cleared by centrifugation (21,000 x g, 15 min, 4 °C) and applied to the wells of a 96-well Ni-NTA (nickel-nitrilotriacetic acid) filter plate (Cytiva) mounted on a vacuum manifold, which had been equilibrated with lysis buffer. The wells were washed three times with 700 µL of lysis buffer and the ligases were eluted in 3 x 100 µL of lysis buffer containing 300 mM imidazole. The eluates were concentrated and rebuffered in a 96-well filter plate with a 10 kDa MWCO (Cytiva) on the same vacuum manifold, exchanging the buffer three times into 20 mM Hepes pH 7.5, 150 mM NaCl. The concentrated ligases were recovered from the plate in a final volume of 50 - 100 µL per design, transferred into 1.5 mL reaction tubes and stored at 4 °C until further use. Purity was assessed by SDS-PAGE.

###### 2.2.2.6 Expression and purification of de novo designed ligases in shaker flasks

Chemically competent *E. coli* BL21(DE3) were transformed with pCool/pET29 plasmids encoding C-terminally H6-tagged de novo designed ligases (see Supplementary Table 10). After recovery with 1 mL of SOC medium for 1 h at 37 °C, the cells were cultured overnight in 50 mL of 2x YT medium containing kanamycin (50 µg/mL) and 1 % glucose at 37 °C, 200 rpm. The overnight culture was diluted into 1.5 L of fresh 2x YT medium supplemented with kanamycin (25 µg/mL) in a 5 L flask and cultured at 37 °C with shaking (200 rpm) until $OD_{600}$ reached 0.6. The culture was cooled to 16 °C, IPTG was added to a final concentration of 0.5 mM and protein expression was induced overnight. The cells were harvested by centrifugation (4,000 x g, 40 min, 4 °C) and resuspended in 50 mL of lysis buffer (20 mM Tris pH 8.0, 300 mM NaCl, 30 mM Imidazole, 1 mM PMSF) containing one tablet of protease inhibitor cocktail (Roche). The cell suspension was lysed by high pressure cell disruption using a Constant Systems CF1 (2x 30,000 psi) followed by centrifugation (14,000 x g, 40 min, 4 °C). The cleared lysate was applied to 1 mL Ni-NTA slurry/1 L culture (Cytiva) equilibrated with wash buffer (20 mM Tris pH 8.0, 300 mM NaCl and 30 mM imidazole). Afterwards, the mixture was incubated with agitation for 1 h at 4 °C. After incubation, the mixture was transferred to an empty plastic column and washed with 10 CV of wash buffer. The protein was eluated in 1 mL fractions with wash buffer supplemented with 300 mM imidazole pH 8.0. The fractions containing the de novo designed ligases were pooled and concentrated using Amicon centrifugal filter units (Millipore) with a suitable molecular weight cutoff (MWCO) followed by SEC using a Superdex 75 16/600 (Cytiva) with SEC buffer (10 mM HEPES pH 7.5, 100 mM NaCl). Fractions containing the de novo designed ligases were pooled and concentrated. Protein concentration was calculated from the measured A280 absorption. De novo ligases were flash frozen using liquid nitrogen and stored at −80 °C until further use.

###### 2.2.2.7 Expression and purification of human PCNA and yeast PCNA

Chemically competent *E. coli* K12 were transformed with pBAD-PCNA (encoding for either C-terminally H6-tagged human PCNA wt/K164R or yeast PCNA wt, see Supplementary Table 10). After recovery with 1 mL of SOC medium for 1 h at 37 °C, the cells were cultured overnight in 50 mL of 2x YT medium containing ampicillin (100 µg/mL) at 37 °C, 200 rpm. The overnight culture was diluted to an $OD_{600}$ of 0.05 in 200 mL of fresh 2x YT medium supplemented with ampicillin (50 µg/mL) and cultured at 37 °C with shaking (200 rpm) until $OD_{600}$ = 0.6. Arabinose was added to a final concentration of 0.02 % (w/v) and protein expression was induced for 3 h at 37 °C. The cells were harvested by centrifugation (4000 x g, 40 min, 4 °C) and resuspended in lysis buffer (20 mM Tris pH 8.0, 300 mM NaCl, 30 mM Imidazole, 1 mM PMSF). Afterwards, the cell suspension was incubated on ice for 30 min and sonicated with cooling in an ice-water bath. The lysed cells were centrifuged (14,000 x g, 20 min, 4 °C) and the clear lysate was added to 1 mL Ni-NTA slurry/1 L culture (Cytiva) equilibrated with wash buffer (20 mM Tris pH 8.0, 300 mM NaCl and 30 mM imidazole). Afterwards, the mixture was incubated with agitation for 1 h at 4 °C. After incubation, the mixture was transferred to an empty plastic column and washed with 10 CV of wash buffer. The protein was eluated in 1 mL fractions with wash buffer supplemented with 300 mM imidazole pH 8.0. The fractions containing PCNA variants were pooled and concentrated using Amicon centrifugal filter units (Millipore) with a 10 kDa MWCO followed by SEC using a Superdex 200 10/300 (Cytiva) with SEC buffer (25 mM HEPES pH 7.5, 150 mM NaCl). Fractions containing PCNA variants were pooled and concentrated. Protein concentration was calculated from the measured A280 absorption. PCNA variants were flash frozen using liquid nitrogen and stored at −80 °C until further use.

###### 2.2.2.8 Expression and purification of linear di/tri/tetraUb

Chemically competent E. coli K12 were transformed with pBAD-di/tri/tetraUb (encoding for MP-H6-TEV-di/tri/tetraUb with G76V mutation(s) at the distal Ub moieties, see Supplementary Table 10). After recovery with 1 mL of SOC medium for 1 h at 37 °C, the cells were cultured overnight in 50 mL of 2x YT medium containing ampicillin (100 µg/mL) at 37 °C, 200 rpm. The overnight culture was diluted to an $OD_{600}$ of 0.05 in 200 mL of fresh 2x YT medium supplemented with ampicillin (50 µg/mL) and cultured at 37 °C with shaking (200 rpm) until $OD_{600}$ = 0.6. Arabinose was added to a final concentration of 0.02 % (w/v) and protein expression was induced for 3 h at 37 °C. The cells were harvested by centrifugation (4000 x g, 40 min, 4 °C) and resuspended in lysis buffer (20 mM Tris pH 8.0, 300 mM NaCl, 30 mM Imidazole, 1 mM PMSF). Afterwards, the cell suspension was incubated on ice for 30 min and sonicated with cooling in an ice-water bath. The lysed cells were centrifuged (14,000 x g, 20 min, 4 °C) and the clear lysate was added to 1 mL Ni-NTA slurry/1 L culture (Cytiva) equilibrated with wash buffer (20 mM Tris pH 8.0, 300 mM NaCl and 30 mM imidazole). Afterwards, the mixture was incubated with agitation for 1 h at 4 °C. After incubation, the mixture was transferred to an empty plastic column and washed with 10 CV of wash buffer. The protein was eluated in 1 mL fractions with wash buffer supplemented with 300 mM imidazole pH 8.0. The fractions containing the di/tri/tetraUb were pooled and concentrated using Amicon centrifugal filter units

(Millipore) with a suitable molecular weight cutoff (MWCO) followed by SEC using a Superdex 75 10/300 (Cytiva) with SEC buffer (25 mM Tris pH 7.5, 100 mM NaCl). Fractions containing di/tri/tetraUb variants were pooled and concentrated. Protein concentration was calculated from the measured A280 absorption. Di/tri/tetraUb were flash frozen using liquid nitrogen and stored at −80 °C until further use.

**2.2.2.9 Expression and purification of Ubch5c and H6-UBA1**

Ubch5c was expressed and purified from *E. coli* as previously described. UBA1 was expressed in Insect cells and purified and previously described.

**2.2.3 In vitro de novo Ub ligase assays**

**2.2.3.1 LC-MS based screening assay for de novo Ub-peptide ligases**

Ub-peptide ligase activity was assayed by following the transfer of Ub from Ub-MESNa onto the acceptor peptide. A 2x master mix containing 250 µM peptide and 100 µM Ub-MESNa was prepared in 10 mM HEPES pH 7.5, 150 mM NaCl and mixed 1:1 (v/v) with the crude ligase eluate from the plate-format purification (Section 2.2.2.5), giving final concentrations of 125 µM peptide, 50 µM Ub-MESNa and 50 % (v/v) ligase eluate. Reactions were incubated at 37 °C for 30 h. At the end point, 1 µL of the reaction was diluted with 9 µL of 1% (v/v) formic acid and analysed by ESI-LC-MS as described in Section 2.2.4. Conversion was determined from the deconvoluted mass spectra by comparing the intensity of the Ub-peptide conjugate with that of the remaining unmodified peptide, with hydrolysed Ub and residual Ub-MESNa monitored in the same spectrum.

**2.2.3.2 LC-MS based kinetic characterisation of Ub-peptide ligases**

The twelve most active designs from the crude screen and the catalytic tetrad mutants of Ligase 29 were re-expressed, purified to homogeneity by SEC and assayed under defined conditions. Reactions contained 200 µM peptide, 100 µM Ub-MESNa and 20 µM ligase in 10 mM HEPES pH 7.5, 150 mM NaCl and were incubated at 37 °C. At denoted time points, 1 µL of the reaction was diluted with 9 µL of 1% (v/v) formic acid and analysed by ESI-LC-MS as described in Section 2.2.4.

**2.2.3.3 LC-MS based thioester donor specificity assays**

To probe which Ub-thioester species the ligase can use as a donor, Ub-MESNa was converted in situ to the corresponding thioester before the reaction was started. Ub-MESNa (100 µM) was pre-incubated with 10 mM of the target thiol (BME (2-mercaptoethanol), DTT (dithiothreitol) or reduced glutathione (GSH)) for 5 min in 10 mM HEPES pH 7.5, 150 mM NaCl containing 200 µM peptide, after which Ligase 29 was added to 20 µM and the reaction was incubated at 37 °C.

At denoted time points, 1 µL of the reaction was diluted with 9 µL of 1% (v/v) formic acid and analysed by ESI-LC-MS as described in Section 2.2.4.

###### 2.2.3.4 In vitro ubiquitylation of SUMO-peptide fusion with Ub-peptide ligase 29

To follow ubiquitylation by gel electrophoresis, the uncleaved H6-SUMO-peptide fusion was used as the acceptor in place of the free peptide. Reactions contained 200 µM SUMO-peptide, 100 µM Ub-MESNa and 20 µM Ligase 29 in 10 mM HEPES pH 7.5, 150 mM NaCl and were incubated at 37 °C. At the time points indicated, 4 µL samples were mixed with 8 µL of 4× SDS sample buffer and heated for 2 min at 95 °C and reaction progress was analysed by SDS-PAGE and Coomassie staining.

###### 2.2.3.5 SDS-PAGE based screening assay for de novo Ub-PCNA ligases activity

The 48 designed ligases were screened for transfer of Ub onto PCNA. A 2x master mix containing 200 µM Ub-MESNa and 100 µM PCNA was prepared in 20 mM HEPES pH 7.5, 150 mM NaCl and mixed 1:1 (v/v) with each Ni-NTA-purified, rebuffered ligase, giving final concentrations of 100 µM Ub-MESNa, 50 µM PCNA and 50 % (v/v) ligase eluate. Reactions were incubated at 37 °C for 20 h. For analysis, 1 µL of each reaction was mixed with 5 µL of 4x SDS sample buffer, heated at 95 °C and resolved by SDS-PAGE followed by Coomassie staining.

The four designs identified as active in the screen were re-assayed in parallel on PCNA wt and on the acceptor-site variant PCNA K164R. Reactions contained 200 µM PCNA (wt or K164R), 600 µM Ub-MESNa and 50 % (v/v) rebuffered ligase in 20 mM HEPES pH 7.5, 150 mM NaCl and were incubated at 37 °C. Samples were taken at 1 and 3 h and analysed via SDS-PAGE as described above.

###### 2.2.3.6 In vitro PCNA ubiquitylation assays with Ub-PCNA ligase 4 and its variants

All assays were performed with SEC-purified Ligase 4 and its variants. Reactions were assembled in 10 mM HEPES pH 7.5, 100 mM NaCl and incubated at 37 °C; PCNA, Ub-MESNa and ligase concentrations for each experiment are listed in the corresponding figures. At the indicated times, samples were taken, mixed with 4x SDS sample buffer, heated at 95 °C and resolved by SDS-PAGE followed by Coomassie staining.

###### 2.2.3.7 E1-coupled in vitro (poly)ubiquitylation of PCNA

For the E1-coupled cascade the Ub thioester donor was generated in situ rather than supplied as pre-formed Ub-MESNa. Reactions contained 100 µM PCNA wt, 200 µM of the respective mono- or linear di-, tri- or tetraubiquitin, 10 µM Ligase 4 N148A, 8 µM Uba1, 5 mM ATP, 6 mM $MgCl_2$ and 5 mM MESNa in 10 mM HEPES pH 7.5, 100 mM NaCl, and were incubated at 37 °C. For analysis, 1 µL of each reaction was mixed with 8 µL of 4x SDS sample buffer, heated at 95 °C and resolved by SDS-PAGE followed by Coomassie staining.

##### 2.2.3.8 Ubch5c-mediated ubiquitylation of PCNA

For Ubch5c-mediated PCNA ubiquitylation, reactions contained 100 µM PCNA wt, 200 µM Ub, 10 µM Ubch5c, 8 µM Uba1, 5 mM ATP and 6 mM $MgCl_2$ in 10 mM HEPES pH 7.5, 100 mM NaCl, and were incubated at 37 °C. For analysis, 1 µL of each reaction was mixed with 8 µL of 4x SDS sample buffer, heated at 95 °C and resolved by SDS-PAGE followed by Coomassie staining.

##### 2.2.4 Intact protein mass spectrometry

ESI-LC-MS analysis was performed on an Agilent 1260 Infinity Series LC system with an Agilent 6210 ESI Single Quadrupole mass spectrometer using a Jupiter C4 column (2 mm, 150 mm, 300 Å, 5 µm) capillary column (Phenomenex, Torrence, USA). The analysis was performed at RT with a flow rate of 900 µL/min and a gradient of 10% - 55% solvent B in 1.65 min, followed by a gradient of 55 %-90 % solvent B in 0.85 min (solvent A: 0.1 % formic acid (FA) in water, solvent B 0.1 % FA in acetonitrile). Data generated on the Agilent LC-MS was analysed with OpenLab ChemStation (Agilent).

##### 2.2.5 X-ray crystallography

**Crystallisation and Structure Determination of PL4 N148A (PL4.1).** Purified PL4 N148A was concentrated to 100 mg $mL^{-1}$ for crystallisation. Crystals were grown at 293 K by sitting-drop vapour diffusion using drops containing 0.2 µL protein solution and 0.1 µL reservoir solution equilibrated against 0.1 M citric acid, pH 4.0, and 0.8 M ammonium sulfate. Prior to data collection, crystals were transferred to reservoir solution supplemented with 30% (v/v) glycerol and flash-cooled in liquid nitrogen. X-ray diffraction data were collected at 100 K at beamline X06DA (PXIII) of the Swiss Light Source (Paul Scherrer Institute, Villigen, Switzerland) using a DECTRIS PILATUS4 2M detector ($\lambda$ = 0.9537 Å) and processed with XDS[5]. Initial phases were obtained by molecular replacement with PHASER[6] using an AlphaFold-predicted model[3] of PL4 N148A as the search model. The crystals belong to space group $P3_221$ and contain one molecule per asymmetric unit. The structure was completed by iterative manual model building in COOT[7] and refinement with PHENIX[8] (phenix.refine) using two TLS groups (residues 0-37 and 38-202) and riding hydrogen atoms; refinement weights were optimised in the final cycles. Water molecules were placed in PHENIX and inspected manually in COOT. Residues are numbered with the initiator methionine as Met0. Residues 148-151, including the mutated position Ala148, as well as the C-terminal linker and $His_6$ tag (GGSHHHHHH) were not defined by electron density. One sulfate ion from the crystallisation solution was bound in the active site. Model geometry and agreement with the diffraction data were monitored throughout refinement, and the final model was validated with MolProbity[9].

Data collection and refinement statistics are summarised in Supplementary Table 11. Coordinates and structure factors have been deposited in the Protein Data Bank under accession code 35ND.

**Supplementary Table 11 | X-ray data collection and refinement statistics**

| | **PL4 N148A (PL4.1)** |
|---|---|
| **Crystal parameters** | |
| Space group | $P3_221$ |
| Cell constants | A = b = 98.6 Å, c = 44.5 Å |
| Subunits per AU[a] | 1 |
| **Data collection** | |
| Beamline | X06DA, SLS |
| Wavelength (Å) | 0.9537 |
| Resolution range (Å)[b] | 28.45-1.60 (1.70-1.60) |
| No. observations | 183691 |
| No. unique reflections[c] | 32962 |
| Completeness (%)[b] | 99.7 (99.6) |
| Redundancy | 5.6 (5.3) |
| $R_{merge}$ (%)[b, d] | 4.6 (78.3) |
| CC½ (%) | 100.0 (82.3) |
| I/σ(I)[b] | 20.1 (2.4) |
| **Refinement** | |
| Resolution range (Å) | 28.45-1.60 |
| No. refl. working set | 31287 |
| No. refl. test set | 1647 |
| No. non-H-atoms | 1720 |
| No. ligand atoms | 5 |
| No. solvent atoms | 171 |
| $R_{work}/R_{free}$ (%)[e] | 17.1 / 18.5 |
| r.m.s.d. bond (Å) / (°)[f] | 0.010 / 1.0 |

| | |
|---|---|
| Average B-factor (Å$^2$) | 32.6 |
| Ramachandran plot (%)[g] | 99.5 / 0.5 / 0 |
| PDB accession code | **35ND** |

[a] Asymmetric unit; each dataset was collected from a single crystal.
[b] The values in parentheses for resolution range, completeness, $R_{merge}$, CC½, and I/σ(I) correspond to the highest-resolution shell.
[c] Data were processed with XDS from a single crystal. Friedel pairs were treated as equivalent reflections.
[d] $R_{merge}(I) = \Sigma_{hkl}\Sigma_j \mid I(hkl)_j - \langle I(hkl)\rangle \mid / \Sigma_{hkl} \Sigma_j I(hkl)_j$, where $I(hkl)_j$ is the j$^{th}$ measurement of the intensity of reflection hkl and $\langle I(hkl)\rangle$ is the average intensity
[e] $R = \Sigma_{hkl} \mid |F_{obs}| - |F_{calc}| \mid / \Sigma_{hkl} |Fobs|$, where $R_{free}$ is calculated without a sigma cut off for a randomly chosen 5% of reflections, which were not used for structure refinement, and $R_{work}$ is calculated for the remaining reflections
[f] Deviations from ideal bond lengths/angles
[g] Percentage of residues in favoured region / allowed region / outlier region

### 3 Amino acid sequences of proteins

#### 3.1 General proteins

Ubiquitin
MQIFVKTLTGKTITLEVEPSDTIENVKAKIQDKEGIPPDQQRLIFAGKQLEDGRTLSDYNIQKESTLHLVLRLR
GG

Ubiquitin-MxeGyrA-CBD fusion
MQIFVKTLTGKTITLEVEPSDTIENVKAKIQDKEGIPPDQQRLIFAGKQLEDGRTLSDYNIQKESTLHLVLRLR
GGCITGDALVALPEGESVRIADIVPGARPNSDNAIDLKVLDRHGNPVLADRLFHSGEHPVYTVRTVEGLRVT
GTANHPLLCLVDVAGVPTLLWKLIDEIKPGDYAVIQRSAFSVDCAGFARGKPEFAPTTYTVGVPGLVRFLEA
HHRDPDAQAIADELTDGRFYYAKVASVTDAGVQPVYSLRVDTADHAFITNGFVSHATGLTGLNSGLTTNPG
VSAWQVNTAYTAGQLVTYNGKTYKCLQPHTSLAGWEPSNVPALWQLQ

H6-UBA1
MHHHHHHAKNGSEADIDEGLYSRQLYVLGHEAMKRLQTSSVLVSGLRGLGVEIAKNIILGGVKAVTLHDQ
GTAQWADLSSQFYLREEDIGKNRAEVSQPRLAELNSYVPVTAYTGPLVEDFLSGFQVVVLTNTPLEDQLRVG
EFCHNRGIKLVVADTRGLFGQLFCDFGEEMILTDSNGEQPLSAMVSMVTKDNPGVVTCLDEARHGFESGDF
VSFSEVQGMVELNGNQPMEIKVLGPYTFSICDTSNFSDYIRGGIVSQVKVPKKISFKSLVASLAEPDFVVTDF
AKFSRPAQLHIGFQALHQFCAQHGRPPRPRNEEDAAELVALAQAVNARALPAVQQNNLDEDLIRKLAYVAT
GDLAPINAFIGGLAAQEVMKACSGKFMPIMQWLYFDALECLPVDKEVLTEDKCLQRQNRYDGQVAVFGSD
LQEKLGKQKYFLVGAGAIGCELLKNFAMIGLGCGEGGEIIVTDMDTIEKSNLNRQFLFRPWDVTKLKSDTA
AAAVRQMNPHIRVTSHQNRVGPDTERIYDDDFFQNLDGVANALDNVDARMYMDRRCVYYRKPLLESGTL
GTKGNVQVVIPFLTESYSSSQDPPEKSIPICTLKNFPNAIEHTLQWARDEFEGLFKQPAENVNQYLTDPKFVE
RTLRLAGTQPLEVLEAVQRSLVLQRPQTWADCVTWACHHWHTQYSNNIRQLLHNFPPDQLTSSGAPFWSGP
KRCPHPLTFDVNNPLHLDYVMAAANLFAQTYGLTGSQDRAAVATFLQSVQVPEFTPKSGVKIHVSDQELQS
ANASVDDSRLEELKATLPSPDKLPGFKMYPIDFEKDDDSNFHMDFIVAASNLRAENYDIPSADRHKSKLIAG
KIIPAIATTTAAVVGLVCLELYKVVQGHRQLDSYKNGFLNLALPFFGFSEPLAAPRHQYYNQEWTLWDRFEV
QGLQPNGEEMTLKQFLDYFKTEHKLEITMLSQGVSMLYSFFMPAAKLKERLDQPMTEIVSRVSKRKLGRHV
RALVLELCCNDESGEDVEVPYVRYTIR

H6-TEV-linear-diUb(G76V)
MPHHHHHHENLYFQGGSMQIFVKTLTGKTITLEVEPSDTIENVKAKIQDKEGIPPDQQRLIFAGKQLEDGRTL
SDYNIQKESTLHLVLRLRGVMQIFVKTLTGKTITLEVEPSDTIENVKAKIQDKEGIPPDQQRLIFAGKQLEDG
RTLSDYNIQKESTLHLVLRLRGG

H6-TEV-linear-triUb(G76V)
MPHHHHHHENLYFQGGSMQIFVKTLTGKTITLEVEPSDTIENVKAKIQDKEGIPPDQQRLIFAGKQLEDGRTL
SDYNIQKESTLHLVLRLRGVMQIFVKTLTGKTITLEVEPSDTIENVKAKIQDKEGIPPDQQRLIFAGKQLEDG
RTLSDYNIQKESTLHLVLRLRGVMQIFVKTLTGKTITLEVEPSDTIENVKAKIQDKEGIPPDQQRLIFAGKQLE
DGRTLSDYNIQKESTLHLVLRLRGG

H6-TEV-linear-tetraUb(G76V)
MPHHHHHHENLYFQGGSMQIFVKTLTGKTITLEVEPSDTIENVKAKIQDKEGIPPDQQRLIFAGKQLEDGRTL
SDYNIQKESTLHLVLRLRGVMQIFVKTLTGKTITLEVEPSDTIENVKAKIQDKEGIPPDQQRLIFAGKQLEDG
RTLSDYNIQKESTLHLVLRLRGVMQIFVKTLTGKTITLEVEPSDTIENVKAKIQDKEGIPPDQQRLIFAGKQLE
DGRTLSDYNIQKESTLHLVLRLRGVMQIFVKTLTGKTITLEVEPSDTIENVKAKIQDKEGIPPDQQRLIFAGKQ
LEDGRTLSDYNIQKESTLHLVLRLRGG

H6-SUMO-Peptide wt
MPGSHHHHHHGSMDSEVNQEAKPEVKPEVKPETHINLKVSDGSSEIFFKIKKTTPLRRLMEAFAKRQGKEM
DSLRFLYDGIRIQADQTPEDLDMEDNDIIEAHREQIGGSIAEAFREAKEYLLAAAQQ

H6-SUMO-Peptide K10A
MPGSHHHHHHGSMDSEVNQEAKPEVKPEVKPETHINLKVSDGSSEIFFKIKKTTPLRRLMEAFAKRQGKEM
DSLRFLYDGIRIQADQTPEDLDMEDNDIIEAHREQIGGSIAEAFREAAEYLLAAAQQ

PCNA-H6 wt
MFEARLVQGSILKKVLEALKDLINEACWDISSSGVNLQSMDSSHVSLVQLTLRSEGFDTYRCDRNLAMGVN
LTSMSKILKCAGNEDIITLRAEDNADTLALVFEAPNQEKVSDYEMKLMDLDVEQLGIPEQEYSCVVKMPSG
EFARICRDLSHIGDAVVISCAKDGVKFSASGELGNGNIKLSQTSNVDKEEEAVTIEMNEPVQLTFALRYLNFF
TKATPLSSTVTLSMSADVPLVVEYKIADMGHLKYYLAPKIEDEEGSHHHHHH

PCNA-H6 K164R
MFEARLVQGSILKKVLEALKDLINEACWDISSSGVNLQSMDSSHVSLVQLTLRSEGFDTYRCDRNLAMGVN
LTSMSKILKCAGNEDIITLRAEDNADTLALVFEAPNQEKVSDYEMKLMDLDVEQLGIPEQEYSCVVKMPSG
EFARICRDLSHIGDAVVISCARDGVKFSASGELGNGNIKLSQTSNVDKEEEAVTIEMNEPVQLTFALRYLNFF
TKATPLSSTVTLSMSADVPLVVEYKIADMGHLKYYLAPKIEDEEGSHHHHHH

Ubch5c S22R
MALKRINKELSDLARDPPAQCRAGPVGDDMFHWQATIMGPNDSPYQGGVFFLTIHFPTDYPFKPPKVAFTT
RIYHPNINSNGSICLDILRSQWSPALTISKVLLSICSLLCDPNPDDPLVPEIARIYKTDRDKYNRISREWTQKYA
M

#### 3.2 De novo ubiquitin ligases

***Note:*** *Residue numbering refers to the designed sequence and excludes the N-terminal residues contributed by the expression construct (Met for the Ub-PCNA ligases, Met-Ser-Gly for the Ub-peptide ligases).*

**Ub-Peptide Ligases**

Ub-Peptide Ligase 29 **wt** H6
MSGITPEEANARAREIAELYAAGRRDEAARRAAELALELLDQYLKEKYGVELTDREVAWEPLPDGRRLPYG
RARLSREDAEKVLVEPGYPHAYEGGRLAGAFGIFTPDGKPVHVVALYEEPDGTVEVLDFLPAQVYYGFIGG
CLAAAAYVLVKRGFPEAARRVVEEGGGVADEKARDFFRDLIRALGHPDIGDLAAERLAGSGSHHWGSTHH
HHHH

Ub-Peptide Ligase 29 **C140A** H6
MSGITPEEANARAREIAELYAAGRRDEAARRAAELALELLDQYLKEKYGVELTDREVAWEPLPDGRRLPYG
RARLSREDAEKVLVEPGYPHAYEGGRLAGAFGIFTPDGKPVHVVALYEEPDGTVEVLDFLPAQVYYGFIGG
ALAAAAYVLVKRGFPEAARRVVEEGGGVADEKARDFFRDLIRALGHPDIGDLAAERLAGSGSHHWGSTHH
HHHH

Ub-Peptide Ligase 29 **D126A** H6
MSGITPEEANARAREIAELYAAGRRDEAARRAAELALELLDQYLKEKYGVELTDREVAWEPLPDGRRLPYG
RARLSREDAEKVLVEPGYPHAYEGGRLAGAFGIFTPDGKPVHVVALYEEPDGTVEVLAFLPAQVYYGFIGG
CLAAAAYVLVKRGFPEAARRVVEEGGGVADEKARDFFRDLIRALGHPDIGDLAAERLAGSGSHHWGSTHH
HHHH

Ub-Peptide Ligase 29 **H110A** H6
MSGITPEEANARAREIAELYAAGRRDEAARRAAELALELLDQYLKEKYGVELTDREVAWEPLPDGRRLPYG
RARLSREDAEKVLVEPGYPHAYEGGRLAGAFGIFTPDGKPVAVVALYEEPDGTVEVLDFLPAQVYYGFIGGC
LAAAAYVLVKRGFPEAARRVVEEGGGVADEKARDFFRDLIRALGHPDIGDLAAERLAGSGSHHWGSTHHH
HHH

Ub-Peptide Ligase 29 **Q131A** H6
MSGITPEEANARAREIAELYAAGRRDEAARRAAELALELLDQYLKEKYGVELTDREVAWEPLPDGRRLPYG
RARLSREDAEKVLVEPGYPHAYEGGRLAGAFGIFTPDGKPVHVVALYEEPDGTVEVLDFLPAAVYYGFIGGC
LAAAAYVLVKRGFPEAARRVVEEGGGVADEKARDFFRDLIRALGHPDIGDLAAERLAGSGSHHWGSTHHH
HHH

**Ub-PCNA Ligases**

Ub-PCNA Ligase 4 **wt** H6
MIEEEIREWLRELGVDEAAIEKLLEAARKAQAALDANDVAGFRAALEELKAAIKEAAATAREVTPEGEFRP
LPDGSVEIELRLPDGRKLTVVVDPKLGGYCAGFAVAILEALGRPGRIIPYPIPTPFGLRPHVAVAPGGDPEKVID
PGNYGGDVVDFDKIVDNIERKLREKGIELTPEQRARLRERIEALVAALKAAQAAARAGGSHHHHHH

Ub-PCNA Ligase 4 **C100A** H6
MIEEEIREWLRELGVDEAAIEKLLEAARKAQAALDANDVAGFRAALEELKAAIKEAAATAREVTPEGEFRP
LPDGSVEIELRLPDGRKLTVVVDPKLGGYAAGFAVAILEALGRPGRIIPYPIPTPFGLRPHVAVAPGGDPEKVID
PGNYGGDVVDFDKIVDNIERKLREKGIELTPEQRARLRERIEALVAALKAAQAAARAGGSHHHHHH

Ub-PCNA Ligase 4 **H131A** H6
MIEEEIREWLRELGVDEAAIEKLLEAARKAQAALDANDVAGFRAALEELKAAIKEAAATAREVTPEGEFRP
LPDGSVEIELRLPDGRKLTVVVDPKLGGYCAGFAVAILEALGRPGRIIPYPIPTPFGLRPAVAVAPGGDPEKVID
PGNYGGDVVDFDKIVDNIERKLREKGIELTPEQRARLRERIEALVAALKAAQAAARAGGSHHHHHH

Ub-PCNA Ligase 4 **D145A** H6
MIEEEIREWLRELGVDEAAIEKLLEAARKAQAALDANDVAGFRAALEELKAAIKEAAATAREVTPEGEFRP
LPDGSVEIELRLPDGRKLTVVVDPKLGGYCAGFAVAILEALGRPGRIIPYPIPTPFGLRPHVAVAPGGDPEKVIA
PGNYGGDVVDFDKIVDNIERKLREKGIELTPEQRARLRERIEALVAALKAAQAAARAGGSHHHHHH

Ub-PCNA Ligase 4 **N148A** H6
MIEEEIREWLRELGVDEAAIEKLLEAARKAQAALDANDVAGFRAALEELKAAIKEAAATAREVTPEGEFRP
LPDGSVEIELRLPDGRKLTVVVDPKLGGYCAGFAVAILEALGRPGRIIPYPIPTPFGLRPHVAVAPGGDPEKVID
PGAYGGDVVDFDKIVDNIERKLREKGIELTPEQRARLRERIEALVAALKAAQAAARAGGSHHHHHH

#### Ub-Peptide Ligases 1-48

***Note:*** *Residue numbering refers to the designed sequence and excludes the N-terminal residues contributed by the expression construct (Met for the Ub-PCNA ligases, Met-Ser-Gly for the Ub-peptide ligases).*

Ub-peptide ligase 1 H6
MSGLEVRYEGYEEFLEKAREEAAKATPEERERIEKGIEAYKAFKEAAIKAVKEALEAGKEEAVEPVGDGRVE
VRLVRLPSGAVVGLVALLDPEGRGVHVAAVVAEGDKAVILDPLPEQLERGLDGGCLPLALRAVREAAERLAT
PEEADAAVRRLAEGGGALAATREGAERAAEGLRELGFEARVRPDPERPGHWVAEVVPPGSGSHHWGSTHH
HHHH

Ub-peptide ligase 2 H6
MSGLEVEFEGKEEVLKAFEDAKKKATPEEKKKIDELKKAYEAALEAAREAVRKAYEEGKERAEAPLGDAK
VVAELVPLSSGNYLGIVQFWNPDGTGVHVLAVYVEGDKAVIWDPLAEQIKKKSDIGCVALAAKRLYEKGKE
LASEEEARAAIKRLLAGGGDVTPTREGAEEAERILRELGLDVTIVEDPDRPGEWEVRIEPVGSGSHHWGSTH
HHHHH

Ub-peptide ligase 3 H6
MSGLKIKIVGREEFEAAIDATLPKATPEEREAAEKGRAAWREYYEAAARAVREAVEAGREEARVAVGDATV
RARLVRLPDGSVVGVVEFLDRNGEGLHVFGVTVEGDEGVIHDPLPQQIREGRSAGCVGRALQFVAEFARAR
ASPEEAREAVARLARGGGDDAESREEAEWAAELFRELGFEAEVREDPERPGRWVARITRAGSGSHHWGSTH
HHHHH

Ub-peptide ligase 4 H6
MSGVEVEYRGEEEFEAIAEKIIAEASPEEAERLRAGLEGYRRFAEAARRAVAQALAEGRDEAEVPVGDARV
RVRLVRLSSGAVVGLVYFYDPDGRGLHAAAVAARGDRAEIIDPLPEQIRKGEDGACFPRALQAVREAARELA
TEEEADAAVRRLARGGGAIAPTEEGAERAKELWAELGLEAEVRPDPEQPGKWVAEVGEPGSGSHHWGSTH
HHHHH

Ub-peptide ligase 5 H6
MSGGGRGYAIVDPEVSREEAERWKARADAVAERLGELLGVPVALAIAVVLRDGRPEIAHVFAVDPEDPRRVI
DPLPFQLLAIHRGILKKNVGGCGEIAAAISAVPYGVDPREAARAFYEAGGGAPPGAELDEEEYTKAAELVLE
YPELKVEYRIEFDDGIARLSPEEWKEFATKPARVLVRFFEERGDEKLAAAYREGLAGAEKAAEERFGSGSHH
WGSTHHHHHH

Ub-peptide ligase 6 H6

MSGGGRGYVIVDPEVSDEEARAWKARADEVARRLGELLGVPVALAVAVVLKDGKPVIAHVFAVDPEDPSK
VIDPLPFQLLAIERGILKKTRGGCGEIAAAISAAPYGVAPEEAARAFYEAGGGAPPGAELDEEEYRRAARLV
NEYPELKIEYKIEFKDGIARLSPEEWREFATRPAEVLVEFYARRGDEKRAAAFREGLAGARAAAERRYGSGS
HHWGSTHHHHHH

Ub-peptide ligase 7 H6
MSGGGRGYAIVDPEVSDEEARAWKARADEVARRLGELLGVPVALAIAVVLKDGKPIVAHVFAVDPEDPTRV
IDPLPFQLLAIHEGILKKRVGGCGEIAAAISATPYGVDPREAARAFYEAGGGAPPGAELDEEEYLEAAELVNR
YPELKVEYRIEFRDGVARLSPEEWKEFATKPARAIVEFYEKRGEKKIAEAFRKGLEGAEKAAEERYGSGSHH
WGSTHHHHHH

Ub-peptide ligase 8 H6
MSGSGIGGAIADPEVPDADAKAIAKKAEEVAKKLGELLGVPVGFAAVVRLEDGRPVVWHVFALYPEEPDRL
VDPLPYQYLAFLHGVLKNRGGGCGVIAAVLAAAPYGVDPRKVARAYFEAGGGLPPGAKLTPEEAERLAEF
AVERPELRVEFRLAPEDGVWRPDGKQWVELIRKPAEVIIREAYRAGKPEAAKKAEEALEKALKKAEERAGS
GSHHWGSTHHHHHH

Ub-peptide ligase 9 H6
MSGSGIGAAIVDPDVPEEAARAIAREAERVARRLGELLGVPVGFAAVVRLEDGRPVVWHVFALYPEDPNRIV
DPLPYQVLALAKGILKHIGGGCGIIAAVLAARPYGVDPEKVARAYFEAGGGLPPGAKLTEEEKIALARFAAE
HPELEVEFEVEPEDGVIRLTPEQWVELIARPAELIIEELRRLGKPEAAEKAEEALRIAVEKARARAGSGSHHW
GSTHHHHHH

Ub-peptide ligase 10 H6
MSGSGRGIAIADPSVPKEEAQRWARRAEAVARRLGELLGVPVGLAAQVYLKDGEPVVFHVFALDPEDPNRI
TDPLPFQVLALYKGILKKQGGGCGEIAAVLAALPYGVDPERAAEAFYEAGGGYPPGFELTEEERRALGRFV
EEHPELIVEFDVKVEDGVAYPDGKTWVEFAKKVAEKIIEFLRKEGKEEEAEKFEKAAEKGIEAAKAKAGSGS
HHWGSTHHHHHH

Ub-peptide ligase 11 H6
MSGSGIGGAIVSPDISDEEARAVVERAQAVADRLSELLGVPVGFAAVLAREDGRPVVWHVFAIDPEDPNRLY
DPLPFQIYALAKGLIKSRRGGCGAVAAVLAAVPYGVEPERAARAFFEAGGGLPEGGKLTEEELTRLAELVVE
RPELIVEFDIPLRDGIARLDAEQWAELARRVADVVIEALRKEGLDEKAEKIEEAKKKGVEKIRAAAGSGSHH
WGSTHHHHHH

Ub-peptide ligase 12 H6
MSGLEIEFEGEEEKEAIVEEHYKKATPEEKEAIEKGKEAYEKLKEAFKKAVKEAVEGGKRRAEVKVGDATV
VVELVPLSSGAKVGIVYAQDKDGNGLHTYAVIVEGDKAKIIDPFPRQYREGLNGGCTPLAAEYILEEAARLS
SPEEAEKAVKELLKDGGAIADTKEGAEKAKELFEKLGFKAEVRPHPTLPGKWLAEIVPEGSGSHHWGSTHH
HHHH

Ub-peptide ligase 13 H6
MSGIKIGWEGEKEYEEIYEEVYKKATPKEKEALEKGREARKVLKEGLKEAVKKAVEEGKEEAKAKAGDVE
VKAKFYKLSSGNKVGIVYYVDEDGDGGHVYAVIIKGDEGKIIDPFPQQYKEGGDGQCVVYAARIIEEKAKE
LANEEEAKEAVRYLFKDGGALARTRAGAEAAVEALKEYGFEGEVRPHPRYPGRWEADIWLPGSGSHHWGS
THHHHHH

Ub-peptide ligase 14 H6

MSGVAPALAEKYRELADLLAETDQAQIDAVLKRLEEIEGKPVEGGALVYRRGDEREWVPISAEGARRVAEG
LRRAAEGEVDVVGGCLLGAIAIAGRAIEKGEAGSVAPVGGGHHYVAVVVHPEDGKATVYDPQAPIIAEGIR
QRFDRLAPQESPEARARVREALERAADAIARLATDEIRGTPEEVAEKVAELAREAAEAAREAERAAETPELR
ELARAVREYAEGFIHDAPNIRRVAEKLAEYAGSGSHHWGSTHHHHHH

Ub-peptide ligase 15 H6
MSGVDPALAEAYRRLAELLRETDQAGIDAVLANLEKITGKPVEGGALVYRKGDEEKWVPLDEAGAREVAR
GLELAAEGKVDVVGGCLLGSIAIGARALRAGKAAAVAPVGGGVHYVAVVVDPATGEATVYDPQAPIIAEGI
RQRFERLAPQESPEAREKVREALERAAEAIGRLASDEIRGTPEEVARKVAELAREAIEAAEEAYKAAENPEL
KELARAVAEYARGYIADAPNIEKVARKLRELAGSGSHHWGSTHHHHHH

Ub-peptide ligase 16 H6
MSGLSPALAAAYRALAERLERLDDEGIERVLAAYEELTGRPIEGGDLVYRRGDEEEWVPISAEGARRVAEGL
RAAAEGRYDVAGGCFLGAIAYGAEALRAGEAATIAPVGGGAHWVAVIVDPADKKARVIDPQADIIAEAIVE
AFERYAPNEPPEKREKVREALEKAAEAIRKLNSDEIYGTPDEVADKVVELAKKAIEEGKEAYEAATDPRVKA
IAKAVIEYAKGYIEDAPEIRRAAEFLQRRAGSGSHHWGSTHHHHHH

Ub-peptide ligase 17 H6
MSGVSPELAQAYRDLADVLRSVDDEEIEAVLANYEKITGEKVEGGALVERRGDEEEWVPISAEGARRVAEG
LRRAARGEVDVVGGCLLGAIAIGGQRLREGKAATIAPVGGGNHYVYVGVDPADGNAEVYDPQAPIIAAAIR
ERFERLKPEEPEEEAEEVKKALEEAADAIAKLASDEITGTPEEVAEKVRELAKKAIENGKKAYEAAKNPKVK
ELAKAVIEYAKGFIADAPEIERAARKLAELAGSGSHHWGSTHHHHHH

Ub-peptide ligase 18 H6
MSGLEETKEAVKRLEDEAIEKALKEGRIKLYFYDWKDGKKYPAEKGGGTGGGCLLLAVYAALGALEAGAE
EAEVVGIAGRRNPDGSAEAGHFVAVAKGPKVPGGEVWIDPAALQYAAWLLVGRGILTPEEAFEKAAELARK
LGYLAVKLGLNNEEIREVDLELARELLEYAGHPELAPEVELNGRGLHALTEGAIGIEEGDAERAKKAAEELR
KLAEEFRKAGRPELAETVEKQAEYFEEKAEKLGSGSHHWGSTHHHHHH

Ub-peptide ligase 19 H6
MSGDAEARATIARLSDERLARALAEGRIRLYAYDWKTGERYEVEEGGDTGGGCLGLAVEAALGALEAGAR
RAEVLGVAGRRNPDGTFTAGHAIARAEGPKAPGGEVWIDPAALQLAAWALIGAGVLTPEEAYERAAELARE
LGRRAVAEGLDQEEIRELGIELGERLLKEAGREELAPRVRLRAEALYALTEGVLALYRGDAEGALRAAERLE
RLAEEFEAAGEPEGAERLRELAAQLRERAAELGSGSHHWGSTHHHHHH

Ub-peptide ligase 20 H6
MSGLEEVRKRVKALEDEALEEALKEGRIELEYYDWRTGERYPAKEGGGTGGGCLLLAVYAALGALEAGAT
RARVVGIAGRRHPDGTATAGHFVAEAEGPKVPGGRVWIDPAALQYAAWLLVGRGILTPEEAFEKAAELAFIL
GYLAVKLGLDEDEIEELDLALAGALLEYAGHPELWPEVELVGRALHALTRGAIALEEGDAEAARAAAEELR
RLAEEFEKAGRPGAAELVRRQAEYFEEEAAKLGSGSHHWGSTHHHHHH

Ub-peptide ligase 21 H6
MSGWTPEEVRAAAARVGEALAAGDRREAGEWAAWVGIAAIEAQLREEYGVRILGTRVEWIRFPDGKEYPV
GVVEVPADEVRPAVLERGAPHAREGGLLAGALPLYDEDGRVVHVAAYYEEPDGRATILDPLGIQVERGLAG
GCIALAGILLALRGFREAAVRVVAEGGGRLTPEEREYLAGLAEALGLPELAEAIRRAYAGSGSHHWGSTHHH
HHH

Ub-peptide ligase 22 H6
MSGWTPEEVRARARRVGEALRAGDRERAGREAAWVGIAAIVAQLREEYGVRILGTRVERVDFPDGRSYPV
GVVEVPADEVRPAVLERGAPHAREGGLLAGALPLYDKEGRVVHVAAYYERPDGRAEILDPLGIQVERGLAG
GCIALAGLLLALRGFREAAVRVVREGGGDLTPEEREWLAGLAEELGLPELAEAIREAFAGSGSHHWGSTHH
HHHH

Ub-peptide ligase 23 H6
MSGWTPEEVRALARRVGELVAAGRRDEAARLAAWVGIAGIEAQLEQRYGIRVERTEVREVAFPDGRTYPIG
VVHVDAAQVRPVVLDRGLPHAEEGGLLAGAIPVFDEDGRAFHVNAYYERPDGTAEILDPLGYQVKVGFGG
GCVGLAGALLAHRGFPEAGVRVAREGGGEVDEETLEYLVGLAEALGVPELAEAIREAYAGSGSHHWGSTH
HHHHH

Ub-peptide ligase 24 H6
MSGLTPEEARALAREIAALVKAGDRDRAARLAARLGVEAYRAYLRERYGVRILGEEVRKIRLPDGEEAWV
GVLRVPAAEVRPVIEELGAPWAREGGLLAGALPLVDPEGRPWHVAAYVERPDGTAEILDPLGIQVQRGWGG
GCLFLSAILLALQGYPEAAVRVAREGGGPVPPEAREYLVRLAEAAGYPELAEAIRAAFAGSGSHHWGSTHH
HHHH

Ub-peptide ligase 25 H6
MSGATPEEVRALAERIAELIRRGEREEAARLAAELGILGFAAWLKERYGVEILGRRVVEEPLPDGRRVPIGVL
EVPADQVRPVLEELGAPWAREGGLLAGALPLVDDEGKVVHVAAYVERPDGTAEIYDALGAQVLYGWKGG
CLGLAALLLAKKGYPEAGLRVAREGGGKVDEKALKRLVELAEALGYPELAEAIREAYAGSGSHHWGSTHH
HHHH

Ub-peptide ligase 26 H6
MSGITPEEVRERARKIGEKYKAGKRDEAAEEAARLAAEAVAKWLEEEYGVKPEEVRAAAEELPDGSKRYY
ARLRIPAEDLRRILLENGYPHAEEGGLLAGIVPIRDENGEIYHVLAAVEHPDGTAELYDFLPQQIYEGLDGGC
LGLAAVELARRGYPEAGLRAAREGGGRYGPKALAYVEEEARAIGHPELGELLRRRYAGSGSHHWGSTHHH
HHH

Ub-peptide ligase 27 H6
MSGVTAEEVRERAREIGEKFRAGEREEAALEAARLFAEALAAYLEREYGIPARGAGARRERLPTGEEAVVG
RVEISAEDARRILEKAGAPHAEENGQVAGLIPVRDADGKAWHLVAVYTNPDGTAEIRDFLGTQLLHGYRGG
CYGLAVLELARAGFPGAAERAAEEGGGRAGPRVRELLRRLAEALGRPELGERLAEGARGSGSHHWGSTHH
HHHH

Ub-peptide ligase 28 H6
MSGATPEEVAELARRIAELLEAGRRDEAARLAAELGLRALDAYLAERFGVRLTRREVKRDKFPDGREIPYG
VGRLSTEDARRVLLEPGYPHAEEGGLLAGAFPLWDPDGRPVHAIALVERPDGTADLYDFLAAQVKYGLNG
GCLAGAAYALVKEGFPEAARAVVEQGGGVADERARAFWRDAIARLGHPDIGDAAAERLAGSGSHHWGST
HHHHHH

Ub-peptide ligase 29 H6
MSGITPEEANARAREIAELYAAGRRDEAARRAAELALELLDQYLKEKYGVELTDREVAWEPLPDGRRLPYG
RARLSREDAEKVLVEPGYPHAYEGGRLAGAFGIFTPDGKPVHVVALYEEPDGTVEVLDFLPAQVYYGFIGG
CLAAAAYVLVKRGFPEAARRVVEEGGGVADEKARDFFRDLIRALGHPDIGDLAAERLAGSGSHHWGSTHH
HHHH

Ub-peptide ligase 30 H6

MSGLTPEEANARAREIAELYAAGRREEAARRAAELAGELLDQYLKEKYGVELTDRRYEELTAPNGTPLPYF
RATLSREDARKVLVEPGYPHAYVDGRLAGAFGIYTPDGRPVHVVALYEEPDGTVVVYDFLPAQVYHGFRG
GCLAAAAYVLVQRGFPEAARAVVEKGGGVADEEAAEYYADLIRELGAPEIAEPARERLLGSGSHHWGSTHH
HHHH

Ub-peptide ligase 31 H6
MSGVTPEEANREAREIAELYAAGRREEAARRAAELALRLLDQYLKQTYGVSLTDRRVEWEPLPDGTRLPYG
RATLSREDARRVLVEPGYPHAYEGGRLAGAFGIYTPDGRPVHVVALYERPDGRVEVVDFLPAQVYYGFIGG
CLAAAAYILVKRGFPEAARRVVEEGGGVADEKAREYYRKLIEDLGYPDIGERAAEILAGSGSHHWGSTHHH
HHH

Ub-peptide ligase 32 H6
MSGITAEEAEAEAREIAKLIAEGKKDEAARKASELAAKVLAKYLEEKYGIKPKGYRAEEEEYPDGTKLYWG
VIEISAEDARKILEEPGYPHAREGGLLAGAFPILDDKGRPVHTVAVVERPDGTAEVLDPLGTQVYYGYPGAC
LALSAYFLAKEGYPAAARAVVEQGGGKASEEARAFVRELIERLGYPDIAERAAERLAGSGSHHWGSTHHH
HHH

Ub-peptide ligase 33 H6
MSGITAEEANARAREIAELIAEGKKDEAARKASELAAEVLAEYLKEKYGIEPSGYRAEEEEFPDGEKLYWG
VIEITAEDARKILVEPGYPHAEEGGLLAGAFPVLDDEGRPVHTVAVVERPDGTAEVYDALGTQVYYGYPGA
CLALAAYFLAKEGFPAAARAVVEQGGGKASPEAREFVRRLIERLGYPDIAEKAAEVLKGSGSHHWGSTHH
HHHH

Ub-peptide ligase 34 H6
MSGFTPEELRALAREIGELIAAGKREEAEEKAAELAAKGIAAYVESEYGIPPLEVRVEKVRFPDGREYWVGV
VRFPAEKVLEVIKERGFPYAREGGLYAGALPIIDEKGRGVHVIAFVEHPDGTATLYDFLPVQIREGIRGGCAG
LAAIYLAHLGYPEAGRRILEEGGGFADERARRLLEELAEAVGYPELGRLGYERFRGSGSHHWGSTHHHHHH

Ub-peptide ligase 35 H6
MSGITPEELRELAREIGELIAAGKREEAEKKAAELAAKGIAAYVEERYGVPPLEVRTEKVKFPDGKEYWVG
VARFPAEPLREIIRERGFPLAREGGLWAGALPYVDDDGRGWHVVAFVEHPDGTAELYDFLPIQLREGVNGGC
AGLAGIYLAYRGFPRAGRRVLEEGGGVADEKAREVLREIAEAVGFPELGELGYERFRGSGSHHWGSTHHHH
HH

Ub-peptide ligase 36 H6
MSGATAEEIRALAAEIGELIAAGDRDRAEELAALLGGLGIEGYLLERYGIKALRIRPVRIELPDGRPAVIGEAV
FPAEEVRRVLLEEGFPWAEEGGRLAGAIPLWHPDGSPVHVVAYVEEPDGTAVLYDFLPIQIRHGLRGGCVGL
AGVYLAARGFPRAGLLTVEQGGGPADEKAREFVAGLAEAAGFPELAEAIRRRLAGSGSHHWGSTHHHHHH

Ub-peptide ligase 37 H6
MSGYEEEAERLVAAVPEIVERAKELKEELEKEGKKARVFSIGGHAYAVIKINGEFYLYDGLLAQARAAAAAA
GRPLPEEVRAAIDRYYEALEKEAEKAGLKLLYPENEDAIAAAAVEAVAEVAGLTPEEAAREACIAIAGGVIA
QEVGLEEALEAVKGGGTRSGAPAEPIVRRVAEALAGSGSHHWGSTHHHHHH

Ub-peptide ligase 38 H6
MSGYDAEAERLIAAVPEIVERAKELKEELEKEGKKAKVFSLGGHAYAVIKIDGEYYLFDGLLEQAEAAAAA
AGRPLPAEVRAAIERYLAALEEKAKEAGLKVLRPENADDIARAAVEAVAEVAGLTPEEAAQEACIAIAGGVI
AREVGLEEALKAVEGGGTRRGVPAEPIVRRVYERLQGSGSHHWGSTHHHHHH

Ub-peptide ligase 39 H6
MSGAAAAADRLIAAVPEIVEAAKRAARELEETGRRARVFVVGGHAFAVIRRDGRFYLYDGLLAQARAAAA
AAGRELPAEVRAAIERYLAALEEEAAKAGLELLRPANEDEIARAAVEAVAEVSGLSPEEAAQEACIAIAGGVI
AEEVGLEEALKAVAGGGTRSGAPAEPIVRRVAELLAGSGSHHWGSTHHHHHH

Ub-peptide ligase 40 H6
MSGAAAAAEKLKAAVPEIVARAKELAEELEKEGRKAVVFSIGGHAYAVIRKDGKFYLYDGLLAQAEAAAA
AAGRPLPADVRAAIEEYLAALEEEAAKAGLETLRPEDADEIARAAVEAVAEVAGLSPEEAAGEACIAIAGGV
IAREVGLEEAVRAVKGGGTRSGAPAEPIVRRVAELLAGSGSHHWGSTHHHHHH

Ub-peptide ligase 41 H6
MSGAAAAAERLIAAVPEIVEEAKKVAEELEKKGEKAKVFSIGGHAYAVIKIDGKYYLYDGLLAQAEAAAAA
AGRPLPADVRAAIERYLEALKKKAKEAGLETLDPEDAVEIARAAVEAVAEVAGLSPEEAAGEACIAIAGGVI
AKKVGLEEAVKAVEGGGTRAGAPAEPIVRRVAELLAGSGSHHWGSTHHHHHH

Ub-peptide ligase 42 H6
MSGARFAVLEYKKPLTPEEQAKVDEAKELAAQGKAYEAAELLTEVHRRRAREVVAEHPEVDVGLVRIGGH
FVAFRREDGEIRVADPLRIQIEAARAYAEALREGKSKEEAFKKAAEVAAEGVRELLAEVFGASEEAREIAAEV
VRIALPYTFVGGGCLVLAAAAVAGLLADLGDREAGREALERVAREGGFAPTPAQIEAWGLTREEAEAYAAR
APEAIREAAEKALAAGSGSHHWGSTHHHHHH

Ub-peptide ligase 43 H6
MSGVRAAILEWDGPLTPEEQAAVERAREAAAAGDAETALALLQELFRRKAREVIEANPDVDVGLVNIAGHF
VAFERRDGEIRVYDPLPIQYEARKYIDEQVKKGKSIEEAIKEAAEYAAEKVRELLLEEAGVSEEAAEVAYEIV
KKAFPYDIYVGGCTTLATAIAVGLLYDRGALEEARRFARRVADEGGFEPNAKQAARFGISEEEAKEIAKKTP
EAIKEAADEALAKGSGSHHWGSTHHHHHH

Ub-peptide ligase 44 H6
MSGPDEVRIYRADGVERLPFADLSPEAKRRAAEALARALELGARAAADPTPEALAEAGLALVDAFVAVLEG
YGFDVSYPREELRALAREAAERKLADPSPEAIRAAYEEALERIAELLREALKEKGLSPEDAVIIVIGGHAVPA
AIRGKKLALVDPLPVQVALGVRGCTVIALEALRRVGAPRAVLDRVAKEGGVLIPADDPLRELLAERAREFGL
EVIEVPGSGSHHWGSTHHHHHH

Ub-peptide ligase 45 H6
MSGRDAVEIIRPDGEERIPLAEAPAEKRARYEELKAEVDKALAAAAADPTPEAIADAAVALLKLRARQLELA
GFEPIIDYDEYRERIKEYRGAILEDPSKEVREKVYEEGIKYLQEKLKKLLEEEGIPPEDIVIIYVGGHWFAAAI
KGKKLIVVDPLIIQAAIGANGCFALALKIAEGAGAPRELIERAAKRGGQLVRKDDPVKDLVVKLAEEYGLPI
YEIPGSGSHHWGSTHHHHHH

Ub-peptide ligase 46 H6
MSGRDAVEIIRPDGREVIPLAEAPAEKREEYEKLKAEVEKALAKAREDPTPENIADAAVALLRLRAKQLELA
GFEPIIDYDEYRERIKEYRKAILEDPSKEVREKVYREGIEYLGEKLKRWLKEEGIPASDIVIIYTGGHWFAAAI
KGKKLIVVDPLIIQAAIGANGCFALSLEIARRAGAPEELIEFAAKQGGQLVREDDPVRDLVEELAKKYGIPIY
VVPGSGSHHWGSTHHHHHH

Ub-peptide ligase 47 H6
MSGDEERFGKEFEEEVYKLTGAKIPPGGGTRGGGCLALAAAAVHLFGEGEIVPVAGGWHVVAAIPVGGKWI
VIDPLPFQIYARHLVSKGKSIEEAAREAVERLREIVEALKRGDLEKARKIVRELLEEVLEEVLKNPKLTPEEKE
AIRKTYEAVFEAIEELFEPTDDPAKAAEAAAAKIGEAYKAATGSGSHHWGSTHHHHHH

Ub-peptide ligase 48 H6
MSGAEITRVERPEHRFELTPEERAALAEAREAAARALAEGRPEEAYGLLLRKFGEILLRHAPTPAARAVAEA
LIEIGERALRGEVWPSAVVPLTPEEVTEALAELYGEAELLRLDFGGRQAAIVGFAGHAVLLEPVPGGTRITDP
GAAQFRAEQVLARRGKRVEPGRAPELLEAAVEAGILPISVEQAREAYDRFLAAGGGDSGGCAALAAARAV
ERGSGSHHWGSTHHHHHH

#### Ub-PCNA Ligases 1-48

***Note:*** *Residue numbering refers to the designed sequence and excludes the N-terminal residues contributed by the expression construct (Met for the Ub-PCNA ligases, Met-Ser-Gly for the Ub-peptide ligases).*

Ub-PCNA ligase 1 H6
MVLEEVREDLRELGVDEEAIEELLAAARKAQAALDANDVKGFRAALEELRAAIKKAAATAKEVTPKGEFR
KLPDGSIAIELELPDGTTIEVVALIYGQGTCAALAVAILKALGKPGRIIPYPIPTPFGLTTHVAVAPGGDPEKTID
PGNWGGSDRDLERIVDAIEKKLKEKGIELTPEQREKLREEIKEAYELLREAQTEAGKGGSHHHHHH

Ub-PCNA ligase 2 H6
MVEERIRRELRELGVDPEAIERLLEAARAAQAALDANDVAAFRAALAALREAIREAARTAKAVRPRGDFYV
TPEGDVAIELELEDGRVLELIDKPGADGGCAGLAAAVLRALGEPGYIIPYPIPTPFGLKPHVAVAPGGRPDAAI
DPGNYGFGNTTVDKIVERIEKKLAAKGIELTPEQRERLREAVAQFEELYLKALKEKEAGGSHHHHHH

Ub-PCNA ligase 3 H6
MVEEIVREVLRELGVDEAAIETLLDAARKAQAALDANDVKGFKAALAELREAIRQAARTAKEVTPVGEFR
KLPDGSIAIELELPDGRKIEVVAFPVGGGSCAALAVAILKALGKPGKIIPYPIPTPFGLTTHVAVAPGGDPEKVI
DPGNYGGDDTPLDRIVDAIEKKLAEKGITLTPEQRARLREEIARARELLREAQKEAEKGGSHHHHHH

Ub-PCNA ligase 4 H6
MIEEEIREWLRELGVDEAAIEKLLEAARKAQAALDANDVAGFRAALEELKAAIKEAAATAREVTPEGEFRP
LPDGSVEIELRLPDGRKLTVVVDPKLGGYCAGFAVAILEALGRPGRIIPYPIPTPFGLRPHVAVAPGGDPEKVID
PGNYGGDVVDFDKIVDNIERKLREKGIELTPEQRARLRERIEALVAALKAAQAAARAGGSHHHHHH

Ub-PCNA ligase 5 H6
MVLREIREQLRELGVDPAAIDRLLAAARAAQAALDANDVKAFRAALADLRAAIREAAATAKEVRPEGDFYI
TEDGDIAIDLRLADGRRLRLIDPPGPKGGCAGRAAAVLRALGEPGYFIPYPIPTPFGLKPHVAVAPGGRPDAAI
DPGNHGIGQTTVDKLVERIEAKLRAEGIELTPEQRERLRAALEEFEKLYLEALAKKQKGGSHHHHHH

Ub-PCNA ligase 6 H6
MVLEEIREQLRELGVDPAAIERLLAAAREAQAALDANDVAAFRKALAALKEAIREAAATAQEVRPVGDFYV
TPDGDIAIELELADGRRLRLIDVPGADGGCAGYAAAVLRALGEPGYVIPYPIPTPFGLKPHVAVAPGGRPEAA
IDPGNYGPGNSTVDKRVARIEKKLAEEGIELTPEQRARLRAALEEFERLYLEALAEKKKGGSHHHHHH

Ub-PCNA ligase 7 H6
MVLEEIRAQLRELGVDPDAIERLLAAAEAAQKALDNNDVKAFRQALADLREAIREAAATAQEVTPEGDFY
VTPDGDVAIRLRLADGREFELIDVPGPDGGCAGRAAAVLRALGEPGYVIPYPIPTPFGLRPHVAVAPGGRPDA
MIDPGNWGPGRTTVDKLVDRIEAKLRAEGIELTPEQRQRLRDALAQFERLYKEALAKKREGGSHHHHHH

Ub-PCNA ligase 8 H6

MVLEEIRAQLEELGVDPAAIDRLLAAAEAAQAALDANDVEAFRKALAELKAAIREAAATAQEVRVEGDFY
VTPDGDIAIRVRLADGRTLEVIDVPGADGGCAGRAAAVLRALGEPGYIIPYPIPTPFGLKPHVAVAPGGRPEA
AIDPGNGGFGNTTVDKIVDRLEALLAAEGIELTPEQRERLRAAVAEFYRLYKEALEKKKKGGSHHHHHH

Ub-PCNA ligase 9 H6
MVEEEIRAQLRELGVDPAAIDRLLEAAREAQAALDANDVAWFRAALARLREAIREAAATARPVTPEGDFYV
TPEGEVAIRLRLDDGRVLELIDVPGKTGGCAGYAAAVLRALGEPGYVIPYPIPTPFGLVPHVAVAPGGRPEAAI
DPGNWGAGRTDVDKIVARIEAKLAAEGIELTPEQRERLRAAVEEFERLSLAALAEHESGGSHHHHHH

Ub-PCNA ligase 10 H6
MVEEEIREQLEELGVDPAAIDRLLEAARAAQAALDANDVAAFRAALAALKEAIREAAATALPVTPVGRFYV
TDDGDIAIKARLADGRELEVIDVPGADGGCAGRAAAVLRALGEPGYIIPYPIPTPFGLTPHVAVAPGGRPDAA
IDPGNWGAGRTTIDKLVERIEAKLKAKGIELTEEQRERLREALAEFEELYKEALAKKAKGGSHHHHHH

Ub-PCNA ligase 11 H6
MVEEAIRRDLRELGVDPEAIERLLEAARAAQAALDANDVAAFRAALAALREAIREAARTALAVRPQGDFYV
TPEGDIAIELELEDGRVLELIDMPGESGGCAGLAAAVLRALGEPGYIIPYAIPTPFGLKPHVAVAPGGRPDAAI
DPGNYGPGNTTVDKIVETIEKKLAAKGIELTPEQRERLREAVAEFEELYLKALKEKAAGGSHHHHHH

Ub-PCNA ligase 12 H6
MVLDEIREQLRELGVDPAAIDRLLAAAEAAQAALDANDVKAFRAALADLKAAIREAAATAWEVRPEGDFY
ITEDGAIAIDLRLADGRRLRLIDWPGADGGCAGRAAAVLRALGEPGYIIPYPIPTPFGLKPHVAVAPGGRPDA
AIDPGNWGTENTTVDKLVERIEAKLRAEGIELTPEQRERLRAALEEFEKLYKEALAKKRAGGSHHHHHH

Ub-PCNA ligase 13 H6
MIEDEIREQLRELGVDPEAIERLLEAARAAQAALDANDVAAFRKALAELREAIREAAATAMEVRPVGEFYV
TPEGDVAIDLELADGRRLRLIDVDGPYGGCAGRAAAVLRALGEPGYVIPYPIPTPFGLVPHVAVAPGGRPEAA
IDPGNWQPGNTTVDKLVERIEKKLAEKGIELSEEQRARLREALAEYERLYKKALAKKRSGGSHHHHHH

Ub-PCNA ligase 14 H6
MVLEAIRAWLEELGVDPAAIDALLAAAREAQKALDNNDVAGFRAALARLKEAIRAAAATAREVTPEGEWR
KLPDGSVEIELRLPDGRRLRAVVDPDGGGNCTGLAIAILRALGRPGRIIPYPIPTPFGLIPHVAVAPGGDPEKVI
DPGNYGLDDVDFDKVVDRIEKKLREKGIELTPEQRERLREEIKRIYDLLKEAKAEELAGGSHHHHHH

Ub-PCNA ligase 15 H6
MGDATFEYVKAAWEAGYALAEELARELPPGEAVLVPVPVIHPGDLDNVRAHVLLAVRHPDGSTEVIDVVN
AGGNAEWGLKGKEILKRFTAREARELARELGVDARPAAPDDDELYLAKNKFGKYKVASKSGKGACVGGA
VWVAAERGFRLRIADMDRVEARYREALLAAGASEEEADRIIALFREALGQEGGSHHHHHH

Ub-PCNA ligase 16 H6
MEDVTWEIAKAAWEAGYELARRLAARLAPGEGILVPVPDLLPGDADKVTPHVLLGVRRPDGSVEVIDVRN
GGGDVKWLLEGRKVEEVLTNREAAELARTLGVDAENTGKPEDKIYFARRPDGRPFVAAEGVGGGCVGGA
VWVARKRGLRLRLAPLDEVLEAYRAALLAAGASPELADRLIELFRIGLGHAGGSHHHHHH

Ub-PCNA ligase 17 H6
MEEVTWEQAKAAWDRGYELARELAAELAPGEGILVPVPVLLPGETEKVTPHVLLGIRRPDGSVEVVDVTN
GGGDVKWLLEGKEVLEEITRAQARELARELGVDARPLGEPDDKIYIAKKPDGSWFVAGEGKGGGCVAGAV
YVAKERGLRLRLAPRDRVLEAYREALLAAGASPELADRIVALFEEGLGYGGGSHHHHHH

Ub-PCNA ligase 18 H6
MEEVTWEQAKAAWERGYELARELAERLEPGEGILVPVPDLLPGDADRVNPHVLLGVRHPDGSVEVIDVRN
GGGDVKWLLEGREVEEVLTRAEARELARELGVDAAPTGAPEDKIYFARRPDGRPFVAFEGKGGGCVGGAV
WVARKLGLRLRLAPLDEVEAAYREALLKAGASPELADRLVALFKEGLGYAGGSHHHHHH

Ub-PCNA ligase 19 H6
MEKVTWEIAKALWDAGYELARELAAELEPGEGILVPVPDLLPGEDDKVTPHVLLGVRHPDGSVEVIDVRN
GGGDVKWLLEGRKVREVLTRAEARELARELGVDARTYGKPEDKIYFAERPDGRPFVAAEGKGGGCVAGAV
WVAKELGLRLRLAPRERVEAAYREAALAAGASPELADRLIELFREGLGWGGGSHHHHHH

Ub-PCNA ligase 20 H6
MAEVTFEHVKAAWKAGYELAEELARELPPGEAVLVPVPELLPGDIRNVRAHVLLAVRHPDGSTEVIDVVNA
GGLAKFGLKGKEILAKFTAAEARALARELGVDARPRAPDDDTLYLAKNKFGRYKVASESGKGACVGGAVK
VAADRGFRLKIADYDRVAARYREALLAAGASEEEADRIIALFREALGVGGGSHHHHHH

Ub-PCNA ligase 21 H6
MEEVTWEQAKAAWDAGYALAEELAKELKPEEGILVPVPDLLPGQKDRVVPHVLLGVRHPDGSVEVIDVRN
GGGDVKWLLEGREVEEVLTRAEAAALARELGVDAETLGKPDDKVYFARRPDGDPFVAAEGMGGGCVAG
AVWVAKKLGLKLKLAPRERVEERYRSALLAAGASPELADRLIALFREGLGWMGGSHHHHHH

Ub-PCNA ligase 22 H6
MPAVVFPEFDPYDPAFPEEVNERQRKELNEIAKKFVEEFGVEVKRLDDEEAYKAWEKSKKAFIEAIKEGKSI
REAYEASKRAAAEVAVEIAKKKGLADKRVVIVLPVAEPSFDSDDLTNPKFVPHYVAVVIEGGKVVAILDPGN
PGYAQGTCGVGAVLIGALVAKGISPEEAQKIVREAVFEAARELGAKASALEIIKKALEKIKQRIGGSHHHHHH

Ub-PCNA ligase 23 H6
MTKLILPPVEINDPNLSAEQTERNDREVKEAVREFAARLGVTVLELDHEEIRAAWEAAREAALEALRAGKP
PEEAFRAAKRAAAEKAVEIVKRRGLGDEDVVIVVPVAVPSYEEDENGEPKVISHMVAVVIRKGKVEAIIDPG
NPMGVPGTCYAGAGLAGALVAAGVSPEEAERLVREALREAYKELGNKATAGEIVRRAVERILERLGGSHHH
HHH

Ub-PCNA ligase 24 H6
MTKLILPEVRIFDPNQTEEQNEFNDREVKEAVREFARRLGVEVLRLDDAEIEAAYEAATEAALEALRAGKPP
EEAFRASQRAAAEKAVEIVKRRGLGDEDVVIVVPVAVPSYEEDENGEPKVISHMVAVVIRNGKVEAIIDPGN
PGGIPGTCYAGAGLAGALVAAGVSPEEAERLVRAALREAYKELGNHATAGEIVRRAVERILERLGGSHHHH
HH

Ub-PCNA ligase 25 H6
MTKLILPEVTLEDPNQSGEQNANNDERVKKAVEDFARELGVEVEYLDDEEIEKAWEAARKAALDALKAGK
PPEEAYRASVEAAAEKAVEIVEERGLGDEDVVIVVPVAVPSYEEDENGEPKVIGHMVAVVIRDGRVEAVIDP
GNPGGVWGTCYAGAGLAGALVAAGVSPEEAERLVREALREAYKELGPKANAGEIVERAVERIRERLGGSH
HHHHH

Ub-PCNA ligase 26 H6
MMKLILPEVEIVDANLSEEQNTRNDREVKEAVREFARRLGVEVLELDEEEINAADEAAREAALEALRAGKP
PEEAFRASQRAAAEKAVEIVKRRGLGDEDVVIVVPVAIPSYEDDENGEPKVISHMVAVVIRNGKVEAIIDPGN

PGGIFGTCYAGAGLAGALVAAGVSPEEAERLVREALREAYKELGNKATAGEIVRRAVERILARLGGSHHHH
HH

Ub-PCNA ligase 27 H6
MTKLILPEVQIVDKNLSEEQNERNDREVKEAVREFARRLGVEVLELDWGEIERAWEAAREAALEALRAGKP
PEEAFRASQRAAAEKAVEIVKMRGLGDEDVVIVVPVAVPSYEEDENGEPKVISHEVAVVIRHGKVEAIIDPG
NPGGVKGTCYAGAGLAGALVAAGVSPEEAERLVREALREAYKEIGNKATAGEIVRRAVERILERLGGSHHH
HHH

Ub-PCNA ligase 28 H6
MTKLILPEVEIYDENKTEEQNKVDDERVKKAVEEFARELGVEVEYLDDEEIEKAWEAARKAALDALKAGK
PPEEAYRASVEAAAEKAVEIVRERGLGDEDVVIVVPVAVPSYEEDEAGKPAVIFHAVAVVIRDGRVEAVIDPG
NPGGIQGTCYAGAGLAGALVAAGVSPEEAERLVREALREAYKELGNKATAGEIVERAVERIRERLGGSHHH
HHH

Ub-PCNA ligase 29 H6
MTKLILPKVEIVDPNLTEEQNKENDERVKKAVEEFARELGVEVEYLDEQEIAKAWEAARKAALDALKAGKP
PEEAYRASVAAAAEKAVEIVRERGLGDEDVVIVVPVAVPSYEEDENGEPKVIGHMVAVVIRDGRVEAVIDPG
NPGGILGTCYAGAGLAGALVAAGVSPEEAERLVREALREAYKELGTKANAGEIVERAVERIRERLGGSHHH
HHH

Ub-PCNA ligase 30 H6
MTNLILPEVTIYDDNLSEEQNKENDERVKKAVEEFARELGVEVLYLDDGEINKAWEAARKAALDALKAGLP
PEEAYRASVEAAAEKAVEIVRERGLGDEDVVIVVPVAVPSYEEDENGEPKVIGHAVAVVIRDGRVEAVIDPGN
PGGVQGTCYAGAGLAGALVAAGVSPEEAERLVREALREAYKELGIKATAGEIVERAVERIRERLGGSHHHH
HH

Ub-PCNA ligase 31 H6
MKVRVEKVEVEGDDIDAQIKAYEKAREKAKELAKKAEKGEIIIVAAVLNTSNGKEAYHLFIVDGETGKVIDP
GNKQTKDGVAGRPIVDGEIQEEIELTKEEAEKIFEEFEKETGIDLRVDLSKPGNECAAIAAKIILEILRERSGGS
HHHHHH

Ub-PCNA ligase 32 H6
MKVRIEEVRVKGDSIDAQIEAYNAAREKAKELAKKAKEGEIIIVAAVLETEDGPKAYHLFIVDGKTGKVIDP
GNKQSKDGVAGRPIVDGEVQEEIELSKEEAEEIFKRFKEKTGIDLSVDLTNPGNECAAIAAKIILEILEEQNGG
SHHHHHH

Ub-PCNA ligase 33 H6
MPVRIEEVVVEGDSIDAQIEAYKKALEKAKELAKEAEPGEIIIVNAVLKTPEGDRPYHLLIVDGTTGEVIDPG
NGPVEKGVSGRPVVNGKVQEKIELSYEEAEEVFKRFKERTGLDISVDTKKPGNGCAALAAKIILELLEKQSG
GSHHHHHH

Ub-PCNA ligase 34 H6
MMVRIEEVRVKGDSIDAQIEAYRKARDKAKELAKKAKEGEIIIVAAVLETEQGPEGYHLFIVDGSTGKVIDP
GNKQQKDGVAGRPIVDGEVQEEIELSKEEAEKIFEEFHKETGIDLQVDLDKPGRECAAIAAKIILEILKDQKG
GSHHHHHH

Ub-PCNA ligase 35 H6
MATKVWTQEEADEEIEKYKKRALEALKNGDKALAGIYYGRIAQLYLLLGDEEKAREYIRKAAELIKEAGEL
KRSIEEIEKAFEDGIERGFAVQIDGFDGVLHVAAIFEKDGERRILDPTNVGGDDWEKKKKAIEDAAKKAKEL
EAEGWKPVKPDAEISATTSGNCAAASALAKGAPIVVVDGDRARAEKFADILREIASKHGIDAEISVEIVGGS
HHHHHH

Ub-PCNA ligase 36 H6
MATKVWTQEEAKEEIAKYKKRAEEALKNGDKALAGIYYGRIAALYLLLGDEEKAREYIRKAAELIKEAGT
LSRSIEEIEAAFEKGIEFGFAVYIDGFDGTLHVAAIFEKDGERRILDPTNVGGQDWEKKKNAIEDAAKKAKEL
EAEGWKPVNPDAETLATTAGNCAAASALAKGSPIVVVDSDRERAEKFADILREIARKHGIDAEIRVEIVGGS
HHHHHH

Ub-PCNA ligase 37 H6
MATKVWTQEEADEEIEKYKKQAEEAIKNGDLALAGIYYGRIAALYYLLGDEEKAREYIRKAAELIKEAGEL
KRSIEEIEEAFEAGVEQGFAVYIDGFDGILHVAAVFEKDGKRRILDPTNVGGDDFEAKKKAIEDAAEKAKELE
KEGWKPVNPEAEVTATTSGNCAAASALAKGSPIVVVDDNEERAQRFADILREIAEKYGIKAEIRVEIDGGSH
HHHHH

Ub-PCNA ligase 38 H6
MATKVWTQEEADELIKKYEKKAEEAKKNGDLALAGIYYGRIAALYLLKGDIEKAREYIRKAAELIEEAGTL
KRSIEDIEKAFEQGIETGFAVYIDGFNGTLHVAAIFEKDGKRRILDPTNVGGHDWEKKRAAIEDAAEKAKEL
RAEGWEPVNPESTIETTTAGNCAAASALAKGSPIVVVDSNEERAQKFADILREIAEKHGIKAEISVKIQGGSH
HHHHH

Ub-PCNA ligase 39 H6
MAAKVWTQEEADELIARYKKQAEEAIKNGDLALAGIYYGRIAALYYLKGDEEKAREYIRKAAELIKEAGE
LKRSVEEILKAFEKGREQGFAVYVDGFNGVLHVAAVFEKDGERRILDPTNVGGTDWEAKRAAIEDAARKA
RELRAEGWEPVNPEAEVQTTTDGNCAAASALAKGSPITVVDSNEERANKFADILREIAKKHGIDAKITVKIN
GGSHHHHHH

Ub-PCNA ligase 40 H6
MVVKLITPEEAKEKIEEYEKRAEEAKKNGDKVMAGIFLLYAGLYALAAGDKERARQYAGEAWDLINEQNPI
PWTRDEVAEGIVENAENGVLSVPIGYDGTPHAFAVFKKDGKYVFIDPKNITGGKAEDYKEPIREAVAKAKEL
EKEGWEPIKPTEEIVDTASNACGAAAVAALGGKVVIEDKDIENAEKLKEYIEKAAKEYGVEVEVEVKKKGG
SHHHHHH

Ub-PCNA ligase 41 H6
MAVKLITPEEAEEKIKEYEERAEEALKNGDKVQAGIFKLYAGLYALAAGDEEKAIEYAKEAWKLINEENPDP
WTEDIVASGIVEDAKKGVLLVPIGYDGTQHAFAVFEKDGKYVFIDPKNITGGKAEDYREPIREAVKKAKELE
KEGWKPIPPTEEIVDTADDACGAAAVAALGGKVVIEDEDIENAEKLKEYILEAAAEYGVEVEVEVKQLGGS
HHHHHH

Ub-PCNA ligase 42 H6
MAQGIKEAYINGIAEKGILSEEEAREYVERAWKKGEELRAQGVELVARNGKLSPILAAQAVENIEAGKALTA
GEAALNGEPAGGEIPSELAEEIKKLVSELASPDVYVVPVDFAGEQHLVVVYFEGGKLRAIVDPGNVGVGKN
QGTCYGAALVAAVAVEKVGPAAAAKIGFEVSKEAAERIKKGVPPNVAVAESALKYLRELKGGSHHHHHH

Ub-PCNA ligase 43 H6

MSKAIKDAYIDGIADKGILSEEEAEEYVERWWKKGEELRKQGVELVARNGELSPILAGQAVENIEEGWALTA
GEAAADGHPAGGEIPEELKAELVRLVRELASPDVYVVPVDYAGESHLVVVVYENGKPKAVIDPGNVGVGK
NQGHCYGAALVAAVAAEKVGPREAAKVGFEVSKEAAKRIKKGADPARAVAEAALRELRRLAGGSHHHHH
H

Ub-PCNA ligase 44 H6
MSKAIKDAYIDGIADKGILSKEEAREYVERWWKKGEELRKQGVELVARNGELSPILAGQAVENIEEGKALTA
GEAAEFGAPAGGEIPEELKAELRRLVRELASPDVYVVPVDYAGENHLVVVVYENGKPKAVVDPGNVGVGP
DQGTCYGAALVAAVAAEKVGPRETAKVGFEVSKEAAKKIKKGADPARAVAESALRELRRLAGGSHHHHHH

Ub-PCNA ligase 45 H6
MSKAIKDAYIDGIADKGILSEEEAREYVEAWWKKGEELRKQGVELVARNGELSPILAGQAVENIEEGKALTA
GEAAAEGEPQGGRIPEELKAELVRLVRELASPDVYVVPVDYAGAFHLVVVVYENGKPKAVIDPGNVGVGS
GQGTCYGAALVAAVAAEKVGPREAAKVGFEVSKEAADRIKKGEDPARAVAEAALRELERLAGGSHHHHHH

Ub-PCNA ligase 46 H6
MPQGIINAYIDGIADKGILSEEEAREYVEAAVERGEALRAQGVELVARNGKLSPILAAEAVENIEKGKALTAG
EAAREGRPAGGEIPEELKEELVRLVEELASPDVYVVPVNFAGDRHLVVVVYEGGKPKAVIDPGNVGVGKQQ
GTCYGAALVAAVAAEKVGPENAAKIGFEVSKEAAERIKKGEDPAEAVARAALMYLRELAGGSHHHHHH

Ub-PCNA ligase 47 H6
MPQGIINAYIDGIADKGILSEEEAREYVEAAVERGEALRAQGVELVARNGKLSPILAAQAVENIEEGKALTAG
EAARHGRPAGGEIPEELKEELVRLVEELASPDVYVVPVNFAGERHLVVVVYEGGKPKAVIDPGNVGVGPEQ
GTCYGAALVAAVAAEKVGPENAAKIGFEVSKKAAERIKKGEDPAEAVARAALEYLRELAGGSHHHHHH

Ub-PCNA ligase 48 H6
MPQGIKEAYINGIAEKGILSEEEAREYVERAWKKGEELRAQGVELVARNGKLSPILAAQAVENIEAGKALTA
GEAAKNGRPAQGDIPEELAEEIKKLVSELASPDVYVVPVDFAGDRHLVVVYFENGKLRAIVDPGNVGVGPG
QGTCYGAALVASVAVEKVGPAAAAKIGFEVSKEAAKRIKKGVPPNVAVAESALKYLRELKGGSHHHHHH

### 4 Supplementary references

1. Hibbert, R. G. & Sixma, T. K. Intrinsic Flexibility of Ubiquitin on Proliferating Cell Nuclear Antigen (PCNA) in Translesion Synthesis *. *J. Biol. Chem.* **287**, 39216–39223 (2012).
2. Butcher, J. *et al.* De novo Design of All-atom Biomolecular Interactions with RFdiffusion3. 2025.09.18.676967 Preprint at https://doi.org/10.1101/2025.09.18.676967 (2025).
3. Abramson, J. *et al.* Accurate structure prediction of biomolecular interactions with AlphaFold 3. *Nature* **630**, 493–500 (2024).
4. Qian, T. *et al.* Complex enzyme active site scaffolding by iteratively detuned catalytic guidance. 2026.10.02.756307 Preprint at https://doi.org/10.64898/2026.10.02.756307 (2026).
5. Kabsch, W. XDS. *Acta Crystallogr. D Biol. Crystallogr.* **66**, 125–132 (2010).
6. McCoy, A. J. Solving structures of protein complexes by molecular replacement with Phaser. *Acta Crystallogr. D Biol. Crystallogr.* **63**, 32–41 (2007).
7. Emsley, P., Lohkamp, B., Scott, W. G. & Cowtan, K. Features and development of Coot. *Acta Crystallogr. D Biol. Crystallogr.* **66**, 486–501 (2010).
8. Liebschner, D. *et al.* Macromolecular structure determination using X-rays, neutrons and electrons: recent developments in Phenix. *Acta Crystallogr. Sect. Struct. Biol.* **75**, 861–877 (2019).
9. Williams, C. J. *et al.* MolProbity: More and better reference data for improved all-atom structure validation. *Protein Sci.* **27**, 293–315 (2018).
